# Metabolic adaptation to progressive mitochondrial dysfunction in aging POLG^D257A^ mice

**DOI:** 10.1101/2021.04.29.441996

**Authors:** Esther W. Lim, Michal K. Handzlik, Elijah Trefts, Jivani M. Gengatharan, Reuben J. Shaw, Christian M. Metallo

## Abstract

A decline in mitochondrial function is associated with neurodegeneration and aging. Progressive mitochondrial defects have diverse metabolic consequences that could drive some of the pathophysiological changes that occur with aging. Here, we comprehensively characterized metabolic alterations in Polg^D257A^ mitochondrial DNA mutator mice. Plasma alanine increased dramatically with time, with lactate and other organic acids accumulating to a lesser extent. These changes were reflective of increased glycolysis, rapid gluconeogenesis, and hypoglycemia. Tracing with [^15^N]ammonium revealed impairment of the urea cycle and diversion to purine catabolism. We also measured alterations in the lipidome, observing a general reduction in canonical lipids and the accumulation of 1-deoxysphingolipids, which are synthesized from alanine via promiscuous serine palmitoyltransferase activity. Consistent with 1-deoxysphingolipid’s association with peripheral neuropathy, Polg^D257A^ mice exhibited thermal hypoalgesia. These results highlight the distinct changes that occur in carbon and nitrogen metabolism upon mitochondrial impairment and key metabolic mechanisms which can drive aging-associated neuropathy.

## Introduction

Mitochondria are multi-faceted organelles that are critical for ATP production, maintenance of redox homeostasis, biosynthesis of metabolites, and recycling of metabolic byproducts (Spinelli and Haigis, 2018). A progressive decline in mitochondrial number and function has long been associated with aging and various neurodegenerative diseases. Importantly, the frequency of mitochondrial DNA (mtDNA) mutations and deletions increases with age in humans and animals (Corral-Debrinski et al., 1992; Cortopassi and Arnheim, 1990; Khaidakov et al., 2003; Schwarze et al., 1995), and these aberrations arise through diverse mechanisms including reactive oxygen species (ROS)-mediated oxidative damage (Yakes and Van Houten, 1997), impaired base excision repair (Stuart and Brown, 2006) or decreased polymerase γ (Polg) fidelity (Johnson and Johnson, 2001). More generally, sustained, low-level mutational stress first impacts mitochondrial function, as evidenced by the impacts of low concentration ethidium bromide (Bao et al., 2016; Park et al., 2001) or radiation exposure due to space travel (da Silveira et al., 2020) on mitochondria. Such changes will manifest in diverse metabolic pools as mitochondrial components are across different tissues.

The Polg^D257A^ mtDNA mutator (Polg) mouse is a model of progressive mitochondrial dysfunction and premature aging (Kujoth et al., 2005; Trifunovic et al., 2004). Polg mice carry a missense mutation that significantly reduces 3′-5′ exonuclease activity required for proofreading, leading to a drastic increase in mtDNA mutations and reduced expression of proteins involved in the electron transport chain (ETC) (Dai et al., 2013; Vermulst et al., 2008). These mice display progeroid phenotypes including alopecia, loss of body fat, sarcopenia, kyphosis, anemia, osteoporosis, reduced fertility, cardiomyopathy, and a shortened life span (Kujoth et al., 2005; Trifunovic et al., 2004). Ultimately, this model results in severe mitochondrial functional deficiency which contributes to these diverse phenotypes (Dai et al., 2010; Joseph et al., 2013; Trifunovic et al., 2004). In humans, mutations in the *POLG* gene are associated with a spectrum of mitochondrial diseases including Alpers-Huttenlocher syndrome, progressive external ophthalmoplegia (PEO), myoclonic epilepsy myopathy sensory ataxia (MEMSA), ataxia neuropathy spectrum (ANS), and myocerebrohepatopathy spectrum (MCHS) disorders (Stumpf et al., 2013). The diagnosis of these mitochondrial diseases is often challenging due to the overlapping range of symptoms with multiple organs and a wide range of severity and timing of onset.

Mitochondrial respiration is inextricably tied to central carbon metabolism, and the progressive metabolic compensation that occurs in Polg mice as they accumulate and adapt to mtDNA mutations has not been explored in detail. By observing changes in metabolism that occur in Polg mice over time, we hope to identify key biochemical processes that could be exploited to mitigate mitochondria-associated aging phenotypes. Here, we comprehensively quantified the metabolic alterations and fluxes in Polg mice between the ages of 3-11 months and compared them to age-matched wild type (WT) controls. Among the metabolites that differed between WT and Polg mice, alanine stood out with its levels increasing dramatically over time and as early as 4 months of age, even preceding changes in lactate. Using ^13^C and ^15^N dynamic labeling, we highlight specific changes in central carbon metabolism and nitrogen homeostasis indicative of defects in tricarboxylic acid (TCA) and urea cycles. In addition, we observed significant alterations in lipid metabolism downstream of these effects, including the accumulation of alanine-derived 1-deoxysphingolipids (doxSLs). Pathologically, doxSLs have been associated with Hereditary Sensory and Autonomic Neuropathy type 1 (HSAN1) (Penno et al., 2010), diabetic sensory neuropathy (Fridman et al., 2021; Othman et al., 2015), and paclitaxel-induced neuropathy (Kramer et al., 2015). Consistent with this metabolic alteration, we reveal the development of thermal hypoalgesia in Polg mice, mechanistically linking mitochondrial defects with peripheral neuropathy through amino acid and sphingolipid metabolism. The longitudinal metabolic data from Polg mice can serve as a roadmap for additional drivers of pathophysiologies associated with mitochondrial dysfunction.

## Results

### Impaired bioenergetic metabolism

We monitored Polg mouse body weight monthly from 3-11 months and observed minimal weight gain as they aged (Figure 1A). These results are consistent with previously published Polg cohorts (Kujoth et al., 2005; Trifunovic et al., 2004) but also suggest that significant metabolic alterations are present at an early age. To assess progressive changes in metabolism, we analyzed plasma obtained from the tail vein from WT and Polg mice every month until they were sacrificed at 12 months old.

**Figure 1.**
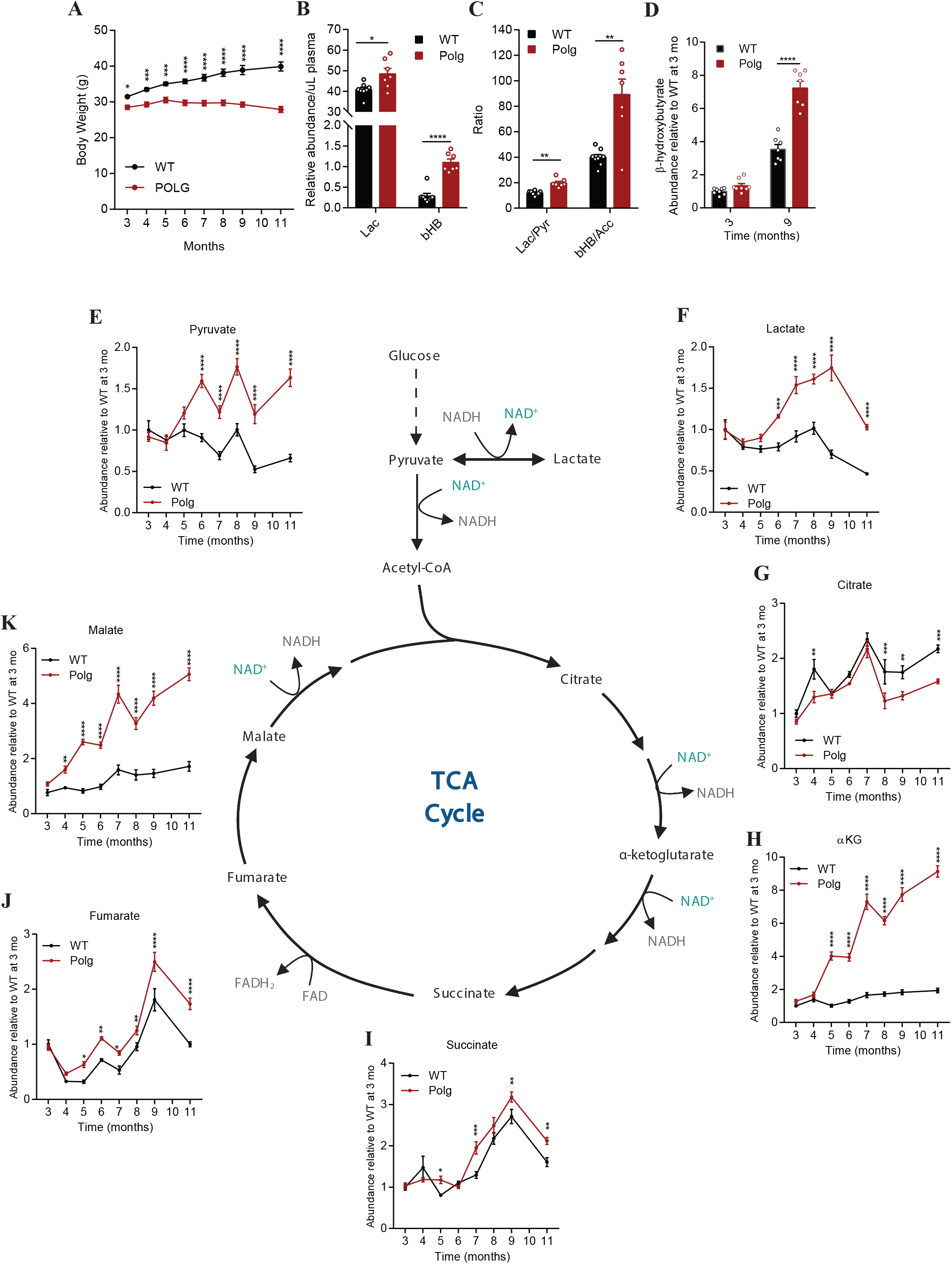
Polg mice have impaired bioenergetic metabolism (A) Body weight of WT and Polg mice over time with images of representative WT and Polg mice. (B) Abundance of lactate (lac) and β-hydroxybutyrate (βHB) relative to internal standard per µL of plasma in WT and Polg mice at 12 months of age. (C) Lactate:pyruvate (lac/pyr) and β-hydroxybutyrate:acetoacetate (βHB/Acc) ratio in WT and Polg plasma at 12 months of age. (D) Abundance of β-hydroxybutyrate relative to internal standard per µL of plasma at 3 and 9 months of age. Abundance of plasma pyruvate (E), lactate (F), citrate (G), α-ketoglutarate (αKG) (H), succinate (I), fumarate (J), malate (K) relative to internal standard, normalized to WT values at 3 months. Two-way ANOVA (A,D,E-K) or two-sided Student’s t-test (B-C) for each comparison with no adjustment for multiple comparisons. Data are mean ± s.e.m. of n=7-8 animals per group. *P < 0.05, **P < 0.01, ***P < 0.001, ****P<0.0001.

Polg mice should accumulate mitochondrial dysfunction as they age, hence we first quantified common mitochondrial dysfunction markers in plasma and saw elevated levels of lactate and β-hydroxybutyrate (Figure 1B). These metabolites are products of key oxidative substrates for tissues throughout the body and reflect alterations in the NAD^+^:NADH ratio. As expected, the lactate:pyruvate and β-hydroxybutyrate:acetoacetate ratios were significantly increased in Polg mice (Figure 1C). Notably, levels of β-hydroxybutyrate were similar at 3 months of age but were significantly elevated in aged Polg mice (Figure 1D). We next quantified organic acids in plasma collected monthly as the mice aged. WT and Polg mice had similar levels of pyruvate and lactate through 5 months of age, but levels of each increased substantially in Polg mice plasma from 6 months onwards (Figure 1E-F).

The tricarboxylic acid (TCA) cycle in the mitochondrial matrix consists of a series of redox reactions to harness energy in the forms of NADH, FADH_2_, and ATP. The electrons in NADH and FADH_2_ are then funneled to the electron transport chain (ETC) through oxidative phosphorylation to generate more ATP. These pathways are coupled in most mitochondria, so accumulation of NADH in the context of reduced mitochondrial function will reduce oxidative TCA metabolism. Consistent with a potential decrease in pyruvate dehydrogenase (PDH) and TCA cycle flux, we observed a reduction in citrate in Polg mice plasma relative to WT mice (Figure 1G). On the other hand, there was a substantial accumulation of TCA cycle intermediates such as α-ketoglutarate, succinate, fumarate, and malate in Polg mice compared to WT (Figure 1H-K). Notably, α-ketoglutarate and malate levels increased strikingly over time in Polg mice, indicating that TCA metabolites are differentially impacted by progressive mtDNA accumulations. Indeed, succinate levels in aged Polg and WT mice differed by only 1.5-fold compared to ∼3-or 5-fold for malate and α-ketoglutarate, respectively (Figure 1 H,I,K). Overall, the accumulation of these metabolites reveals a progressive impairment in bioenergetic metabolism and redox stress that becomes evident as early as 4 months of age for certain metabolites in Polg mice (Figure 1E-K).

### Differential amino acid metabolism

Mitochondria and Complex I of the ETC are intimately linked to amino acid metabolism given their role in α-keto acid oxidation (Figure 2A). Indeed, we observed significant alterations in several amino acids within Polg plasma, with some increasing further over time (Figure 2B). Alanine increased steadily over time in Polg mice but remained unchanged in aging WT mice (Figure 2C). This finding mirrors the observed increase in pyruvate and lactate (Fig 1 E-F) as well as findings in mitochondrial disease patients (Haas et al., 2008). However, the rise in alanine preceded that of other metabolites such as pyruvate and lactate (Figure 1 E-F) by two months, suggesting that alanine levels are extremely responsive to mild reductions in mitochondrial function (Figure 2C). Similar to alanine, proline levels increased steadily over time in Polg mice whereas levels were maintained in WT mice (Figure 2D). Importantly, proline metabolism is involved in a redox coupling and regulatory system where proline and pyrroline-5-carboxylate serve as redox couples that can influence NAD^+^:NADH and NADP^+^:NADPH ratios (Phang, 1985; Tran et al., 2021; Zhu et al., 2021). Proline can also be oxidized indirectly via glutamate and α-ketoglutarate in the TCA cycle, and these catabolic fluxes decrease progressively with Complex I loss.

**Figure 2.**
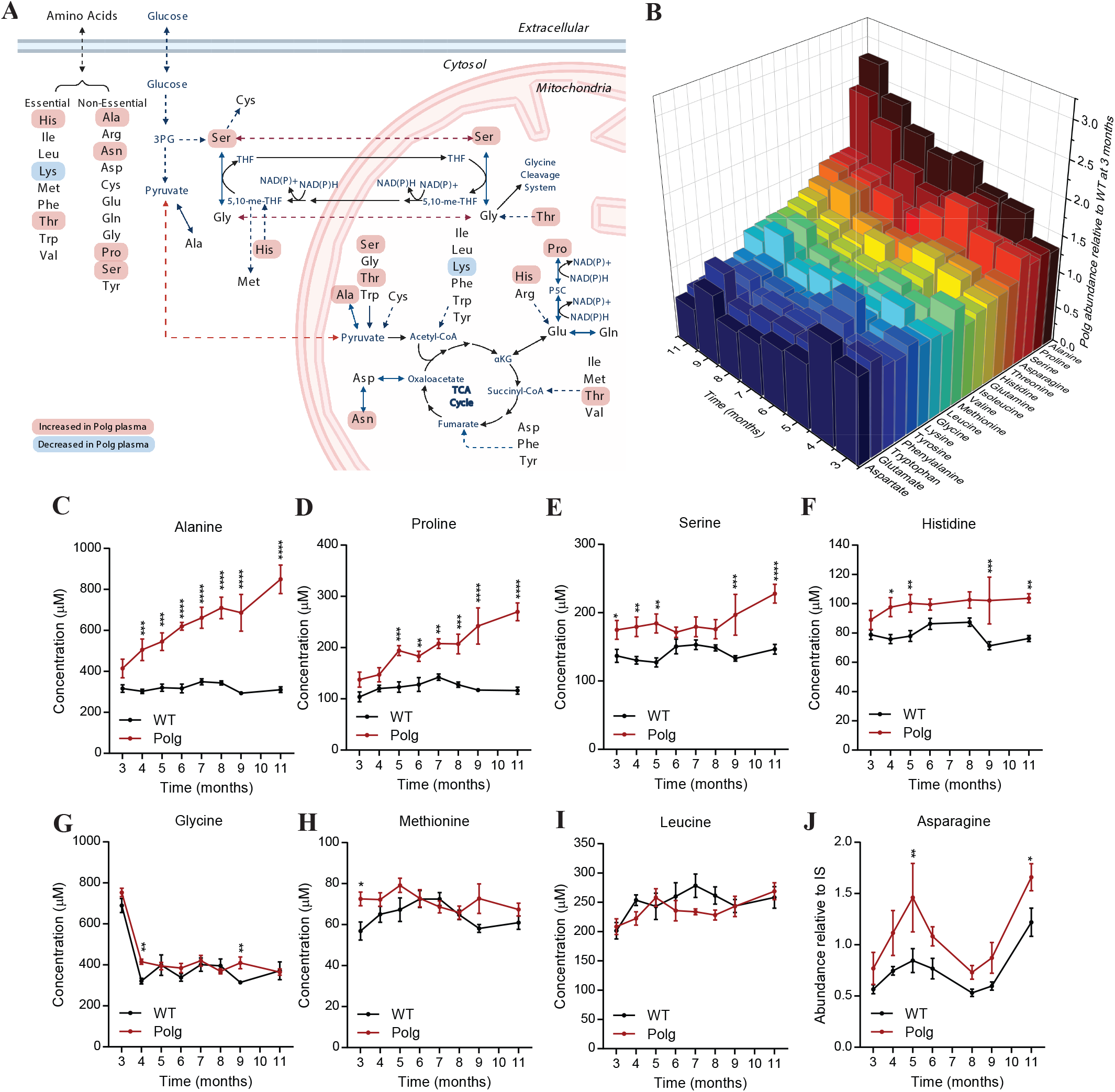
Alanine and proline accumulate in Polg mice plasma with age (A) Simplified pathway map of mitochondrial amino acid metabolism. Created with Biorender.com. (B) 3D plot of average plasma amino acid fold-change in Polg mice relative to WT mice at 3 months. Refer to Supplemental Table 1 for plasma amino acid concentration. Concentration of plasma alanine (C), proline (D), serine (E), histidine (F), glycine (G), methionine (H), and leucine (I) in WT and Polg mice from 3 to 11 months of age. (J) Abundance of plasma asparagine relative to internal standard from 3 to 11 months of age. Two-way ANOVA for each comparison with no adjustment for multiple comparisons. Data are mean ± s.e.m. of n=7-8 animals per group. *P < 0.05, **P < 0.01, ***P < 0.001, ****P<0.0001.

In contrast to the steady increase observed with alanine and proline, serine and histidine abundances remained consistently elevated in Polg plasma from 3 months onward (Figure 2 E-F). On the other hand, glycine levels were largely unchanged between Polg and WT mice (Figure 2G). Given the importance of NAD+ regeneration in *de novo* serine synthesis this result is unexpected (Diehl et al., 2019), and these changes likely reflect reduced catabolic flux through mitochondrial one-carbon metabolism (Bao et al., 2016; Tibbetts and Appling, 2010). Methionine, another amino acid associated with one-carbon metabolism, was generally unchanged between WT and Polg mice (Figure 2H).

Surprisingly, branched-chain amino acids (BCAAs) such as leucine were unchanged between WT and Polg mice (Figure 2I, Supplemental Figure 1A-B). BCAA catabolic flux generates significant NADH and is sensitive to hypoxic stress (Wallace et al., 2018); however, reduced protein synthesis could balance this effect (Lynch et al., 2002). Asparagine is another amino acid noted to increase during mitochondrial stress (Quirós et al., 2017), and levels were generally higher in Polg mouse plasma (Figure 2J). In contrast, the levels of aspartate, an amino acid whose levels are dependent on mitochondrial function (Birsoy et al., 2015; Sullivan et al., 2015) were mostly unchanged between Polg and WT mice (Supplemental Figure 1C). Plasma concentrations of other non-essential amino acids were either unaltered or only significantly different at the last time point (Supplemental Figure 1 D-J).

### Tissue-specific alterations in metabolites

The drastic changes in circulating organic and amino acids led us to measure protein expression of the ETC complexes in the liver, kidney, and skeletal muscle (gastrocnemius) at the termination of the study, as these tissues are major contributors to the inter-organ metabolite pool and exchange. We observed a significant reduction in key subunits of Complex I and IV protein in the liver, kidney, and skeletal muscle (gastrocnemius) (Figure 3A-C). In contrast to the decrease in Complex I and IV, we detected no significant alteration in Complex II and III subunits within these tissues (Figure 3A-C), while Complex V was either increased or unchanged (Figure 3 A-C). Overall, these data corroborate previously published data indicating that the primary impact of dysfunctional Polg encompasses a specific reduction in Complex I and IV (Hauser et al., 2014; McLaughlin et al., 2020; Ross et al., 2019), which are predominantly encoded from the mitochondrial genome. The selective loss of the ETC complexes is consistent with the larger increase in plasma TCA intermediates such as α-ketoglutarate and malate whose catabolism is directly NAD+-dependent (Figure 1 H, K). By contrast, there was a more modest difference in the Complex II-dependent intermediates, succinate and fumarate (Figure 1 I-J).

**Figure 3.**
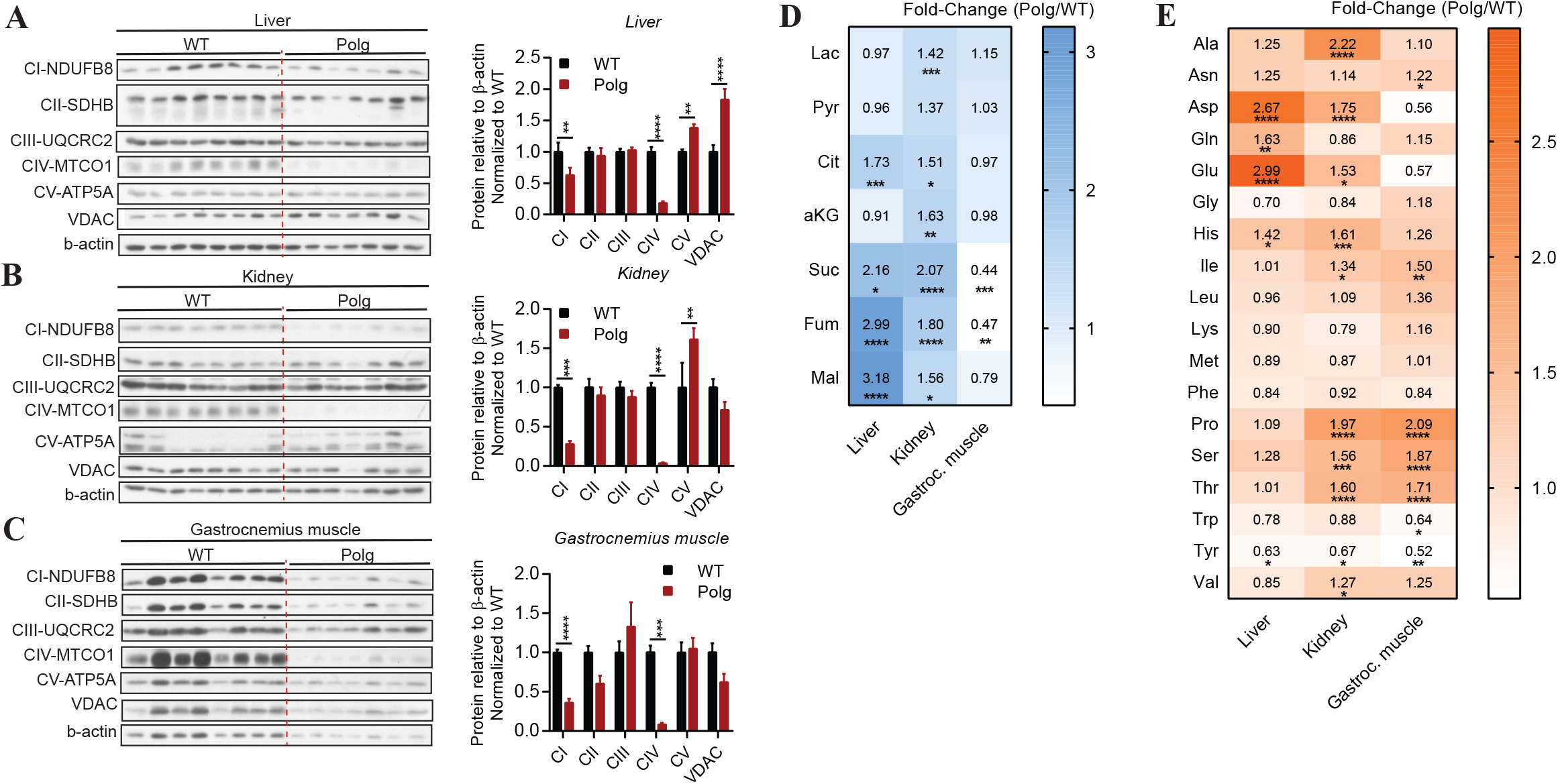
Polg mice exhibit tissue-specific alterations in metabolites Western blot analysis of lysates and protein band quantification (relative density of protein to β-actin) from WT and Polg liver (A), kidney (B), and gastrocnemius muscle (C) targeting subunits of the electron transport chain and voltage-dependent anion-selective channel (VDAC) protein. (D) Heat map representing average fold-change of lactate, pyruvate, and TCA cycle intermediates of Polg liver, kidney, and gastrocnemius muscle relative to WT tissue at 12 months of age. Refer to Supplemental Table 2 for tissue metabolite abundance. (E) Heat map representing average fold-change of amino acids in Polg liver, kidney, and gastrocnemius muscle relative to WT tissue at 12 months of age. Refer to Supplemental Table 3 for tissue amino acid concentration. Two-sided Student’s t-test for each comparison with adjustment of multiple comparisons (Holm-Sidak method) (A-C). Two-way ANOVA with no adjustment for multiple comparisons (D-E). Data are mean ± s.e.m. of n=7-8 animals per group. *P < 0.05, **P < 0.01, ***P < 0.001, ****P<0.0001.

We also quantified metabolite abundances across these tissues. Notably, lactate and pyruvate were significantly increased in the kidney only (Figure 3D). Additionally, there was a significant increase in succinate, fumarate, and malate in Polg liver and kidney, whereas these intermediates were reduced in skeletal muscle (Figure 3D). Amino acids were altered in distinct tissues as well. Aspartate, glutamate, and glutamine accumulated significantly in the liver (Figure 3E). Kidney and skeletal muscle were elevated in histidine, isoleucine, proline, serine, threonine, and valine but showed reduced tyrosine abundance (Figure 3E). Notably, alanine and lactate were dramatically elevated in plasma but showed no significant accumulation in liver or skeletal muscle, which is suggestive of increased Cori and Cahill cycling between these tissues in Polg mice (Supplemental Figure 2A). Their accumulation in the kidney highlights the importance of mitochondrial function in the renal system with respect to bioenergetics, gluconeogenesis, and nitrogen handling.

### Increased glycolysis and gluconeogenesis

Given the profound changes in central carbon metabolism observed in Polg mice, we next sought to assess glucose flux and turnover in these animals. Similar to patients diagnosed with POLG mutations (Montassir et al., 2015; Scalais et al., 2012), Polg mice were hypoglycemic with fasting blood glucose levels that were significantly decreased relative to WT mice (Figure 4A). Additionally, in response to acute glucose challenge, plasma glucose cleared more rapidly in Polg compared to WT mice (Supplemental Figure 3A). Consistent with hypoglycemia, fasting plasma glucagon levels were significantly increased in Polg mice compared to WT mice (Figure 4B). On the other hand, insulin levels were not different between WT and Polg mice (Supplemental Figure 3B).

**Figure 4.**
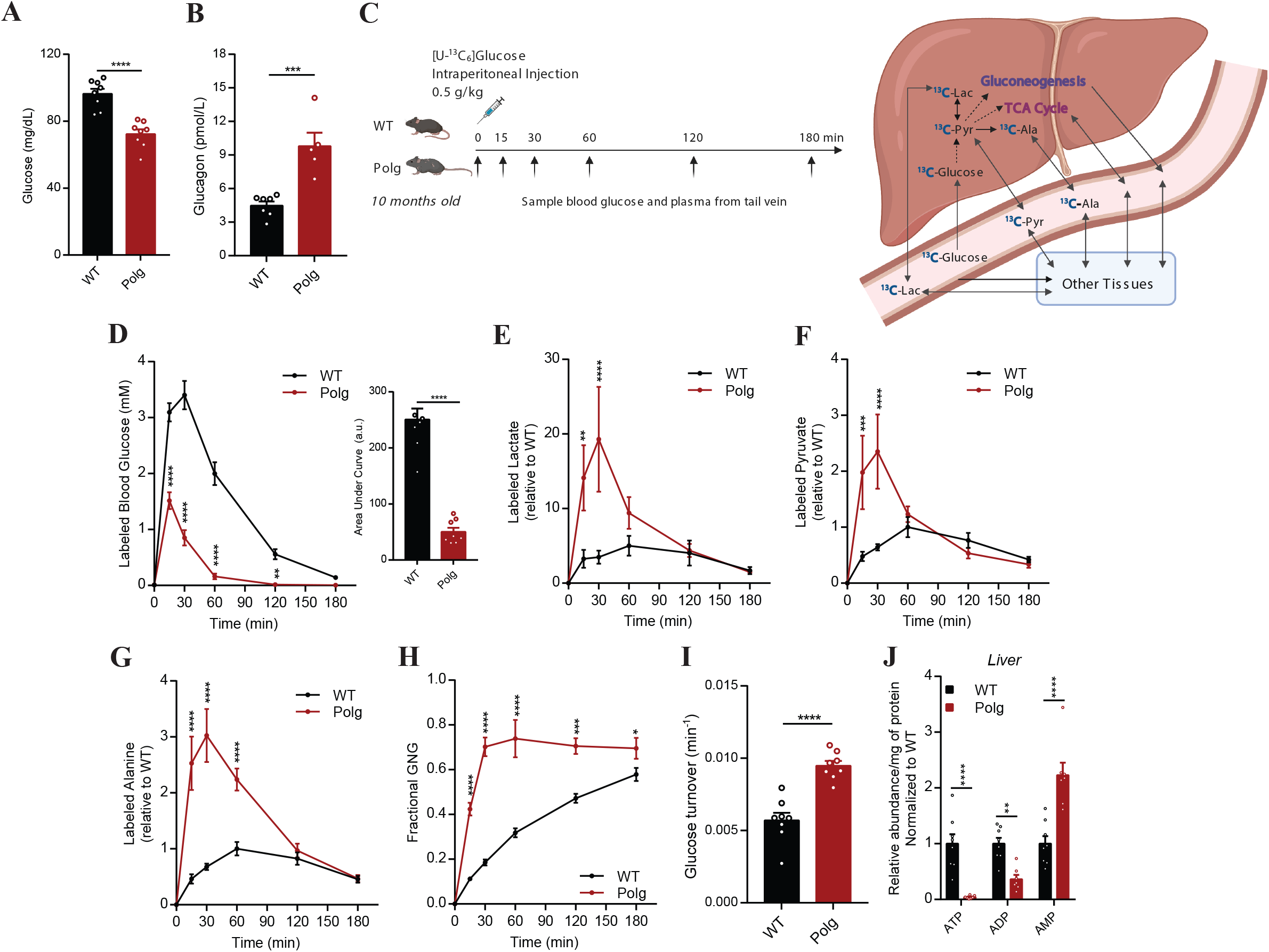
Polg mice exhibit hypoglycemia and increased Cori and Cahill cycling Plasma glucose (A) and glucagon (B) levels in WT and Polg mice measured in fasted state. (C) Schematic of [U-^13^C_6_]glucose tracing experiment and how the ^13^C label is incorporated into newly synthesized pyruvate, alanine, lactate, and glucose. Created with Biorender.com. (D) Levels of ^13^C labeled glucose and quantitation of area under the curve (a.u., arbitrary units). Levels of ^13^C labeled lactate (E), pyruvate (F), and alanine (G) relative to internal standard in plasma over time. Values are normalized to WT, maximum WT value is set to 1. (H) Fractional gluconeogenesis (GNG) contributing to the total glucose flux in plasma. (I) Glucose turnover rate from intraperitoneal bolus injection. (J) Abundance of nucleotide phosphates relative to internal standard in liver at 12 months of age. Abundances were normalized to mg protein per tissue. Refer to Supplemental Table 4 for tissue metabolite abundance. Two-sided Student’s t-test for each comparison with no adjustment for multiple comparisons. Data are mean ± s.e.m. of n=5-8 animals per group. *P < 0.05, **P < 0.01, ***P < 0.001, ****P<0.0001.

To better quantify how glucose was metabolized differently in these mice we administered uniformly-labeled ^13^C [U-^13^C_6_]glucose to mice aged 10 months via intraperitoneal injection and collected plasma from the tail vein thereafter (Figure 4C). Again, we observed that labeled glucose was more quickly metabolized by tissues, as reflected by a rapid decline and lower levels of circulating labeled glucose in Polg mice relative to WT (Figure 4D). Concomitantly, we observed a significant increase in levels of labeled downstream metabolites such as lactate, pyruvate, and alanine in plasma, indicating a rapid uptake and metabolism of glucose in Polg mice (Figure 4 E-G). Indeed, this rapid turnover of glucose was also reflected in the substantial contribution of gluconeogenesis to circulating glucose in Polg mice (Figure 4H). Specifically, we observed a significant increase in the fractional contribution of gluconeogenesis to plasma glucose in Polg mice compared to WT animals (Figure 4H), presumably to maintain euglycemia and circumvent limitations in oxidative phosphorylation. Furthermore, the glucose turnover rate from the intraperitoneal bolus injection was significantly increased in Polg mice relative to WT (Figure 4I). To gain further insight into the energy state in these tissues, we measured levels of nucleotide phosphates in the liver and found that ATP levels were drastically reduced in Polg tissue, even in fed state (Figure 4J). The significant reduction of ATP and increase in AMP in Polg liver is consistent with increased gluconeogenesis to support the demand for Cori and Cahill cycling, further highlighting the importance of hepatic metabolism in regulating the whole-body energy status. Together, these data indicate that Polg mice exhibit an interesting glycolytic/gluconeogenic phenotype, likely to compensate for the lack of mitochondrial respiratory capacity.

### Altered nitrogen metabolism

Mitochondria and TCA intermediates are also associated with the urea cycle, and Polg mice exhibited significant accumulation of several amino acids in plasma, most notably alanine and proline, suggestive of impaired nitrogen homeostasis. Next, we decided to assess how nitrogen handling is altered in Polg mice. *POLG* mutations result in disorders that span a continuum of phenotypes, including liver dysfunction and failure (Harding, 1990; Hikmat et al., 2017; McKiernan et al., 2016). Notably, we observed that Polg mice had significantly reduced urea levels in plasma, suggesting impairment of the urea cycle (Figure 5A).

**Figure 5.**
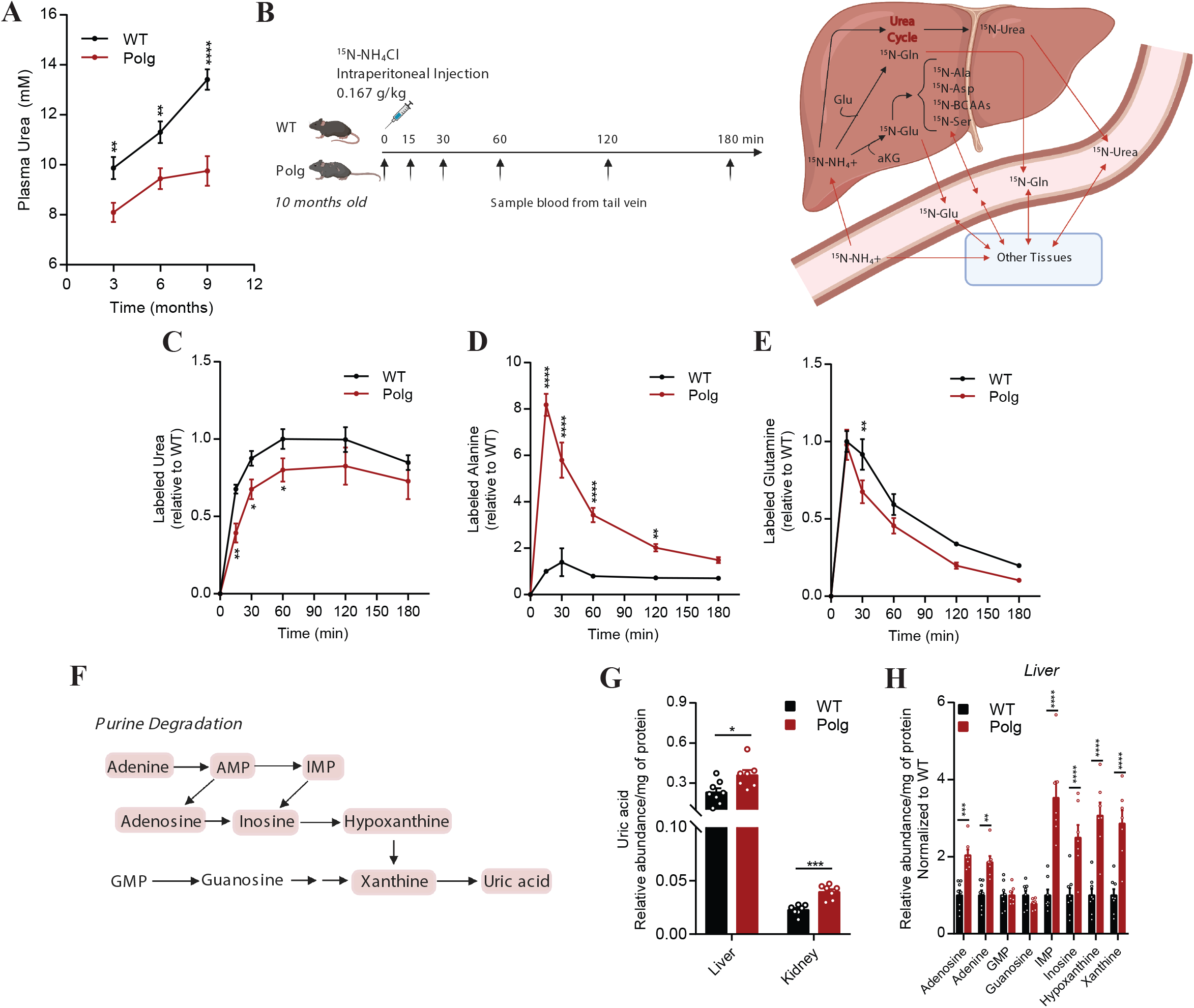
Altered nitrogen handling in Polg mice (A) Concentration of plasma urea in WT and Polg mice at 3, 6, and 9 months of age. (B) Schematic of ^15^N-labeled ammonium chloride (^15^N-NH_4_Cl) tracing experiment and how the ^15^N label is incorporated into newly synthesized urea, glutamine, glutamate, and other metabolites. Created with Biorender.com. Levels of ^15^N-labeled urea (C), alanine (D), and glutamine (E) in plasma relative to internal standard over time. Values are normalized to WT, maximum WT value is set to 1. (F) Schematic of purine degradation pathway. Created with Biorender.com. (G) Abundance of uric acid relative to internal standard in liver and kidney at 12 months of age. Abundances were normalized to mg protein per tissue. (H) Abundance of purine catabolism intermediates relative to internal standard in liver at 12 months of age. Abundances were normalized to mg protein per tissue. Refer to Supplemental Table 5 for tissue metabolite abundance. Two-sided Student’s t-test for each comparison with no adjustment for multiple comparisons. Data are mean ± s.e.m. of n=6-8 animals per group. *P < 0.05, **P < 0.01, ***P < 0.001, ****P<0.0001.

As urea levels were decreased in Polg mice, we wondered if there was compensation in the form of glutamate dehydrogenase using the excess ammonium to form glutamate from α-ketoglutarate (Spinelli et al., 2017). Furthermore, this reaction catalyzed by glutamate dehydrogenase would be driven by the high intracellular NAD(P)H: NAD(P)^+^ ratio. When combined with alanine transaminase, these reactions could also contribute to alanine accumulation in Polg mice. To this end, we administered ^15^N-ammonium chloride (NH_4_Cl) via intraperitoneal injection and collected plasma from the tail over three hours (Figure 5B). We observed a decrease in urea enrichment from ^15^N-NH_4_Cl, consistent with impaired urea cycling (Figure 5C). Although there was no significant difference in ^15^N glutamate abundance (Supplemental Figure 4A), we observed a substantial increase in labeled alanine in the plasma of Polg mice (Figure 5D). We also detected elevated ^15^N enrichment of isoleucine, although the pool size and turnover of alanine are much larger (Supplemental Figure 4B). The considerable labeling on alanine from ^15^N-NH_4_Cl indicates that alanine transaminase (ALT) activity is significantly elevated in Polg mice and matches our findings with [U-^13^C_6_]glucose (Supplemental Figure 4C). The other metabolites were either not significantly different between WT and Polg mice (Supplemental Figure 4 D-E) or had minimal enrichment from ^15^N-NH_4_Cl (Supplemental Figure 4 F-G), indicating that ALT-mediated formation of alanine is a very specific buffer when urea cycle flux decelerates. Glutamine is another important nitrogen pool in plasma, as glutamine synthetase incorporates nitrogen from ammonium into the glutamine pool. Incorporation of ^15^N-NH_4_Cl into plasma glutamine was slightly reduced in Polg mice (Figure 5E). Overall, these results suggest that mitochondrial dysfunction in Polg mice impairs TCA metabolism and nitrogen handling via the urea cycle, which synergize to drive alanine accumulation in Polg mice.

Another mechanism for excreting nitrogenous waste is via purine metabolism through generation of uric acid (Cantor et al., 2017) (Figure 5F). While uric acid was slightly elevated in plasma (Supplemental Figure 4H), we detected a significant increase in uric acid within the liver and kidney of Polg mice (Figure 5G). Furthermore, there was a significant increase in various purine metabolites in Polg liver (Figure 5H) and to a lesser extent in kidney (Supplemental Figure 4I). Interestingly, only adenine and its related metabolites but not guanosine were strongly modulated in Polg liver (Figure 5H). Together, the tracing data suggest that excess nitrogen is buffered and recycled into select amino acids and purines due to reduced urea cycle flux.

### Dysregulated lipid metabolism

Polg mice have reduced body fat compared to WT mice and are resistant to obesity even when fed a high-fat diet (Fox et al., 2012; Wall et al., 2015). Beyond these changes in adiposity, the lipid composition for Polg mice has not been explored in detail. To this end, we quantified various lipid species in our Polg and WT cohorts (Figure 6A). The levels of cholesterol, the most abundant lipid in plasma, were unaltered in Polg plasma (Figure 6B). In contrast, there was a significant reduction in total fatty acids (Figure 6C). Interestingly, the reduction in fatty acids was not uniform across the different species with significant decreases only in myristic acid, palmitic acid, and the essential fatty acid linoleic acid (Figure 6D). These changes reflect potential impacts on fatty acid synthesis as well as absorption from the gut from severe mitochondrial dysfunction. We also observed a significant decrease in triacylglycerols (TAGs) but not diacylglycerols (DAGs) in Polg plasma (Figure 6 E-F, Supplemental Figure 5 A-B). Analysis of total circulating glycerophospholipids revealed a decrease in phosphatidylethanolamine (PE) (Figure 6G, Supplemental Figure 5C) but no change in phosphatidylcholine (PCs) in Polg mice (Figure 6H, Supplemental 5D). Notably, PE synthesis occurs in the mitochondria (Zborowski et al., 1983) in addition to the endoplasmic reticulum (ER) (Kennedy and Weiss, 1956). On the other hand, PC synthesis is dependent on the ER and Golgi membranes in most tissues although the liver and adipocytes are capable of methylating PE to PC in ER-mitochondria membrane domains (Hörl et al., 2011; Vance, 2014; Vance and Ridgway, 1988).

**Figure 6.**
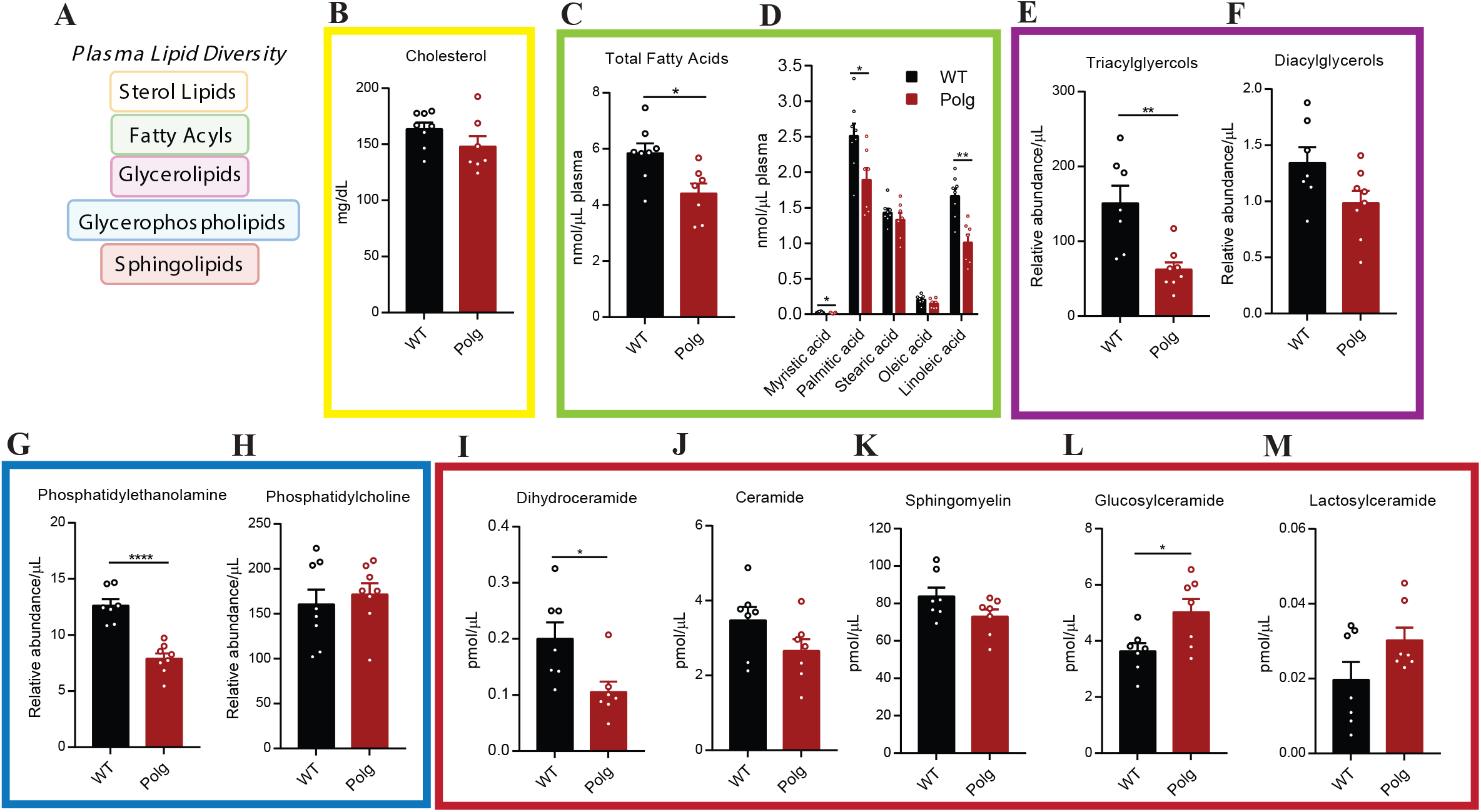
Dysregulated lipid metabolism in Polg mice. (A) Schematic of major plasma lipids. Created with Biorender.com. (B) Concentration of plasma cholesterol in WT and Polg mice. Concentration of total (C) and individual (D) plasma fatty acids in WT and Polg mice. Abundance of plasma triacylglycerols (TAGs) (E) and diacylglycerols (DAGs) (F), phosphatidylethanolamine (G), and phosphatidylcholine (H) relative to internal standard per µL of plasma in WT and Polg mice. Concentration of plasma dihydroceramide (I), ceramide (J), sphingomyelin (K), glucosylceramide (L), and lactosylceramide (M) in WT and Polg mice. Two-sided Student’s t-test for each comparison with no adjustment for multiple comparisons. Data are mean ± s.e.m. of n=7-8 animals per group. *P < 0.05, **P < 0.01, ***P < 0.001, ****P<0.0001.

Sphingolipids are an important class of bioactive lipids that incorporate metabolic signals from fatty acids/acyl-CoAs and amino acids. We observed no differences in the sphingoid bases, sphinganine and sphingosine, between Polg and WT mice (Supplemental Figure 5 E-F). In contrast, there was a significant reduction in plasma dihydroceramides in Polg plasma (Figure 6I, Supplemental Figure 5G), while ceramides remained unchanged (Figure 6J, Supplemental Figure 5H). This decrease in dihydroceramides is suggestive of reduced *de novo* sphingolipid biosynthesis, as dihydroceramides are the direct product of serine palmitoyltransferase (SPT). On the other hand, ceramides are more readily salvaged from sphingomyelin, which was also similarly abundant in Polg and WT mouse plasma (Figure 6K). These data are consistent with plasma cholesterol results, as cholesterol, sphingomyelin, and ceramides are abundant components of lipoproteins. Finally, we observed a slight increase in plasma hexosylceramides but more varied results with less abundant lactosylceramides (Figure 6 L-M, Supplemental Figure 5I). In various sphingolipid classes we observed a specific reduction in species containing acyl chains of C20:0 or longer in Polg mice but the reverse for shorter chain length species (Supplemental Figure 5 G-J). These data suggest that fatty acid elongation or the activity of specific ceramide synthases is compromised in Polg mice.

The sustained increase in plasma alanine in Polg mice led to a progressive increase in the alanine:serine ratio (Figure 7A), which was also elevated in the sciatic nerve (Figure 7B). Alterations in the balance of serine and alanine drive the accumulation of 1-deoxysphingolipids (doxSLs) due to promiscuous activity of SPT (Figure 7C) (Duan and Merrill, 2015; Lone et al., 2019). DoxSLs accumulate in various contexts, including Hereditary Sensory and Autonomic Neuropathy type 1 (HSAN1) (Eichler et al., 2009; Penno et al., 2010), metabolic syndrome (Bertea et al., 2010; Fridman et al., 2021; Mwinyi et al., 2017; Othman et al., 2015), macular telangiectasia (MacTel) (Gantner et al., 2019), dietary serine restriction (Muthusamy et al., 2020), chronic kidney disease (Gui et al., 2021) and aging (Ando et al., 2019). Indeed, 1-deoxysphinganine (doxSA) was significantly elevated in the plasma and sciatic nerve of Polg mice (Figure 7 D-E), and both 1-deoxy dihydroceramide (doxDHCer) and 1-deoxyceramide (doxCer) pools were significantly increased in Polg sciatic nerve (Figure 7F, Supplemental Figure 6 A-B). Importantly, assessment of the canonical sphingolipids including ceramide, sphingomyelin, glucosylceramides, and lactosylceramides revealed no significant differences between WT and Polg mice (Supplemental Figure 6 C-H).

**Figure 7.**
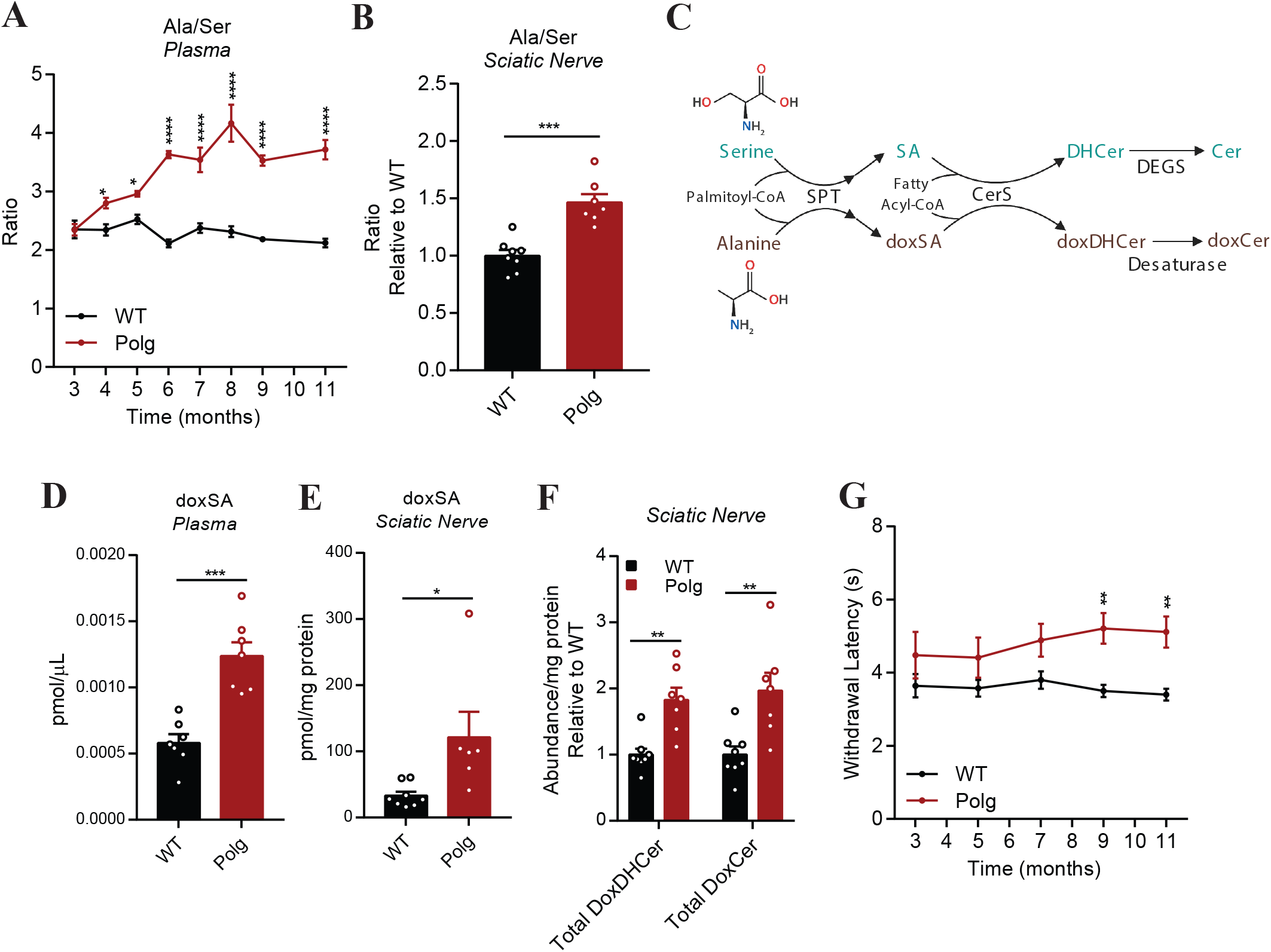
Polg mice accumulate 1-deoxysphingolipids and develop thermal hypoalgesia Ratio of serine:alanine (ser/ala) in plasma from 3 to 11 months of age (A) and sciatic nerve (B). (C) Schematic describing synthesis of canonical sphingolipids from serine (teal) and 1-deoxysphingolipids from alanine (brown) via promiscuous serine palmitoyltransferase (SPT) activity. Ceramide synthase (CerS) produces dihydroceramides (DHCer) or deoxydihydroceramides (doxDHCer). Dihydroceramide desaturase (DEGS) forms ceramides (Cer) and desaturases such as fatty acid desaturase 3 (FADS3) form deoxyceramides (DoxCer) (Karsai et al., 2020). Created with Biorender.com. Concentration of deoxysphinganine (doxSA) in plasma (D) and sciatic nerve (E) in WT and Polg mice. (F) Abundance of sciatic nerve deoxydihydroceramide (doxDHCer) and deoxyceramide (doxCer) relative to internal standard per mg of protein per tissue in WT and Polg mice. (G) Paw heat withdrawal response latency measured every 2 months in WT and Polg mice. Two-way ANOVA (A, G) or two-sided Student’s t-test (B, D-F) for each comparison with no adjustment for multiple comparisons. Data are mean ± s.e.m. of n=7-8 animals per group. *P < 0.05, **P < 0.01, ***P < 0.001, ****P<0.0001.

### Development of thermal hypoalgesia

The elevation in alanine and doxSLs in Polg is intriguing since these lipids have been shown to cause HSAN1 (Penno et al., 2010), an axonal neuropathy marked by thermal hypoalgesia and driven by gain-of-function SPT variants that preferentially use alanine as a substrate. Furthermore, doxSLs have been associated with diabetic peripheral neuropathy (Fridman et al., 2021; Othman et al., 2015) and chemotherapy-induced peripheral neuropathy (Kramer et al., 2015). Peripheral neuropathy is common in mitochondrial diseases although the clinical manifestation spans a wide spectrum (Pareyson et al., 2013). We specifically hypothesized that the increase in circulating alanine drives doxSL synthesis and transport to drive thermal hypoalgesia in Polg mice. Indeed, feeding a serine and glycine-free diet to C57BL/6 mice drives both doxSL accumulation and thermal hypoalgesia measured by hot plate assay (Gantner et al., 2019). Polg mice presented with a slight increase in thermal latency compared to WT mice through 7 months of age, and thermal hypoalgesia became significant at 9 months as determined by Hargreaves test (Figure 7G). On the other hand, we observed no change in nerve conduction velocity or mechanical nociception in Polg mice (Supplemental 6 I-J). This finding is significant given the specificity of the neuropathic phenotype, as doxSLs are genetically linked to HSAN1/thermal hypoalgesia while other sphingolipids (e.g. S1P) have been implicated in other neuropathic phenotypes (Hill et al., 2018). Furthermore, by linking mitochondrial dysfunction to alanine and doxSL accumulation we highlight a distinct mechanism through which aging biochemically drives the accumulation of toxic lipids which cause thermal hypoalgesia (Figure 7G).

## Discussion

Here, we comprehensively characterized the metabolic compensation that occurs during progressive loss of mitochondrial function using the Polg mouse model. We longitudinally profiled metabolites from both Polg and WT mice and observed an accumulation of alanine, pyruvate, and lactate as well as several TCA intermediates and other amino acids. The accumulation of these metabolites is a hallmark of mitochondrial respiratory chain disorders and is consistent with defects associated with loss of Complex I and Complex IV, which are predominantly encoded by the mitochondrial genome. The time course data presented here not only agrees with published clinical data (Clarke et al., 2013; Esteitie et al., 2005; Haas et al., 2008; Legault et al., 2015) but reveals the order and prioritization of the metabolic alterations that occur in response to cumulative mitochondrial stress. The increase in alanine is particularly striking considering how its accumulation preceded lactate and pyruvate.

The hypoglycemia and increased glycolytic flux in Polg mice complement published data indicating that Polg mice rely on glycolysis for energy production (Saleem et al., 2015). Administration of [U-^13^C_6_]glucose allowed us to demonstrate high rates of Cori and Cahill cycling via lactate and alanine, respectively, which also correlated with high fasting glucagon levels compared to WT mice. While glucose was quickly absorbed by tissues and broken down, it was also regenerated rapidly in Polg mice as evidenced by the increase in the fractional contribution of gluconeogenesis to plasma glucose, presumably to achieve euglycemia. The reduction in circulating glucose when combined with increased circulating lactate and alanine further supports the notion of increased Cori and Cahill cycling in Polg mice. The presence of upregulated concomitant glycolytic and gluconeogenic fluxes may further exacerbate substrate and energy deficits stemming from impaired mitochondrial function. Indeed, hepatic ATP levels were significantly diminished at 12 months of age in Polg mice. Notably, patients with POLG mutations have been reported to be hypoglycemic and glucagon insensitive (Montassir et al., 2015; Scalais et al., 2012). The role of endocrine hormones in regulating glucose homeostasis in the context of *POLG*-driven mitochondrial impairment and stress will be insightful to identify therapeutic targets for mitochondrial disease patients.

While alanine is a known gluconeogenic substrate (Felig, 1973), the role of dysfunctional nitrogen metabolism contributing to the alanine pool is underexplored. The impairment in urea cycle as evidenced by ^15^N-NH_4_Cl tracing here provides another mechanism for the accumulation of alanine in Polg mice which may also be relevant for other models of mitochondrial disease. An impaired urea cycle compromises nitrogen disposal, driving the animal to use alternate pathways such as pyruvate transamination to alanine or purine metabolism. Indeed, we observed an increase in uric acid and other purines in Polg mice, which could further exacerbate energetic defects since purine synthesis is an energy-intensive process. These findings highlight additional mechanisms through which mitochondrial defects can influence one-carbon metabolism (Bao et al., 2016; Nikkanen et al., 2016). Notably, a recent mitochondrial genome sequencing study identified several alleles associated with pathogenic gout (Tseng et al., 2018), highlighting the clinical relevance of this pathway alteration. The increase in AMP in Polg mice is intriguing as it suggests potential activation of AMPK, which may be engaged as a mechanism to reverse the bioenergetic stress caused from the cumulative effects of the Polg mutation. As AMPK regulates a diverse set of metabolic pathways to restore ATP levels (Herzig and Shaw, 2018), it would be interesting to understand the role of AMPK in a model of progressive mitochondrial dysfunction such as the Polg mice.

Mitochondrial dysfunction in respiratory chain diseases has been shown to alter a wide range of lipid species that are suggestive of disrupted fatty acid oxidation and/or impaired biosynthesis (Clarke et al., 2013; Legault et al., 2015; Ren et al., 2018). Our survey of the plasma lipidome in Polg mice revealed a reduction in multiple classes of lipids including fatty acids, glycerolipids, glycerophospholipids, and sphingolipids. While the reduction of some (e.g. glycerolipids, glycerophospholipid) is not consistent with previously published data from humans diagnosed with primary mitochondrial disease (Clarke et al., 2013; Legault et al., 2015; Ren et al., 2018), the discrepancy may point toward a different mechanism for the lipid alteration. Importantly, plasma triglyceride content is heavily dependent on diet. In the case of Polg mice, the *POLG* D257A mutation has been shown to limit fat absorption by the intestine leading to attenuated weight gain even on a high-fat diet (Fox et al., 2012). The observed reduction in some lipids could be due to reduced intestinal absorption, which, in turn, could drive shifts in the acyl chain distributions of ceramides, sphingomyelin, and glucosylceramides.

We also observed that the increase in alanine drives the accumulation of doxSLs in Polg mice, which ultimately develop thermal hypoalgesia. While previous work has focused on the role of SPT variants or serine availability in doxSL accumulation, we identify a mitochondria-driven mechanism whereby elevated alanine drives increased levels of doxSLs. In fact, elevated doxDHcer and doxSO have been detected in a small subset of individuals with primary mitochondrial disease and peripheral neuropathy (Ferreira et al., 2018). Others have shown that doxSLs increase in the central nervous system with age (Schwartz et al., 2019). Mechanistically, doxSLs have been shown to disrupt neuronal cytoskeleton structures (Penno et al., 2010), reduce neurite length (Penno et al., 2010), cause mitochondrial fragmentation (Alecu et al., 2017), and induce the collapse of the endoplasmic reticulum membrane (Haribowo et al., 2019). Here we demonstrate that mitochondrial defects can directly drive changes in membrane lipid metabolism. While bioenergetic defects also likely contribute to this peripheral neuropathy phenotype, doxSL accumulation in the context of mitochondrial dysfunction may perturb multiple membrane-associated cellular functions that are deleterious for nervous system function.

Finally, our results highlight the utility of Polg mice in deciphering how to design diets that mitigate phenotypes associated with low-level mitochondrial stress which is experienced during space travel and other environments (da Silveira et al., 2020). By performing longitudinal metabolomic and phenotypic measurements during aging of Polg mice we identified key pathway alterations that lead to neuropathy. The use of stable isotope tracers in such models allowed us to decipher key pathway alterations driving these phenotypes. Since these findings are highly dependent on dietary amino acid composition and nitrogen loading, they can be leveraged to understand how dietary (or pharmacological) interventions could mitigate the deleterious effects of both mild (early) and severe (late) mitochondrial dysfunction. One can also envision using this Polg model and accompanying data to design healthy, sustainable diets for long-term space travel or planetary colonization.

## Acknowledgements

We thank all members of the laboratory of C.M.M. for support and helpful discussions; and Katie E. Frizzi and Nigel A. Calcutt of the University of California, San Diego for providing training and instruments for Hargreaves, von Frey, and nerve conduction velocity analysis and measurement. This work was supported by the NIH (R01CA234245 to C.M.M.; U54CA132379; R35CA220538 and R01DK080425 to R.J.S), a Camille and Henry Dreyfus Teacher-Scholar Award (to C.M.M.), the Lowy Medical Research Foundation (to C.M.M.), and Ruth L. Kirschstein National Research Service Award (NRSA) Individual Postdoctoral Fellowship (F32DK126418 to E.T.).

## Author Contributions

E.W.L., M.K.H., and C.M.M. designed the study. E.W.L., M.K.H., and J.M. G. performed *in vivo* experiments. E.W.L. generated and analyzed metabolomics data. E.T. performed mitochondrial protein isolation and immunoblotting. C.M.M and R.J.S. supervised studies. E.W.L. and C.M.M wrote the manuscript with input from all authors.

## Methods

### Polg^D257A^ mtDNA mutator mouse

Mouse handling and care followed the NIH Guide for Care and Use of Laboratory Animals. The experimental protocols were approved by the UCSD Institutional Animal Care and Use Committee. 12-week old Polg^D257A^ mice were purchased from the Jackson Laboratory. Food and water were provided ad libitum.

### Plasma and tissue sampling

Blood samples were collected in fed state (unless otherwise noted) via tail bleed in EDTA-coated tubes (Sarstedt, Inc.) and spun at 2000 *g* for 5 minutes. The supernatant was transferred to a new eppendorf tube and stored at −80°C until analysis. Tissues were collected in fed state when the mice were 12 months old. Tissues such as the liver, kidney, skeletal muscle, and sciatic nerve were freeze-clamped immediately upon collection using Wollenberger clamps pre-cooled to the temperature of liquid nitrogen, and stored at −80 C until further analysis.

### Polar metabolite analysis (Gas chromatography-mass spectrometry)

Metabolites from 3 µL of plasma were extracted with 250 µL of −20°C methanol, with inclusion of 3 µL 100 µM labeled (^13^C,^15^N) amino acid standard mix (Cambridge Isotope Laboratories, Inc, MSK-CAA-1 or MSK-A2-1.2.). The tubes were vortexed for 10 min and centrifuged at 16,000 x *g* at 4°C for 10 min. The supernatant was collected and dried under vacuum. Derivatization for polar metabolites was performed using a Gerstel MPS with 15μL of 2% (w/v) methoxyamine hydrochloride (Thermo Scientific) in pyridine (incubated for 60 minutes at 45°C) and followed by 15μL N-tertbutyldimethylsilyl-N-methyltrifluoroacetamide (MTBSTFA) with 1% tert-butyldimethylchlorosilane (Regis Technologies) (incubated for 30 minutes at 45°C). Polar derivatives were analyzed by GC-MS using a DB-35MSUI column (30 m x 0.25 mm i.d. x 0.25 m) installed in an Agilent 7890B gas chromatograph (GC) interfaced with an Agilent 5977A mass spectrometer (MS). For metabolite separation, the GC oven was held at 100 °C for 2 min followed by increasing the temperature to 320 °C at a ramp rate of 10 °C/min, and held for 4 min. The eluates from the gas chromatograph were subjected to electron impact ionization. Mass spectrometer scanning was performed over m/z range of 100–650. The MS source and quadrupole were held at 230°C and 150°C, respectively.

Details on specific fragments are provided elsewhere (Cordes and Metallo, 2019). The % isotopologue distribution of each fatty acid and polar metabolite was determined by integration using an in-house MATLAB-based algorithm and corrected for natural abundance using the method described elsewhere (Fernandez et al., 1996).

### Polar metabolite analysis (Liquid chromatography-mass spectrometry)

Details below adapted from MacKay et al. 2015 with some modifications (MacKay et al., 2015). Metabolites from 15 µL of plasma were extracted with 400 µL of −20°C 5:3:2 acetonitrile:methanol:water solution, with inclusion of 10 µL of 100 µg/mL norvaline. The tubes were vortexed for 5 min and centrifuged at 16,000 x g at 4°C for 5 min. The supernatant was then transferred for analysis.

For metabolite extraction from tissue, 10-20 mg of frozen tissue was homogenized with a Precellys Evolution Homogenizer (Bertin Technologies, Inc.) at 6800 rpm in 3 cycles of 20 s, with 30 s pause in between each cycle, in 1 mL −20°C 5:3:2 acetonitrile:methanol:water solution, with inclusion of 10 µL 100 µg/mL labeled (^13^C) nicotinamide adenine dinucleotide (NAD+) (Cambridge Isotope Laboratories, Inc., CLM-10671-PK) and 10 µL of 100 µg/mL norvaline (Sigma-Aldrich, N7502). 20 μL of homogenate was dried under air and redissolved in 20 μL of M-PER buffer (Thermo Fisher Scientific, 78501) for protein estimation with BCA assay. The tubes were centrifuged at 16,000 x *g* at 4°C for 5 min. The supernatant was then transferred for analysis.

Q Exactive orbitrap mass spectrometer with a Vanquish Flex Binary UHPLC system (Thermo Scientific) was used with an iHILIC - (P) Classic, 150 x 2.1mm, 5 µm particle, 200 Å (Hilicon) column at 45°C. 2-5 μL of sample was injected. Chromatography was performed using a gradient of 20 mM ammonium carbonate, adjusted to pH 9.4 with 0.1% ammonium hydroxide (25%) solution (mobile phase A) and 100% acetonitrile (mobile phase B), both at a flow rate of 0.2 mL/min. The LC gradient ran linearly from 80% to 20% B from 2-17 min, then from 20% to 80% B from 17-18 min, then held at 80% B from 18-25 min.

Plasma and liver samples were analyzed in negative mode using spray voltage 3.25 kV. Kidney samples were analyzed in polarity switching mode, with the positive mode using spray voltage 4.25 kV. For both positive and negative mode, auxiliary gas flow 15 arbitrary units and sheath gas flow 25 arbitrary units, with a capillary temperature of 275°C. Full MS (scan range 75-1000 m/z) was used at 70 000 resolution with 1e6 automatic gain control and a maximum injection time of 100 ms. Metabolite retention times were verified using standards and in addition a subset of samples was analyzed in MS2 (top 6) mode to increase confidence in metabolite identification. Mass accuracy was below 5 ppm for all analytes. Data was analyzed using EI-Maven software, and peaks normalized to norvaline or labeled (^13^C) nicotinamide adenine dinucleotide (NAD+) internal standard. Refer to Supplemental Table 6 for ion m/z and retention time.

### Fatty acid and cholesterol analysis

Fatty acid and cholesterol were extracted from 10 µL of plasma with 250 µL methanol, 250 µL saline, and 500 µL chloroform, with inclusion of [^2^H_31_] palmitate (Cambridge Isotopes Laboratory, Inc., DLM-215-1) and [^2^H_7_]cholesterol (Cambridge Isotopes Laboratories, Inc., DLM-3057-PK) as internal standards. The tubes were vortexed for 10 min and centrifuged at 16,000 x *g* at 4°C for 10 min to allow for phase separation. The lower organic layer was collected and dried under air at room temperature. The organic phase was derivatized to form fatty acid methyl esters (FAMEs) via addition of 500 μL 2% H_2_SO_4_ in methanol and incubation at 50°C for 2 hours. FAMEs were extracted via addition of 100 μL saturated salt solution and 500 μL hexane. FAMEs were then analyzed using a Select FAME column (100 m x 0.25mm i.d.) installed in an Agilent 7890A GC interfaced with an Agilent 5975C MS. GC oven was held at 80°C for 1 min after injection, increased to 170°C by 20°C/min, increased to 188°C by 1°C/min, and finally increased to 250°C by 20°C/min and held for 10 minutes.

Post-FAME analysis, the samples were dried under air and converted to cholesterol trimethylsilyl (TMS) derivatives by adding 30 µL N-Methyl-N-(trimethylsilyl) trifluoroacetamide (Regis Technologies) and incubating at 37°C for 30 minutes. Cholesterol derivatives were analyzed by GC-MS using a DB-35MS column (30 m x 0.25 mm i.d. x 0.25 m) installed in an Agilent 7890B gas chromatograph (GC) interfaced with an Agilent 5977B mass spectrometer (MS). The GC oven temperature was held at 150°C for 1 minute, then to 260°C at 20°C/min, then held for 3 minutes, and further increased to 280°C at 10°C/min and held for 15 minutes, and finally ramped up to 325°C for a total run time of approximately 30 minutes. The MS source and quadrupole were held at 230°C and 150°C, respectively, and the detector was operated in scanning mode, recording ion abundance in the range of 100–650 m/z.

### Lipid analysis

Lipids (glycerolipids and glycerophospholipids) were extracted from 10 µL of plasma with 750 µL of ice cold 1:1 methanol/water solution and 500 µL of ice cold chloroform with inclusion of EquiSLASH (Avanti, Croda International Plc, 330731), C15 Glucosyl(β) Ceramide-d7 (d18:1-d7/15:0) (Avanti, Croda International Plc, 330729), and C15 Lactosyl(β) Ceramide-d7 (d18:1-d7/15:0) (Avanti, Croda International Plc, 330727) as internal standards. The tubes were vortexed for 5 min and centrifuged at 16,000g at 4°C for 5 min. The lower organic phase was collected and 2 μL of formic acid was added to the remaining polar phase which was re-extracted with 1 mL of chloroform. Combined organic phases were dried and the pellet was resuspended in 50 μL isopropyl alcohol.

Q Exactive orbitrap mass spectrometer with a Vanquish Flex Binary UHPLC system (Thermo Scientific) was used with an Accucore C30, 150 x 2.1mm, 2.6 µm particle (Thermo) column at 40°C. 5 μL of sample was injected. Chromatography was performed using a gradient of 40:60 v/v water: acetonitrile with 10 mM ammonium formate and 0.1% formic acid (mobile phase A) and 10:90 v/v acetonitrile: 2-propanol with 10 mM ammonium formate and 0.1% formic acid (mobile phase B), both at a flow rate of 0.2 mL/min. The LC gradient ran from 30% to 43% B from 3-8 min, then from 43% to 50% B from 8-9min, then 50-90% B from 9-18min, then 90-99% B from 18-26 min, then held at 99% B from 26-30min, before returning to 30% B in 6 min and held for a further 4 min.

Lipids were analyzed in positive mode using spray voltage 3.2 kV. Sweep gas flow was 1 arbitrary units, auxiliary gas flow 2 arbitrary units and sheath gas flow 40 arbitrary units, with a capillary temperature of 325°C. Full MS (scan range 200-2000 m/z) was used at 70 000 resolution with 1e6 automatic gain control and a maximum injection time of 100 ms. Data dependent MS2 (Top 6) mode at 17 500 resolution with automatic gain control set at 1e5 with a maximum injection time of 50 ms was used. Data was analyzed using EI-Maven software, and peaks normalized to Avanti EquiSPLASH internal standard. Refer to Supplemental Table 7 for ion transitions of lipid species.

For targeted sphingolipid analysis, 50 µL of plasma was extracted with 500 µL of −20°C methanol, 400 µL saline and 100 µL of water spiked with deuterated internal standards (100 picomoles of D7-sphingosine (Avanti, Croda International Plc, 860657), 20 picomoles of D7-sphinganine (Avanti, Croda International Plc, 860658), 2 picomoles of D3-deoxysphinganine (Avanti, Croda International Plc, 860474), 200 picomoles of C15 ceramide-d7 (d18:1-d7/15:0) (Avanti, Croda International Plc, 860681), 100 picomoles of C13-dihydroceramide-d7 (d18:0-d7/13:0) (Avanti, Croda International Plc, 330726), 10 picomoles of glucosylsphingosine (d18:1-d7), (Avanti, Croda International Plc, 860695), 100 picomoles of glucosylceramide (d18:1-d7/15:0), (Avanti, Croda International Plc, 330729), 100 picomoles of lactosylceramide (d18:1-d7/15:0), (Avanti, Croda International Plc, 330727), 200 picomoles of sphingomyelin (d18:1/18:1-d9), (Avanti, Croda International Plc, 791649)). 1 mL of chloroform was then added to the tubes. The tubes were vortexed for 5 min and centrifuged at 16,000 x *g* at 4°C for 5 min. The lower organic phase was collected and 2 μL of formic acid was added to the remaining polar phase which was re-extracted with 1 mL of chloroform. Combined organic phases were dried and were resuspended in 100 μL of 0.2% formic acid and 1 mM ammonium formate in methanol. Next, the tubes were sonicated in a bath sonicator for 10 min and spun at 16,000 x *g* for 10 min at 4 °C.

For sphingolipid extraction from sciatic nerve, frozen tissue was homogenized with a ball mill (Retsch Mixer Mill MM 400) at 30 Hz for 3 minutes in 500 μL −20°C methanol, 400 μL of ice-cold saline, and 100 μL of ice-cold MiliQ water, spiked with deuterated internal standards as described earlier. The mixture was then transferred into a 2 mL Eppendorf tube containing 1 mL of chloroform. The tubes were vortexed for 5 min and centrifuged at 16,000g at 4°C for 5 min.

The lower organic phase was collected and 2 μL of formic acid was added to the remaining polar phase which was re-extracted with 1 mL of chloroform. Combined organic phases were dried and were resuspended in 50 μL of 0.2% formic acid and 1 mM ammonium formate in methanol. Finally, the tubes were sonicated in a bath sonicator for 10 min and spun at 16,000 x *g* for 10 min at 4 °C.

Ceramide species in the supernatant was then quantified via liquid chromatography–mass spectrometry (Agilent 6460 QQQ, MassHunter LC/MS Acquisition (v.B.08.02)). Ceramides were separated on a C8 column (Spectra 3 μm C8SR 150 × 3 mm inner diameter, Peeke Scientific) as previously described (Bielawski et al., 2009). Mobile phase A was composed of 100% HPLC-grade water containing 2 mM ammonium formate and 0.2% formic acid and mobile phase B consisted of 100% methanol containing 0.2% formic acid and 1 mM ammonium formate. The gradient elution program consisted of the following profile: 0 min, 82% B; 3 min, 82% B; 4 min, 90% B, 18 min, 99% B; 25 min, 99%, 27 min, 82% B, 30 min, 82% B. Column re-equilibration followed each sample and lasted 10 min. The capillary voltage was set to 3.5 kV, the drying gas temperature was 350 °C, the drying gas flow rate was 10 l/min and the nebulizer pressure was 60 psi. Ceramide species were analyzed by selective reaction monitoring of the transition from precursor to product ions at associated optimized collision energies and fragmentor voltages are provided elsewhere (Muthusamy et al., 2020). Ceramide and dihydroceramide species were then quantified from spiked internal standards.

### Stable-isotope tracing

For stable-isotope tracing experiments, food was removed from cages at 5:00pm the night before the study. Blood was collected via tail bleed at baseline, 15, 30, 60, 120, and 180 minutes post intraperitoneal (I.P.) injection of stable-isotope tracer. Mice remained in their original cage with original bedding for the duration of the experiment.

Uniformly labeled ^13^C-glucose ([U-^13^C_6_]glucose) (Cambridge Isotopes Laboratories, Inc., CLM-1396-PK) was prepared as a 50 g/L solution in sterile phosphate buffered saline (PBS). On the day of experiment, mice were weighed and the appropriate volume of [U-^13^C_6_]glucose solution was administered via I.P. injection at 0.5 g/kg. 15N-labeled ammonium chloride (^15^N-NH_4_Cl) (Cambridge Isotopes Laboratories, Inc., NLM-467-PK) was prepared as a 16.7 g/L solution in sterile PBS. On the day of experiment, mice were weighed and the appropriate volume of ^15^N-NH_4_Cl was administered via I.P. injection at 0.167 g/kg.

Fractional gluconeogenesis was determined as described elsewhere (Kelleher, 1999):

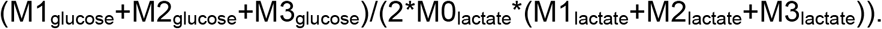

Using the single-pool model, glucose turnover rate was calculated as described elsewhere (Wolfe and Chinkes, 2004): *Metabolite turnover*,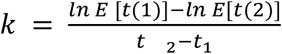with E being the plasma enrichment from labeled glucose at t_1_ = 15 min and t_2_ = 180 min.

### Insulin and glucagon measurement

Insulin and glucagon levels were determined in fasted plasma using commercially available ELISA kits from Mercodia, Inc. (Insulin ELISA (10-1247-01)) Glucagon ELISA (10-1281-01)). Concentrations were measured according to manufacturer’s instructions.

### Protein extraction and immunoblotting

Tissue lysates (Liver, whole kidney, and skeletal muscle) were generated from 10-20 mg pieces of frozen tissue homogenized in CST lysis buffer (20mM Tris-HCl pH 7.5, 150 mM NaCl, 1 mM EDTA, 1 mM EGTA, 50 mM sodium fluoride, 1% Triton, 2.5 mM sodium pyrophosphate, 1 mM beta-glycerophosphate, 1 mM Na3VO4, 10 nM Calyculin A) supplemented with protease inhibitors (Roche, 11836170001). Tissues were mechanically homogenized for 30 seconds and left on ice for 10 minutes. Lysates were then centrifuged at 16,000 x g for 15 min at 4°C and supernatant was transferred to a second microtube. Protein concentration was determined using Pierce™ BCA protein assay kit (Thermo Scientific, 23225). SDS-PAGE running samples were prepared and incubated either at room temperature, for resolution of mitochondrial oxphos complex proteins, or at 95°C, for resolution of β-actin and voltage dependent anion channel (VDAC). Samples were then resolved on 12% SDS-PAGE gels and transferred to PVDF membranes for immunoblotting. Total OXPHOS Rodent WB Antibody Cocktail (abcam, ab110413), β-ACTIN (Sigma, A5441), and VDAC (CST, 4661) were used to probe their respective targets. Horseradish peroxidase-conjugated secondary antibodies were anti-rabbit IgG (Millipore, AP132P) and anti-mouse IgG (Millipore, AP124P).

### Hargreaves test

Hind limb withdrawal reaction to escalating heat stimuli was measured in conscious, unrestrained mice, as described elsewhere (Jolivalt et al., 2016). Briefly, mice were allowed to acclimate to an observation chamber for 30 minutes prior to testing. Paw response latency to surface heat escalating at a rate of 1°C/s from a starting surface temperature of 30°C was measured using a Hargreaves apparatus. Triplicate measurements were made at 5 minute intervals, and the mean was used to average paw heat response latency.

### von Frey test

Hind limb withdrawal reaction to light touch was measured in conscious, unrestrained mice, as described elsewhere (Jolivalt et al., 2016). Briefly, mice were allowed to acclimate to an observation chamber for 30 minutes prior to testing. Paw withdrawal to pressure applied at the plantar surface was measured using von Frey filaments applied sequentially in an up-down protocol as originally described for rats (Chaplan et al., 1994) and subsequently modified for use in mice (Jolivalt et al., 2016). Filaments were applied for 1 s at 1 s intervals with 5 min break between each set of stimulations, with 5 stimulations per filament. Response frequency for each filament was recorded and 50% threshold was calculated using Hill equation (Origin 2017, OriginLab).

### Nerve conduction velocity measurement

Conduction velocity of large myelinated motor fibers was measured in the sciatic nerve as described elsewhere (Jolivalt et al., 2016). Briefly, mice were anesthetized with isoflurane, core and nerve temperatures were held at 37°C using heating pads and lamps, and fine needle stimulating electrodes were placed at the sciatic notch and Achilles tendon. Recording electrodes were inserted into the interosseus muscles of the ipsilateral paw. The sciatic nerve was stimulated using a PowerLab 4/30 to achieve maximal M wave amplitude (0.2–1.0 mV, 0.05-millisecond square waves), and the resulting electromyogram was stored to a computer running LabChart Pro (AD Instruments). Nerve conduction velocity (NCV) was calculated as the distance between stimulation sites divided by the latency of M wave peaks produced by stimulation at the two sites. The median of three separate measurements was used to represent NCV for each animal.

### Quantification and statistical analysis

All results are depicted as mean ± s.e.m. of at least five biological replicates, as indicated in figure legends, using GraphPad Prism (v.8.0.1). P values were calculated using two-way ANOVA or two-sided Student’s t-test for each comparison either with or without adjustment for multiple comparisons. Data are mean ± s.e.m. of n=5-8 animals per group. *P < 0.05, **P < 0.01, ***P < 0.001, ****P<0.0001.

## Supplemental Figure Legends

Supplementary Figure 1. Amino acid profiles in Polg mice

Concentration of plasma isoleucine (A), valine (B), aspartate (C), threonine (D), lysine (E), phenylalanine (F), tyrosine (G), and glutamate (H) in WT and Polg mice from 3 to 11 months of age.

Abundance of plasma tryptophan (I) and glutamine (J) relative to internal standard from 3 to 11 months of age.

Two-way ANOVA for each comparison with no adjustment for multiple comparisons. Data are mean ± s.e.m. of n=7-8 animals per group. *P < 0.05, **P < 0.01, ***P < 0.001, ****P<0.0001.

Supplementary Figure 2. Metabolite cycling in Polg mice

(A) Schematic of Cori and Cahill cycling. Created with Biorender.com.

Supplementary Figure 3. Differential glucose metabolism in Polg mice

(A) Concentration of blood glucose over time upon bolus administration of glucose via intraperitoneal injection and quantitation of area under the curve (a.u., arbitrary units).

(B) Concentration of fasting plasma insulin in WT and Polg mice.

Two-sided Student’s t-test for each comparison with no adjustment for multiple comparisons. Data are mean ± s.e.m. of n=8 animals per group. *P < 0.05, **P < 0.01, ***P < 0.001, ****P<0.0001.

Supplementary Figure 4. Nitrogen metabolism and ^15^N-NH_4_Cl tracing in Polg mice.

Levels of ^15^N-labeled glutamate (A) and isoleucine (B) in plasma relative to internal standard over time. Values are normalized to WT, maximum WT value is set to 1.

(C) Fraction of ^15^N labeling on plasma alanine over duration of experiment.

Levels of ^15^N-labeled aspartate (D) and valine (E) in plasma relative to internal standard over time. Values are normalized to WT, maximum WT value is set to 1.

M0 enrichment of ^15^N label on leucine (F) and serine (G) over time. 100% M0 labeling indicates no incorporation of ^15^N label onto metabolite.

(H) Abundance of uric acid relative to internal standard in plasma at 12 months of age.

(I) Abundance of purine catabolism intermediates relative to internal standard in the kidney at 12 months of age. Abundances were normalized to mg protein per tissue.

Two-sided Student’s t-test for each comparison with no adjustment for multiple comparisons. Data are mean ± s.e.m. of n=7-8 animals per group. *P < 0.05, **P < 0.01, ***P < 0.001, ****P<0.0001.

Supplementary Figure 5. Lipid abundances in WT and Polg mice

Abundances of individual plasma triacylglycerols (TAGs) (A), diacylglycerols (DAGs) (B), phosphatidylethanolamine (C), and phosphatidylcholine (D) relative to internal standard per µL of plasma in WT and Polg mice.

Concentrations of plasma sphinganine (E) and sphingosine (F) in WT and Polg mice. Concentration of individual dihydroceramides (DHCer) (G), ceramides (Cer) (H), sphingomyelin (SM) (I), and glucosylceramides (GlucCer) (J) in WT and Polg mice.

Two-sided Student’s t-test for each comparison with no adjustment for multiple comparisons. Data are mean ± s.e.m. of n=7-8 animals per group. *P < 0.05, **P < 0.01, ***P < 0.001, ****P<0.0001.

Supplementary Figure 6. Sciatic nerve sphingolipids, nerve conduction velocity and mechanical nociception in Polg mice

Abundances of individual sciatic nerve deoxydihydroceramides (doxDHCer) (A) and deoxyceramides (doxCer) (B) in WT and Polg mice.

Concentrations of total sciatic nerve sphinganine (SA) (C), dihydroceramide (DHCer) (D), ceramide (Cer) (E), sphingomyelin (SM) (F), glucosylceramides (GlucCer) (G), and lactosylceramides (LactCer) (H) in WT and Polg mice.

(I) Motor nerve conduction velocity measured every 2 months in WT and Polg mice.

(J) Paw response to von Frey filaments measured at 3 and 9 months in WT and Polg mice.

Two-sided Student’s t-test for each comparison with no adjustment for multiple comparisons. Data are mean ± s.e.m. of n=7-8 animals per group. *P < 0.05, **P < 0.01, ***P < 0.001, ****P<0.0001.

## Supplemental Table Titles

Supplemental Table 1. Plasma Amino Acid Concentration and fold-change (Polg/WT) as shown in Figure 2.

Supplemental Table 2. Tissue metabolite abundance and fold-change (Polg/WT) as shown in Figure 3D.

Supplemental Table 3. Tissue amino acid concentration and fold-change (Polg/WT) as shown in Figure 3E.

Supplemental Table 4. Tissue nucleotide phosphate abundance and fold-change (Polg/WT) as shown in Figure 3F.

Supplemental Table 5. Tissue purine catabolism intermediate abundance and fold-change (Polg/WT) as shown in Figure 5H and Supplemental Figure 4I.

Supplemental Table 6. m/z and retention time for LC/MS (iHILIC) analysis

Supplemental Table 7. Ion transitions for LC/MS (Accucore C30) analysis

