## Supplemental Figures & Tables for "Metabolic adaptation to progressive mitochondrial dysfunction in aging POLG^D257A^ mice"

Supplemental Figure 1. Amino acid profiles in Polg mice

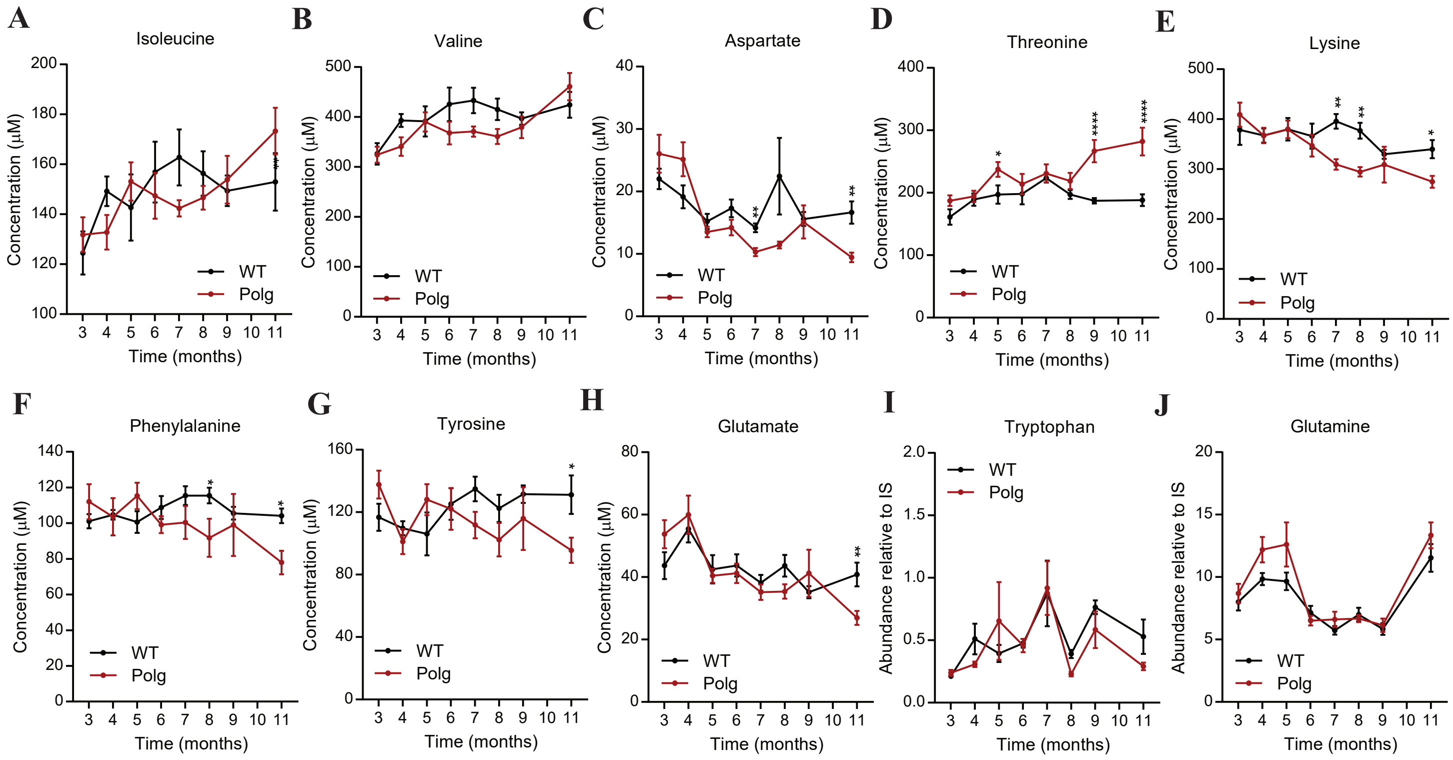

Supplemental Figure 2. Metabolite Cycling in Polg mice

**A**

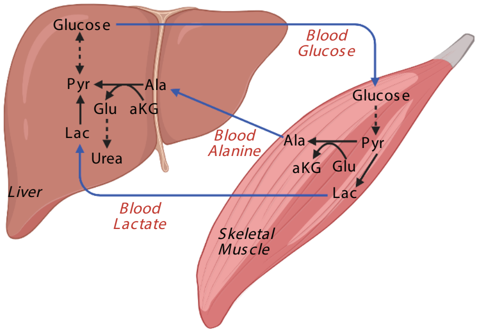

Supplemental Figure 3. Differential glucose metabolism in Polg mice

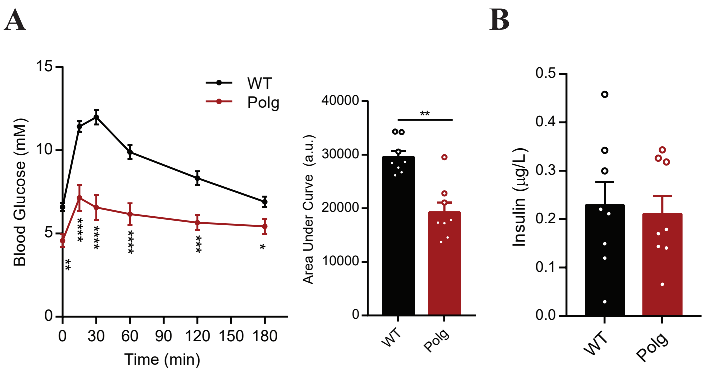

### Supplemental Figure 4. Nitrogen metabolism and 15N-NH4Cl tracing in Polg mice

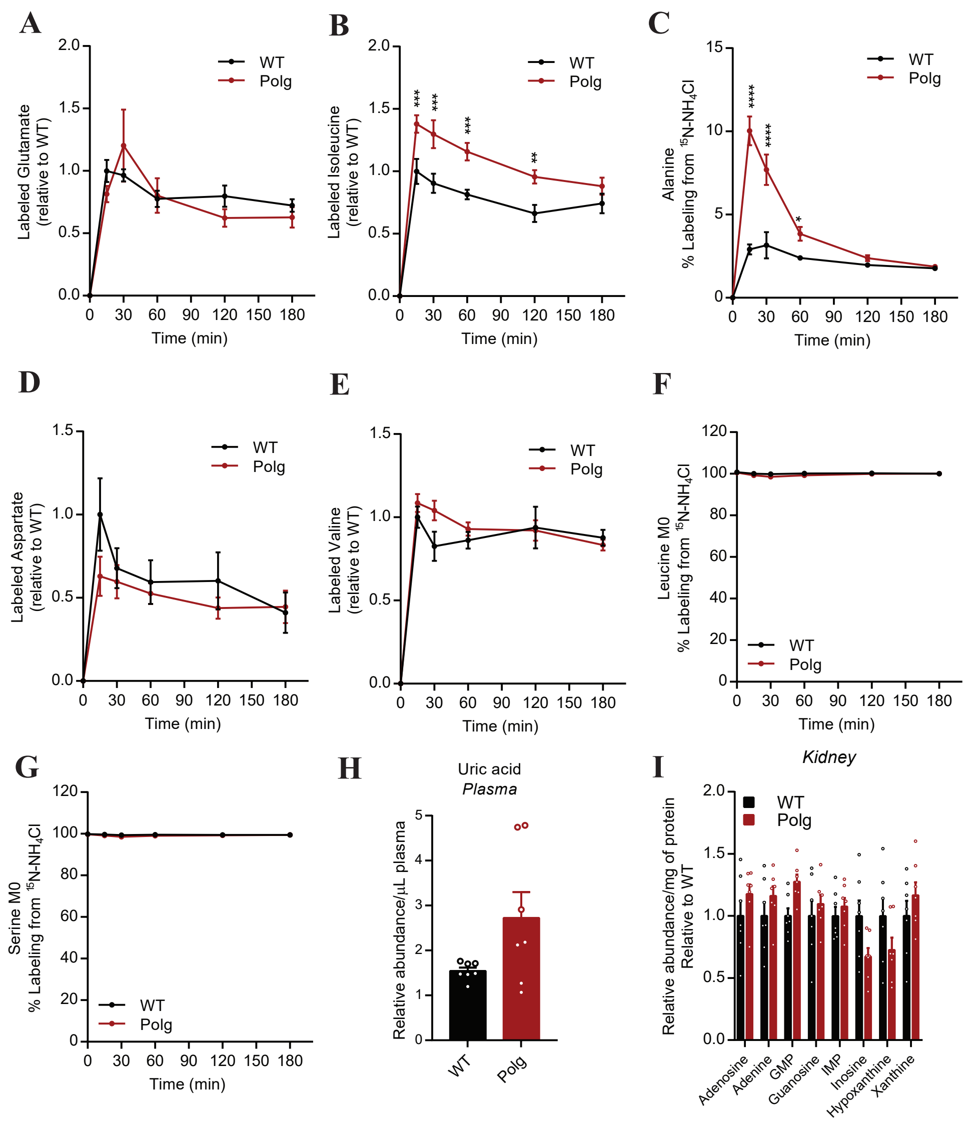

Supplemental Figure 5. Lipid abundances in WT and Polg mice

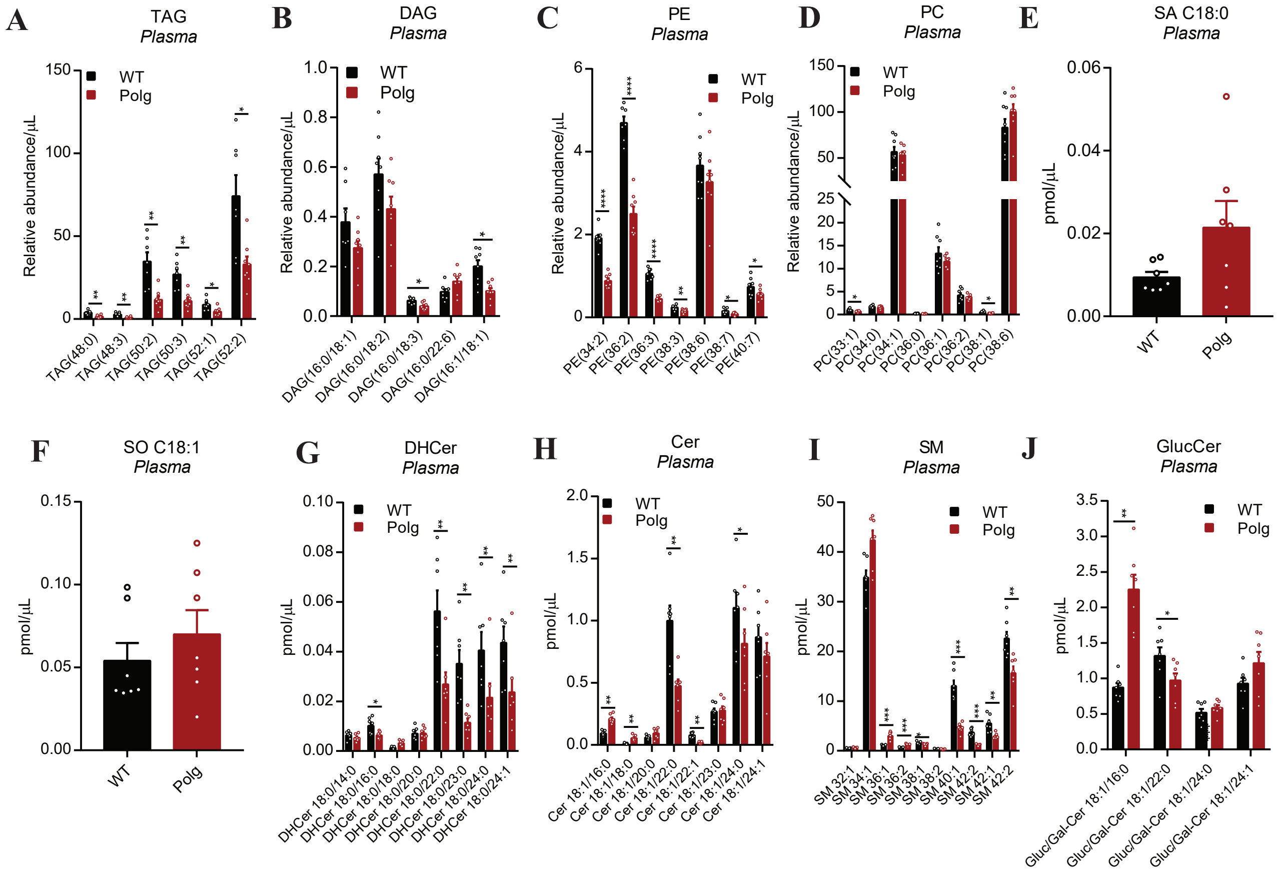

Supplemental Figure 6. Sciatic nerve sphingolipids, nerve conduction velocity and mechanical nociception in Polg mice

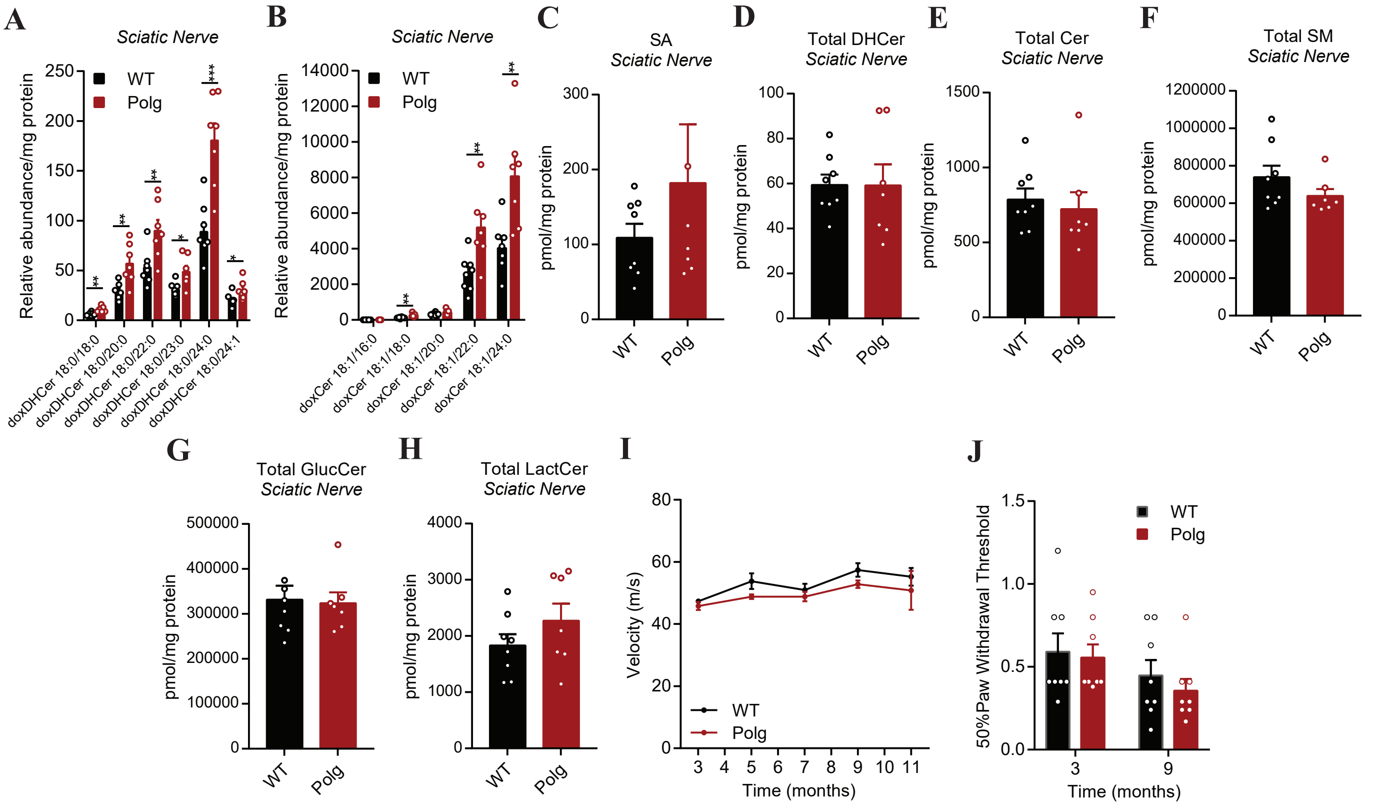

**Supplemental Table 1. Plasma Amino Acid Concentration and fold-change (Polg/WT) as shown in Figure 2**

**ALANINE**

Concentration (µM)

| Time (months) | WT | | | | | | | | Polg | | | | | | | |
|---|---|---|---|---|---|---|---|---|---|---|---|---|---|---|---|---|
| 3 | 331.81 | 360.71 | 330.77 | 324.79 | 385.05 | 314.96 | 244.01 | 238.53 | 404.90 | 447.53 | 701.28 | 404.81 | 293.31 | 397.30 | 369.68 | 294.60 |
| 4 | 309.90 | 354.00 | 285.20 | 320.30 | 269.80 | 268.30 | 290.30 | 324.50 | 437.20 | 503.50 | 859.50 | 520.80 | 425.70 | 443.20 | 404.60 | 450.00 |
| 5 | 272.70 | 400.40 | 358.70 | 276.00 | 266.10 | 354.40 | 294.90 | 341.50 | 447.60 | 472.50 | 826.90 | 518.90 | 556.60 | 453.00 | 554.20 | 532.30 |
| 6 | 285.90 | 393.80 | 384.00 | 210.50 | 314.70 | 352.60 | 285.00 | 307.90 | 591.90 | 619.40 | 704.00 | 541.70 | 567.60 | 619.20 | 670.70 | 642.80 |
| 7 | 404.00 | 353.90 | 397.90 | 305.30 | 299.70 | 351.40 | 308.20 | 373.10 | 566.90 | 553.30 | 964.00 | 643.80 | 595.00 | 517.50 | 758.10 | 694.70 |
| 8 | 361.00 | 367.60 | 331.60 | 383.00 | 299.50 | 341.40 | 314.40 | 353.90 | 726.90 | 518.10 | 1012.50 | 737.80 | 643.20 | 691.10 | 765.90 | 580.90 |
| 9 | 274.80 | 334.70 | 295.20 | 304.60 | 310.20 | 288.20 | 250.20 | 290.60 | 558.70 | 464.00 | 1261.60 | 515.20 | 659.80 | 679.20 | 779.40 | 569.20 |
| 11 | 350.40 | 312.20 | 351.30 | 361.10 | 273.50 | 301.50 | 270.50 | 266.00 | 705.10 | 600.90 | 1015.90 | 827.20 | 1225.00 | 834.00 | 879.80 | 705.50 |

*Values when normalized to WT 3 months (i.e. set WT 3 month vales to 1)*

| Time (months) | WT | | | | | | | | Polg | | | | | | | |
|---|---|---|---|---|---|---|---|---|---|---|---|---|---|---|---|---|
| 3 | 1.05 | 1.14 | 1.05 | 1.03 | 1.22 | 1.00 | 0.77 | 0.75 | 1.28 | 1.41 | 2.22 | 1.28 | 0.93 | 1.26 | 1.17 | 0.93 |
| 4 | 0.98 | 1.12 | 0.90 | 1.01 | 0.85 | 0.85 | 0.92 | 1.03 | 1.38 | 1.59 | 2.72 | 1.65 | 1.35 | 1.40 | 1.28 | 1.42 |
| 5 | 0.86 | 1.27 | 1.13 | 0.87 | 0.84 | 1.12 | 0.93 | 1.08 | 1.41 | 1.49 | 2.61 | 1.64 | 1.76 | 1.43 | 1.75 | 1.68 |
| 6 | 0.90 | 1.24 | 1.21 | 0.67 | 0.99 | 1.11 | 0.90 | 0.97 | 1.87 | 1.96 | 2.23 | 1.71 | 1.79 | 1.96 | 2.12 | 2.03 |
| 7 | 1.28 | 1.12 | 1.26 | 0.97 | 0.95 | 1.11 | 0.97 | 1.18 | 1.79 | 1.75 | 3.05 | 2.04 | 1.88 | 1.64 | 2.40 | 2.20 |
| 8 | 1.14 | 1.16 | 1.05 | 1.21 | 0.95 | 1.08 | 0.99 | 1.12 | 2.30 | 1.64 | 3.20 | 2.33 | 2.03 | 2.18 | 2.42 | 1.84 |
| 9 | 0.87 | 1.06 | 0.93 | 0.96 | 0.98 | 0.91 | 0.79 | 0.92 | 1.77 | 1.47 | 3.99 | 1.63 | 2.09 | 2.15 | 2.46 | 1.80 |
| 11 | 1.11 | 0.99 | 1.11 | 1.14 | 0.86 | 0.95 | 0.86 | 0.84 | 2.23 | 1.90 | 3.21 | 2.62 | 3.87 | 2.64 | 2.78 | 2.23 |

**GLYCINE**

Concentration (µM)

| Time (months) | WT | | | | | | | | Polg | | | | | | | |
|---|---|---|---|---|---|---|---|---|---|---|---|---|---|---|---|---|
| 3 | 773.82 | 757.39 | 590.56 | 550.95 | 826.09 | 642.93 | 646.10 | 730.90 | 808.18 | 807.07 | 745.55 | 806.45 | 643.22 | 715.24 | 734.87 | 764.90 |
| 4 | 267.10 | 386.90 | 319.40 | 314.10 | 309.20 | 298.30 | 307.00 | 364.90 | 381.20 | 422.70 | 485.00 | 439.50 | 406.00 | 415.70 | 388.40 | 385.10 |
| 5 | 332.60 | 486.50 | 696.80 | 268.30 | 457.90 | 320.50 | 315.80 | 312.10 | 314.90 | 398.30 | 445.10 | 376.10 | 460.70 | 347.10 | 381.40 | 430.40 |
| 6 | 393.60 | 372.90 | 353.60 | 230.30 | 387.60 | 331.70 | 325.30 | 315.50 | 353.90 | 529.80 | 344.60 | 352.40 | 347.60 | 352.40 | 380.90 | 417.30 |
| 7 | 467.60 | 364.50 | 353.20 | 303.40 | 389.00 | 599.90 | 372.40 | 361.90 | 464.60 | 329.10 | 397.40 | 376.40 | 437.60 | 334.30 | 485.90 | 537.30 |
| 8 | 403.40 | 619.00 | 384.80 | 329.90 | 370.60 | 364.90 | 325.30 | 364.60 | 373.40 | 291.70 | 397.20 | 381.20 | 384.00 | 350.90 | 401.60 | 369.60 |
| 9 | 312.80 | 345.40 | 289.10 | 292.40 | 322.40 | 343.00 | 305.20 | 305.00 | 377.80 | 304.90 | 579.40 | 366.20 | 430.50 | 409.10 | 444.60 | 365.30 |
| 11 | 363.20 | 647.70 | 357.30 | 403.50 | 292.60 | 300.00 | 359.30 | 246.20 | 310.20 | 306.60 | 397.80 | 374.30 | 425.20 | 352.80 | 402.20 | 350.20 |

*Values when normalized to WT 3 months (i.e. set WT 3 month vales to 1)*

| Time (months) | WT | | | | | | | | Polg | | | | | | | |
|---|---|---|---|---|---|---|---|---|---|---|---|---|---|---|---|---|
| 3 | 1.12 | 1.10 | 0.86 | 0.80 | 1.20 | 0.93 | 0.94 | 1.06 | 1.17 | 1.17 | 1.08 | 1.17 | 0.93 | 1.04 | 1.07 | 1.11 |
| 4 | 0.39 | 0.56 | 0.46 | 0.46 | 0.45 | 0.43 | 0.45 | 0.53 | 0.55 | 0.61 | 0.70 | 0.64 | 0.59 | 0.60 | 0.56 | 0.56 |
| 5 | 0.48 | 0.71 | 1.01 | 0.39 | 0.66 | 0.46 | 0.46 | 0.45 | 0.46 | 0.58 | 0.65 | 0.55 | 0.67 | 0.50 | 0.55 | 0.62 |
| 6 | 0.57 | 0.54 | 0.51 | 0.33 | 0.56 | 0.48 | 0.47 | 0.46 | 0.51 | 0.77 | 0.50 | 0.51 | 0.50 | 0.51 | 0.55 | 0.60 |
| 7 | 0.68 | 0.53 | 0.51 | 0.44 | 0.56 | 0.87 | 0.54 | 0.52 | 0.67 | 0.48 | 0.58 | 0.55 | 0.63 | 0.48 | 0.70 | 0.78 |
| 8 | 0.58 | 0.90 | 0.56 | 0.48 | 0.54 | 0.53 | 0.47 | 0.53 | 0.54 | 0.42 | 0.58 | 0.55 | 0.56 | 0.51 | 0.58 | 0.54 |
| 9 | 0.45 | 0.50 | 0.42 | 0.42 | 0.47 | 0.50 | 0.44 | 0.44 | 0.55 | 0.44 | 0.84 | 0.53 | 0.62 | 0.59 | 0.64 | 0.53 |
| 11 | 0.53 | 0.94 | 0.52 | 0.58 | 0.42 | 0.43 | 0.52 | 0.36 | 0.45 | 0.44 | 0.58 | 0.54 | 0.62 | 0.51 | 0.58 | 0.51 |

**VALINE**

Concentration (µM)

| Time (months) | WT | | | | | | | | Polg | | | | | | | |
|---|---|---|---|---|---|---|---|---|---|---|---|---|---|---|---|---|
| 3 | 383.59 | 326.21 | 352.75 | 385.96 | 376.73 | 309.79 | 239.32 | 235.83 | 343.94 | 366.15 | 362.01 | 384.79 | 262.51 | 300.77 | 294.64 | 281.19 |
| 4 | 371.60 | 435.00 | 404.90 | 447.30 | 357.30 | 376.10 | 345.80 | 407.50 | 342.10 | 344.00 | 355.80 | 440.70 | 293.60 | 366.70 | 266.50 | 318.10 |
| 5 | 302.30 | 445.90 | 498.40 | 320.80 | 291.40 | 467.10 | 339.30 | 465.40 | 410.50 | 404.80 | 319.80 | 458.10 | 330.30 | 344.40 | 457.90 | 395.30 |
| 6 | 342.50 | 557.80 | 531.00 | 323.80 | 352.20 | 492.80 | 346.50 | 456.80 | 389.70 | 334.20 | 258.70 | 353.40 | 349.90 | 397.30 | 485.20 | 375.40 |
| 7 | 366.60 | 474.70 | 570.40 | 423.80 | 341.50 | 410.70 | 404.90 | 472.60 | 401.40 | 377.40 | 371.30 | 383.30 | 349.40 | 319.80 | 407.90 | 356.30 |
| 8 | 441.50 | 485.90 | 364.40 | 521.70 | 392.80 | 388.20 | 362.00 | 366.40 | 419.30 | 331.60 | 352.50 | 389.00 | 332.90 | 358.80 | 408.20 | 296.20 |
| 9 | 430.40 | 434.20 | 372.40 | 430.50 | 409.80 | 380.00 | 333.70 | 384.40 | 341.40 | 323.90 | 469.70 | 396.10 | 301.20 | 462.30 | 377.50 | 362.70 |
| 11 | 453.30 | 497.40 | 459.70 | 524.70 | 363.60 | 426.90 | 330.70 | 339.00 | 417.70 | 404.20 | 384.20 | 530.30 | 447.60 | 466.90 | 616.20 | 420.70 |

*Values when normalized to WT 3 months (i.e. set WT 3 month vales to 1)*

| Time (months) | WT | | | | | | | | Polg | | | | | | | |
|---|---|---|---|---|---|---|---|---|---|---|---|---|---|---|---|---|
| 3 | 1.18 | 1.00 | 1.08 | 1.18 | 1.15 | 0.95 | 0.73 | 0.72 | 1.05 | 1.12 | 1.11 | 1.18 | 0.80 | 0.92 | 0.90 | 0.86 |
| 4 | 1.14 | 1.33 | 1.24 | 1.37 | 1.10 | 1.15 | 1.06 | 1.25 | 1.05 | 1.05 | 1.09 | 1.35 | 0.90 | 1.12 | 0.82 | 0.97 |
| 5 | 0.93 | 1.37 | 1.53 | 0.98 | 0.89 | 1.43 | 1.04 | 1.43 | 1.26 | 1.24 | 0.98 | 1.40 | 1.01 | 1.06 | 1.40 | 1.21 |
| 6 | 1.05 | 1.71 | 1.63 | 0.99 | 1.08 | 1.51 | 1.06 | 1.40 | 1.19 | 1.02 | 0.79 | 1.08 | 1.07 | 1.22 | 1.49 | 1.15 |
| 7 | 1.12 | 1.45 | 1.75 | 1.30 | 1.05 | 1.26 | 1.24 | 1.45 | 1.23 | 1.16 | 1.14 | 1.17 | 1.07 | 0.98 | 1.25 | 1.09 |
| 8 | 1.35 | 1.49 | 1.12 | 1.60 | 1.20 | 1.19 | 1.11 | 1.12 | 1.29 | 1.02 | 1.08 | 1.19 | 1.02 | 1.10 | 1.25 | 0.91 |
| 9 | 1.32 | 1.33 | 1.14 | 1.32 | 1.26 | 1.16 | 1.02 | 1.18 | 1.05 | 0.99 | 1.44 | 1.21 | 0.92 | 1.42 | 1.16 | 1.11 |
| 11 | 1.39 | 1.52 | 1.41 | 1.61 | 1.11 | 1.31 | 1.01 | 1.04 | 1.28 | 1.24 | 1.18 | 1.63 | 1.37 | 1.43 | 1.89 | 1.29 |

**LEUCINE**

Concentration (µM)

| Time (months) | WT | | | | | | | | Polg | | | | | | | |
|---|---|---|---|---|---|---|---|---|---|---|---|---|---|---|---|---|
| 3 | 229.42 | 205.26 | 207.02 | 247.99 | 241.96 | 188.83 | 138.64 | 152.32 | 238.31 | 255.17 | 226.06 | 249.43 | 163.29 | 185.78 | 185.03 | 164.48 |
| 4 | 237.80 | 291.80 | 260.70 | 283.80 | 259.50 | 225.20 | 219.70 | 250.10 | 218.60 | 216.30 | 234.60 | 286.60 | 203.10 | 232.00 | 173.30 | 215.60 |
| 5 | 187.10 | 289.10 | 315.90 | 185.00 | 165.00 | 312.10 | 204.30 | 285.70 | 271.10 | 261.70 | 202.20 | 318.00 | 217.70 | 217.90 | 307.60 | 267.90 |
| 6 | 194.60 | 357.00 | 326.10 | 172.80 | 221.30 | 300.40 | 223.00 | 284.80 | 248.60 | 206.40 | 151.70 | 214.10 | 243.10 | 247.40 | 325.50 | 248.80 |
| 7 | 232.80 | 297.10 | 393.00 | 278.90 | 204.30 | 261.30 | 258.60 | 300.30 | 250.40 | 230.80 | 230.70 | 246.40 | 226.90 | 205.60 | 246.10 | 233.40 |
| 8 | 274.80 | 307.80 | 220.80 | 335.90 | 249.20 | 251.20 | 225.50 | 227.30 | 264.30 | 200.20 | 232.90 | 244.10 | 215.70 | 217.10 | 252.00 | 199.50 |
| 9 | 277.50 | 263.00 | 236.30 | 261.30 | 271.10 | 234.00 | 183.00 | 223.70 | 214.60 | 191.70 | 323.90 | 270.80 | 188.80 | 296.10 | 225.90 | 232.70 |
| 11 | 276.30 | 318.00 | 300.30 | 317.70 | 206.60 | 248.40 | 201.10 | 198.50 | 238.70 | 232.70 | 239.00 | 308.80 | 265.90 | 256.50 | 350.30 | 259.10 |

*Values when normalized to WT 3 months (i.e. set WT 3 month vales to 1)*

| Time (months) | WT | | | | | | | | Polg | | | | | | | |
|---|---|---|---|---|---|---|---|---|---|---|---|---|---|---|---|---|
| 3 | 1.14 | 1.02 | 1.03 | 1.23 | 1.20 | 0.94 | 0.69 | 0.76 | 1.18 | 1.27 | 1.12 | 1.24 | 0.81 | 0.92 | 0.92 | 0.82 |
| 4 | 1.18 | 1.45 | 1.29 | 1.41 | 1.29 | 1.12 | 1.09 | 1.24 | 1.09 | 1.07 | 1.16 | 1.42 | 1.01 | 1.15 | 0.86 | 1.07 |
| 5 | 0.93 | 1.44 | 1.57 | 0.92 | 0.82 | 1.55 | 1.01 | 1.42 | 1.35 | 1.30 | 1.00 | 1.58 | 1.08 | 1.08 | 1.53 | 1.33 |
| 6 | 0.97 | 1.77 | 1.62 | 0.86 | 1.10 | 1.49 | 1.11 | 1.41 | 1.23 | 1.02 | 0.75 | 1.06 | 1.21 | 1.23 | 1.62 | 1.24 |
| 7 | 1.16 | 1.47 | 1.95 | 1.38 | 1.01 | 1.30 | 1.28 | 1.49 | 1.24 | 1.15 | 1.15 | 1.22 | 1.13 | 1.02 | 1.22 | 1.16 |
| 8 | 1.36 | 1.53 | 1.10 | 1.67 | 1.24 | 1.25 | 1.12 | 1.13 | 1.31 | 0.99 | 1.16 | 1.21 | 1.07 | 1.08 | 1.25 | 0.99 |
| 9 | 1.38 | 1.31 | 1.17 | 1.30 | 1.35 | 1.16 | 0.91 | 1.11 | 1.07 | 0.95 | 1.61 | 1.34 | 0.94 | 1.47 | 1.12 | 1.16 |
| 11 | 1.37 | 1.58 | 1.49 | 1.58 | 1.03 | 1.23 | 1.00 | 0.99 | 1.19 | 1.16 | 1.19 | 1.53 | 1.32 | 1.27 | 1.74 | 1.29 |

**ISOLEUCINE**

Concentration (µM)

| Time (months) | WT | | | | | | | | Polg | | | | | | | |
|---|---|---|---|---|---|---|---|---|---|---|---|---|---|---|---|---|
| 3 | 148.18 | 125.09 | 125.45 | 150.83 | 148.85 | 115.19 | 91.00 | 91.24 | 137.45 | 156.47 | 151.61 | 151.16 | 107.01 | 117.34 | 119.63 | 113.87 |
| 4 | 144.90 | 170.40 | 159.90 | 166.30 | 154.30 | 122.70 | 130.70 | 144.60 | 128.20 | 127.20 | 142.10 | 167.40 | 120.50 | 147.40 | 102.30 | 127.10 |
| 5 | 109.10 | 172.30 | 184.80 | 105.30 | 99.80 | 183.70 | 120.10 | 166.60 | 158.50 | 157.40 | 124.90 | 182.60 | 134.60 | 129.20 | 178.10 | 159.40 |
| 6 | 127.60 | 206.60 | 190.70 | 111.90 | 132.80 | 179.70 | 134.90 | 170.60 | 163.20 | 128.70 | 99.90 | 148.50 | 149.00 | 149.60 | 189.90 | 150.20 |
| 7 | 137.50 | 177.70 | 225.50 | 159.50 | 120.30 | 152.10 | 153.20 | 176.30 | 147.60 | 141.90 | 134.90 | 152.20 | 139.70 | 125.10 | 151.60 | 145.90 |
| 8 | 167.10 | 181.00 | 128.60 | 202.80 | 143.30 | 147.70 | 141.70 | 138.70 | 167.10 | 132.20 | 143.20 | 155.20 | 144.30 | 148.10 | 157.30 | 125.90 |
| 9 | 165.00 | 162.70 | 145.10 | 160.00 | 166.00 | 142.80 | 116.50 | 137.20 | 138.70 | 124.60 | 200.50 | 165.10 | 123.00 | 180.60 | 148.20 | 150.20 |
| 11 | 161.80 | 190.80 | 183.50 | 188.30 | 118.60 | 144.60 | 117.70 | 118.20 | 154.70 | 145.70 | 152.30 | 195.80 | 183.00 | 168.60 | 224.10 | 162.40 |

*Values when normalized to WT 3 months (i.e. set WT 3 month vales to 1)*

| Time (months) | WT | | | | | | | | Polg | | | | | | | |
|---|---|---|---|---|---|---|---|---|---|---|---|---|---|---|---|---|
| 3 | 1.19 | 1.00 | 1.01 | 1.21 | 1.20 | 0.93 | 0.73 | 0.73 | 1.10 | 1.26 | 1.22 | 1.21 | 0.86 | 0.94 | 0.96 | 0.91 |
| 4 | 1.16 | 1.37 | 1.28 | 1.34 | 1.24 | 0.99 | 1.05 | 1.16 | 1.03 | 1.02 | 1.14 | 1.34 | 0.97 | 1.18 | 0.82 | 1.02 |
| 5 | 0.88 | 1.38 | 1.48 | 0.85 | 0.80 | 1.48 | 0.96 | 1.34 | 1.27 | 1.26 | 1.00 | 1.47 | 1.08 | 1.04 | 1.43 | 1.28 |
| 6 | 1.03 | 1.66 | 1.53 | 0.90 | 1.07 | 1.44 | 1.08 | 1.37 | 1.31 | 1.03 | 0.80 | 1.19 | 1.20 | 1.20 | 1.53 | 1.21 |
| 7 | 1.10 | 1.43 | 1.81 | 1.28 | 0.97 | 1.22 | 1.23 | 1.42 | 1.19 | 1.14 | 1.08 | 1.22 | 1.12 | 1.00 | 1.22 | 1.17 |

| | | | | | | | | | | | | | | | | |
|---|---|---|---|---|---|---|---|---|---|---|---|---|---|---|---|---|
| 8 | 1.34 | 1.45 | 1.03 | 1.63 | 1.15 | 1.19 | 1.14 | 1.11 | 1.34 | 1.06 | 1.15 | 1.25 | 1.16 | 1.19 | 1.26 | 1.01 |
| 9 | 1.33 | 1.31 | 1.17 | 1.29 | 1.33 | 1.15 | 0.94 | 1.10 | 1.11 | 1.00 | 1.61 | 1.33 | 0.99 | 1.45 | 1.19 | 1.21 |
| 11 | 1.30 | 1.53 | 1.47 | 1.51 | 0.95 | 1.16 | 0.95 | 0.95 | 1.24 | 1.17 | 1.22 | 1.57 | 1.47 | 1.35 | 1.80 | 1.30 |

**SERINE**

Concentration (µM)

| Time (months) | WT | | | | | | | | Polg | | | | | | | |
|---|---|---|---|---|---|---|---|---|---|---|---|---|---|---|---|---|
| 3 | 196.00 | 136.69 | 116.14 | 146.25 | 138.84 | 124.71 | 108.26 | 128.35 | 157.79 | 170.60 | 261.71 | 160.84 | 135.52 | 193.78 | 171.68 | 147.35 |
| 4 | 115.50 | 156.00 | 122.30 | 123.80 | 144.60 | 125.20 | 111.80 | 144.00 | 151.00 | 196.70 | 269.30 | 181.20 | 150.90 | 181.30 | 156.00 | 148.80 |
| 5 | 115.60 | 165.80 | 135.70 | 101.10 | 125.00 | 126.90 | 120.70 | 127.80 | 139.90 | 160.20 | 271.00 | 174.00 | 192.50 | 160.20 | 184.90 | 191.70 |
| 6 | 149.70 | 183.70 | 192.80 | 101.50 | 168.60 | 149.10 | 126.90 | 133.40 | 158.80 | 159.80 | 199.90 | 141.00 | 156.80 | 172.20 | 195.30 | 187.40 |
| 7 | 184.60 | 144.90 | 174.00 | 128.10 | 163.10 | 155.60 | 132.70 | 142.20 | 149.50 | 146.80 | 263.90 | 159.30 | 174.40 | 142.00 | 204.10 | 195.80 |
| 8 | 161.90 | 134.50 | 158.70 | 153.00 | 152.70 | 158.90 | 129.60 | 139.10 | 177.20 | 132.00 | 262.10 | 178.40 | 151.70 | 157.60 | 191.50 | 156.10 |
| 9 | 126.20 | 151.00 | 130.30 | 125.80 | 145.90 | 134.50 | 116.40 | 135.00 | 160.70 | 127.80 | 399.60 | 158.60 | 176.40 | 189.10 | 199.00 | 164.70 |
| 11 | 165.70 | 158.50 | 158.40 | 162.60 | 126.00 | 133.00 | 156.30 | 111.90 | 195.10 | 168.00 | 262.70 | 245.10 | 260.50 | 211.90 | 277.60 | 202.50 |

*Values when normalized to WT 3 months (i.e. set WT 3 month vales to 1)*

| Time (months) | WT | | | | | | | | Polg | | | | | | | |
|---|---|---|---|---|---|---|---|---|---|---|---|---|---|---|---|---|
| 3 | 1.43 | 1.00 | 0.85 | 1.07 | 1.01 | 0.91 | 0.79 | 0.94 | 1.15 | 1.25 | 1.91 | 1.17 | 0.99 | 1.42 | 1.25 | 1.08 |
| 4 | 0.84 | 1.14 | 0.89 | 0.90 | 1.06 | 0.91 | 0.82 | 1.05 | 1.10 | 1.44 | 1.97 | 1.32 | 1.10 | 1.32 | 1.14 | 1.09 |
| 5 | 0.84 | 1.21 | 0.99 | 0.74 | 0.91 | 0.93 | 0.88 | 0.93 | 1.02 | 1.17 | 1.98 | 1.27 | 1.41 | 1.17 | 1.35 | 1.40 |
| 6 | 1.09 | 1.34 | 1.41 | 0.74 | 1.23 | 1.09 | 0.93 | 0.97 | 1.16 | 1.17 | 1.46 | 1.03 | 1.15 | 1.26 | 1.43 | 1.37 |
| 7 | 1.35 | 1.06 | 1.27 | 0.94 | 1.19 | 1.14 | 0.97 | 1.04 | 1.09 | 1.07 | 1.93 | 1.16 | 1.27 | 1.04 | 1.49 | 1.43 |
| 8 | 1.18 | 0.98 | 1.16 | 1.12 | 1.12 | 1.16 | 0.95 | 1.02 | 1.29 | 0.96 | 1.91 | 1.30 | 1.11 | 1.15 | 1.40 | 1.14 |
| 9 | 0.92 | 1.10 | 0.95 | 0.92 | 1.07 | 0.98 | 0.85 | 0.99 | 1.17 | 0.93 | 2.92 | 1.16 | 1.29 | 1.38 | 1.45 | 1.20 |
| 11 | 1.21 | 1.16 | 1.16 | 1.19 | 0.92 | 0.97 | 1.14 | 0.82 | 1.43 | 1.23 | 1.92 | 1.79 | 1.90 | 1.55 | 2.03 | 1.48 |

**THREONINE**

Concentration (µM)

| Time (months) | WT | | | | | | | | Polg | | | | | | | |
|---|---|---|---|---|---|---|---|---|---|---|---|---|---|---|---|---|
| 3 | 177.26 | 224.44 | 150.38 | 176.15 | 172.79 | 149.13 | 127.18 | 111.37 | 188.98 | 212.93 | 203.27 | 212.25 | 142.73 | 189.44 | 182.29 | 165.92 |
| 4 | 171.90 | 232.90 | 163.70 | 207.20 | 158.00 | 166.00 | 194.30 | 218.20 | 184.80 | 201.90 | 185.20 | 252.70 | 183.60 | 195.70 | 169.00 | 180.90 |
| 5 | 164.80 | 252.50 | 238.30 | 142.10 | 165.90 | 219.30 | 167.40 | 227.80 | 218.00 | 231.60 | 180.90 | 269.00 | 247.50 | 215.70 | 288.30 | 247.90 |
| 6 | 175.20 | 270.30 | 251.80 | 121.50 | 188.50 | 213.00 | 170.30 | 195.70 | 217.60 | 204.70 | 136.80 | 180.70 | 210.10 | 225.00 | 299.30 | 236.90 |
| 7 | 239.60 | 237.90 | 249.20 | 202.60 | 208.20 | 220.30 | 194.90 | 234.20 | 238.80 | 195.80 | 170.70 | 226.90 | 247.30 | 216.40 | 310.60 | 240.60 |
| 8 | 222.10 | 199.60 | 173.10 | 228.80 | 181.40 | 194.60 | 180.80 | 195.90 | 278.50 | 183.90 | 182.10 | 243.40 | 205.90 | 220.10 | 255.40 | 181.30 |
| 9 | 184.30 | 214.60 | 179.20 | 184.50 | 179.10 | 179.20 | 180.10 | 196.00 | 251.10 | 200.90 | 362.00 | 235.60 | 252.10 | 310.10 | 283.40 | 238.30 |
| 11 | 208.10 | 206.90 | 203.70 | 216.00 | 148.00 | 173.40 | 194.70 | 154.50 | 257.50 | 217.80 | 199.50 | 319.60 | 338.80 | 305.00 | 377.10 | 241.10 |

*Values when normalized to WT 3 months (i.e. set WT 3 month vales to 1)*

| Time (months) | WT | | | | | | | | Polg | | | | | | | |
|---|---|---|---|---|---|---|---|---|---|---|---|---|---|---|---|---|
| 3 | 1.10 | 1.39 | 0.93 | 1.09 | 1.07 | 0.93 | 0.79 | 0.69 | 1.17 | 1.32 | 1.26 | 1.32 | 0.89 | 1.18 | 1.13 | 1.03 |
| 4 | 1.07 | 1.45 | 1.02 | 1.29 | 0.98 | 1.03 | 1.21 | 1.35 | 1.15 | 1.25 | 1.15 | 1.57 | 1.14 | 1.21 | 1.05 | 1.12 |
| 5 | 1.02 | 1.57 | 1.48 | 0.88 | 1.03 | 1.36 | 1.04 | 1.41 | 1.35 | 1.44 | 1.12 | 1.67 | 1.54 | 1.34 | 1.79 | 1.54 |
| 6 | 1.09 | 1.68 | 1.56 | 0.75 | 1.17 | 1.32 | 1.06 | 1.21 | 1.35 | 1.27 | 0.85 | 1.12 | 1.30 | 1.40 | 1.86 | 1.47 |
| 7 | 1.49 | 1.48 | 1.55 | 1.26 | 1.29 | 1.37 | 1.21 | 1.45 | 1.48 | 1.22 | 1.06 | 1.41 | 1.54 | 1.34 | 1.93 | 1.49 |
| 8 | 1.38 | 1.24 | 1.07 | 1.42 | 1.13 | 1.21 | 1.12 | 1.22 | 1.73 | 1.14 | 1.13 | 1.51 | 1.28 | 1.37 | 1.59 | 1.13 |
| 9 | 1.14 | 1.33 | 1.11 | 1.15 | 1.11 | 1.11 | 1.12 | 1.22 | 1.56 | 1.25 | 2.25 | 1.46 | 1.56 | 1.93 | 1.76 | 1.48 |
| 11 | 1.29 | 1.28 | 1.26 | 1.34 | 0.92 | 1.08 | 1.21 | 0.96 | 1.60 | 1.35 | 1.24 | 1.98 | 2.10 | 1.89 | 2.34 | 1.50 |

**METHIONINE**

Concentration (µM)

| Time (months) | WT | | | | | | | | Polg | | | | | | | |
|---|---|---|---|---|---|---|---|---|---|---|---|---|---|---|---|---|
| 3 | 79.92 | 55.41 | 51.08 | 65.93 | 63.76 | 53.16 | 41.51 | 44.32 | 73.49 | 85.51 | 77.08 | 83.60 | 56.99 | 69.45 | 67.16 | 66.84 |
| 4 | 56.10 | 81.00 | 57.50 | 65.40 | 53.20 | 56.70 | 70.00 | 80.10 | 65.20 | 77.70 | 74.50 | 86.60 | 64.50 | 80.80 | 59.20 | 69.40 |
| 5 | 54.80 | 95.70 | 80.00 | 46.10 | 55.20 | 71.00 | 60.10 | 75.70 | 72.10 | 76.40 | 67.90 | 83.90 | 81.30 | 68.80 | 96.00 | 86.90 |
| 6 | 83.70 | 89.10 | 78.20 | 51.60 | 69.00 | 68.00 | 65.40 | 74.80 | 73.80 | 68.70 | 56.30 | 62.30 | 73.80 | 70.00 | 95.10 | 82.50 |
| 7 | 79.10 | 73.00 | 89.60 | 70.20 | 62.90 | 66.00 | 64.30 | 74.60 | 73.20 | 58.30 | 65.70 | 66.70 | 70.60 | 57.80 | 81.80 | 74.90 |
| 8 | 73.20 | 66.10 | 59.20 | 75.50 | 58.80 | 59.80 | 61.50 | 64.10 | 72.90 | 53.60 | 76.90 | 69.20 | 62.80 | 63.40 | 74.60 | 53.50 |
| 9 | 59.10 | 67.10 | 54.10 | 59.80 | 55.90 | 54.50 | 50.60 | 64.00 | 64.40 | 52.80 | 118.50 | 62.70 | 61.60 | 81.10 | 75.50 | 64.90 |
| 11 | 67.10 | 66.40 | 68.60 | 73.30 | 49.70 | 52.50 | 59.90 | 50.40 | 62.60 | 56.40 | 72.20 | 79.40 | 68.70 | 61.70 | 79.00 | 58.70 |

*Values when normalized to WT 3 months (i.e. set WT 3 month vales to 1)*

| Time (months) | WT | | | | | | | | Polg | | | | | | | |
|---|---|---|---|---|---|---|---|---|---|---|---|---|---|---|---|---|
| 3 | 1.40 | 0.97 | 0.90 | 1.16 | 1.12 | 0.93 | 0.73 | 0.78 | 1.29 | 1.50 | 1.35 | 1.47 | 1.00 | 1.22 | 1.18 | 1.18 |
| 4 | 0.99 | 1.42 | 1.01 | 1.15 | 0.94 | 1.00 | 1.23 | 1.41 | 1.15 | 1.37 | 1.31 | 1.52 | 1.13 | 1.42 | 1.04 | 1.22 |
| 5 | 0.96 | 1.68 | 1.41 | 0.81 | 0.97 | 1.25 | 1.06 | 1.33 | 1.27 | 1.34 | 1.19 | 1.47 | 1.43 | 1.21 | 1.69 | 1.53 |
| 6 | 1.47 | 1.57 | 1.37 | 0.91 | 1.21 | 1.20 | 1.15 | 1.31 | 1.30 | 1.21 | 0.99 | 1.10 | 1.30 | 1.23 | 1.67 | 1.45 |
| 7 | 1.39 | 1.28 | 1.58 | 1.23 | 1.11 | 1.16 | 1.13 | 1.31 | 1.29 | 1.02 | 1.15 | 1.17 | 1.24 | 1.02 | 1.44 | 1.32 |
| 8 | 1.29 | 1.16 | 1.04 | 1.33 | 1.03 | 1.05 | 1.08 | 1.13 | 1.28 | 0.94 | 1.35 | 1.22 | 1.10 | 1.11 | 1.31 | 0.94 |
| 9 | 1.04 | 1.18 | 0.95 | 1.05 | 0.98 | 0.96 | 0.89 | 1.13 | 1.13 | 0.93 | 2.08 | 1.10 | 1.08 | 1.43 | 1.33 | 1.14 |
| 11 | 1.18 | 1.17 | 1.21 | 1.29 | 0.87 | 0.92 | 1.05 | 0.89 | 1.10 | 0.99 | 1.27 | 1.40 | 1.21 | 1.08 | 1.39 | 1.03 |

**PHENYLALANINE**

Concentration (µM)

| Time (months) | WT | | | | | | | | Polg | | | | | | | |
|---|---|---|---|---|---|---|---|---|---|---|---|---|---|---|---|---|
| 3 | 113.04 | 111.03 | 104.60 | 110.82 | 104.32 | 93.90 | 85.67 | 85.68 | 110.20 | 114.17 | 175.49 | 116.50 | 89.42 | 104.50 | 99.01 | 86.98 |
| 4 | 97.50 | 115.90 | 108.40 | 111.70 | 106.30 | 94.70 | 96.70 | 106.60 | 85.30 | 95.90 | 174.20 | 102.50 | 96.90 | 97.00 | 77.80 | 99.10 |
| 5 | 83.30 | 116.50 | 119.90 | 84.40 | 84.90 | 119.40 | 87.10 | 109.70 | 104.30 | 106.50 | 163.30 | 119.30 | 97.10 | 100.40 | 116.00 | 115.30 |
| 6 | 94.90 | 134.30 | 129.60 | 81.30 | 106.50 | 111.50 | 95.80 | 116.40 | 95.00 | 86.90 | 121.00 | 82.10 | 96.30 | 97.30 | 116.40 | 98.30 |
| 7 | 105.60 | 123.50 | 146.10 | 115.90 | 100.90 | 115.70 | 100.90 | 114.90 | 94.30 | 89.70 | 163.80 | 89.00 | 92.60 | 80.50 | 98.70 | 94.40 |
| 8 | 114.20 | 129.80 | 105.80 | 137.30 | 115.80 | 113.80 | 104.50 | 102.40 | 94.90 | 77.00 | 164.40 | 83.30 | 79.20 | 73.50 | 87.00 | 74.80 |
| 9 | 102.60 | 120.80 | 102.00 | 108.90 | 111.40 | 106.60 | 85.30 | 106.80 | 77.00 | 71.00 | 219.00 | 86.60 | 73.90 | 98.30 | 82.20 | 84.00 |
| 11 | 109.30 | 113.60 | 110.30 | 122.10 | 90.30 | 102.90 | 91.00 | 93.50 | 67.90 | 67.50 | 123.10 | 77.40 | 66.90 | 70.00 | 78.30 | 72.40 |

*Values when normalized to WT 3 months (i.e. set WT 3 month vales to 1)*

| Time (months) | WT | | | | | | | | Polg | | | | | | | |
|---|---|---|---|---|---|---|---|---|---|---|---|---|---|---|---|---|
| 3 | 1.12 | 1.10 | 1.03 | 1.10 | 1.03 | 0.93 | 0.85 | 0.85 | 1.09 | 1.13 | 1.74 | 1.15 | 0.88 | 1.03 | 0.98 | 0.86 |
| 4 | 0.96 | 1.15 | 1.07 | 1.10 | 1.05 | 0.94 | 0.96 | 1.05 | 0.84 | 0.95 | 1.72 | 1.01 | 0.96 | 0.96 | 0.77 | 0.98 |
| 5 | 0.82 | 1.15 | 1.19 | 0.83 | 0.84 | 1.18 | 0.86 | 1.08 | 1.03 | 1.05 | 1.61 | 1.18 | 0.96 | 0.99 | 1.15 | 1.14 |
| 6 | 0.94 | 1.33 | 1.28 | 0.80 | 1.05 | 1.10 | 0.95 | 1.15 | 0.94 | 0.86 | 1.20 | 0.81 | 0.95 | 0.96 | 1.15 | 0.97 |
| 7 | 1.04 | 1.22 | 1.44 | 1.15 | 1.00 | 1.14 | 1.00 | 1.14 | 0.93 | 0.89 | 1.62 | 0.88 | 0.92 | 0.80 | 0.98 | 0.93 |
| 8 | 1.13 | 1.28 | 1.05 | 1.36 | 1.15 | 1.13 | 1.03 | 1.01 | 0.94 | 0.76 | 1.63 | 0.82 | 0.78 | 0.73 | 0.86 | 0.74 |
| 9 | 1.01 | 1.19 | 1.01 | 1.08 | 1.10 | 1.05 | 0.84 | 1.06 | 0.76 | 0.70 | 2.17 | 0.86 | 0.73 | 0.97 | 0.81 | 0.83 |
| 11 | 1.08 | 1.12 | 1.09 | 1.21 | 0.89 | 1.02 | 0.90 | 0.92 | 0.67 | 0.67 | 1.22 | 0.77 | 0.66 | 0.69 | 0.77 | 0.72 |

**ASPARTATE**

Concentration (µM)

| Time (months) | WT | | | | | | | | Polg | | | | | | | |
|---|---|---|---|---|---|---|---|---|---|---|---|---|---|---|---|---|
| 3 | 27.01 | 18.12 | 18.21 | 17.98 | 25.07 | 23.60 | 28.75 | 17.36 | 41.81 | 23.22 | 24.61 | 17.14 | 18.52 | 36.48 | 23.90 | 22.69 |
| 4 | 13.90 | 25.40 | 18.40 | 16.20 | 26.40 | 21.70 | 12.00 | 19.20 | 29.30 | 38.60 | 27.60 | 24.50 | 14.80 | 28.80 | 21.00 | 16.70 |
| 5 | 11.80 | 14.50 | 22.30 | 12.60 | 18.10 | 13.30 | 14.40 | 14.60 | 12.60 | 11.90 | 17.90 | 10.80 | 12.70 | 13.90 | 12.30 | 16.10 |
| 6 | 24.00 | 19.60 | 19.00 | 12.00 | 20.00 | 15.70 | 13.20 | 14.90 | 13.10 | 20.80 | 9.80 | 13.10 | 11.70 | 15.60 | 12.30 | 17.40 |
| 7 | 14.30 | 13.30 | 18.30 | 14.30 | 14.50 | 11.30 | 13.00 | 14.50 | 8.20 | 12.60 | 10.80 | 8.80 | 11.00 | 8.20 | 10.10 | 12.80 |
| 8 | 63.40 | 12.80 | 28.20 | 13.80 | 19.70 | 15.90 | 12.50 | 13.30 | 11.60 | 11.00 | 11.80 | 12.10 | 8.70 | 9.80 | 13.10 | 13.40 |
| 9 | 19.40 | 17.70 | 15.30 | 16.30 | 19.30 | 13.30 | 11.20 | 12.30 | 14.90 | 7.90 | 31.60 | 19.40 | 12.20 | 13.20 | 10.00 | 11.80 |
| 11 | 15.60 | 27.50 | 12.70 | 15.80 | 20.00 | 14.90 | 15.00 | 11.60 | 8.70 | 14.00 | 8.10 | 8.60 | 7.40 | 8.10 | 9.60 | 11.10 |

*Values when normalized to WT 3 months (i.e. set WT 3 month vales to 1)*

| Time (months) | WT | | | | | | | | Polg | | | | | | | |
|---|---|---|---|---|---|---|---|---|---|---|---|---|---|---|---|---|
| 3 | 1.23 | 0.82 | 0.83 | 0.82 | 1.14 | 1.07 | 1.31 | 0.79 | 1.90 | 1.05 | 1.12 | 0.78 | 0.84 | 1.66 | 1.09 | 1.03 |
| 4 | 0.63 | 1.15 | 0.84 | 0.74 | 1.20 | 0.99 | 0.55 | 0.87 | 1.33 | 1.75 | 1.25 | 1.11 | 0.67 | 1.31 | 0.95 | 0.76 |
| 5 | 0.54 | 0.66 | 1.01 | 0.57 | 0.82 | 0.60 | 0.65 | 0.66 | 0.57 | 0.54 | 0.81 | 0.49 | 0.58 | 0.63 | 0.56 | 0.73 |

| | | | | | | | | | | | | | | | | |
|---|---|---|---|---|---|---|---|---|---|---|---|---|---|---|---|---|
| 6 | 1.09 | 0.89 | 0.86 | 0.55 | 0.91 | 0.71 | 0.60 | 0.68 | 0.60 | 0.94 | 0.45 | 0.60 | 0.53 | 0.71 | 0.56 | 0.79 |
| 7 | 0.65 | 0.60 | 0.83 | 0.65 | 0.66 | 0.51 | 0.59 | 0.66 | 0.37 | 0.57 | 0.49 | 0.40 | 0.50 | 0.37 | 0.46 | 0.58 |
| 8 | 2.88 | 0.58 | 1.28 | 0.63 | 0.89 | 0.72 | 0.57 | 0.60 | 0.53 | 0.50 | 0.54 | 0.55 | 0.40 | 0.45 | 0.60 | 0.61 |
| 9 | 0.88 | 0.80 | 0.70 | 0.74 | 0.88 | 0.60 | 0.51 | 0.56 | 0.68 | 0.36 | 1.44 | 0.88 | 0.55 | 0.60 | 0.45 | 0.54 |
| 11 | 0.71 | 1.25 | 0.58 | 0.72 | 0.91 | 0.68 | 0.68 | 0.53 | 0.40 | 0.64 | 0.37 | 0.39 | 0.34 | 0.37 | 0.44 | 0.50 |

**PROLINE**

Concentration (µM)

| Time (months) | WT | | | | | | | | Polg | | | | | | | |
|---|---|---|---|---|---|---|---|---|---|---|---|---|---|---|---|---|
| 3 | 126.95 | 123.95 | 106.16 | 110.27 | 137.52 | 97.00 | 64.60 | 64.52 | 155.27 | 158.11 | 207.43 | 166.30 | 83.75 | 123.79 | 110.41 | 94.33 |
| 4 | 103.90 | 153.00 | 113.50 | 123.70 | 115.00 | 105.80 | 108.70 | 138.60 | 123.90 | 138.70 | 221.90 | 190.60 | 133.10 | 132.30 | 104.70 | 130.80 |
| 5 | 100.60 | 158.20 | 150.30 | 85.70 | 90.20 | 145.10 | 115.00 | 138.00 | 164.20 | 172.90 | 237.00 | 217.10 | 182.30 | 164.10 | 211.00 | 205.30 |
| 6 | 95.70 | 184.40 | 168.50 | 67.90 | 118.70 | 142.20 | 108.70 | 137.00 | 161.40 | 172.00 | 174.30 | 140.90 | 174.00 | 208.80 | 220.00 | 213.20 |
| 7 | 154.80 | 148.10 | 170.40 | 124.60 | 126.30 | 139.80 | 121.30 | 152.60 | 191.70 | 176.50 | 257.90 | 208.00 | 210.10 | 179.90 | 225.40 | 213.10 |
| 8 | 135.80 | 142.10 | 122.20 | 154.40 | 116.80 | 120.30 | 113.00 | 116.60 | 223.70 | 150.10 | 319.50 | 229.10 | 171.80 | 172.40 | 220.10 | 170.00 |
| 9 | 119.30 | 122.80 | 116.50 | 121.50 | 118.20 | 119.80 | 93.80 | 126.40 | 181.60 | 168.00 | 482.50 | 211.10 | 204.50 | 240.20 | 237.70 | 211.80 |
| 11 | 130.40 | 128.40 | 134.50 | 137.20 | 96.40 | 111.60 | 103.20 | 86.80 | 205.90 | 205.10 | 319.60 | 274.10 | 340.00 | 255.60 | 298.30 | 259.00 |

*Values when normalized to WT 3 months (i.e. set WT 3 month vales to 1)*

| Time (months) | WT | | | | | | | | Polg | | | | | | | |
|---|---|---|---|---|---|---|---|---|---|---|---|---|---|---|---|---|
| 3 | 1.22 | 1.19 | 1.02 | 1.06 | 1.32 | 0.93 | 0.62 | 0.62 | 1.49 | 1.52 | 2.00 | 1.60 | 0.81 | 1.19 | 1.06 | 0.91 |
| 4 | 1.00 | 1.47 | 1.09 | 1.19 | 1.11 | 1.02 | 1.05 | 1.33 | 1.19 | 1.34 | 2.14 | 1.83 | 1.28 | 1.27 | 1.01 | 1.26 |
| 5 | 0.97 | 1.52 | 1.45 | 0.83 | 0.87 | 1.40 | 1.11 | 1.33 | 1.58 | 1.66 | 2.28 | 2.09 | 1.76 | 1.58 | 2.03 | 1.98 |
| 6 | 0.92 | 1.78 | 1.62 | 0.65 | 1.14 | 1.37 | 1.05 | 1.32 | 1.55 | 1.66 | 1.68 | 1.36 | 1.68 | 2.01 | 2.12 | 2.05 |
| 7 | 1.49 | 1.43 | 1.64 | 1.20 | 1.22 | 1.35 | 1.17 | 1.47 | 1.85 | 1.70 | 2.48 | 2.00 | 2.02 | 1.73 | 2.17 | 2.05 |
| 8 | 1.31 | 1.37 | 1.18 | 1.49 | 1.12 | 1.16 | 1.09 | 1.12 | 2.15 | 1.45 | 3.08 | 2.21 | 1.65 | 1.66 | 2.12 | 1.64 |
| 9 | 1.15 | 1.18 | 1.12 | 1.17 | 1.14 | 1.15 | 0.90 | 1.22 | 1.75 | 1.62 | 4.65 | 2.03 | 1.97 | 2.31 | 2.29 | 2.04 |
| 11 | 1.26 | 1.24 | 1.29 | 1.32 | 0.93 | 1.07 | 0.99 | 0.84 | 1.98 | 1.97 | 3.08 | 2.64 | 3.27 | 2.46 | 2.87 | 2.49 |

**GLUTAMATE**

Concentration (µM)

| Time (months) | WT | | | | | | | | Polg | | | | | | | |
|---|---|---|---|---|---|---|---|---|---|---|---|---|---|---|---|---|
| 3 | 49.03 | 36.59 | 40.37 | 34.80 | 70.45 | 43.64 | 42.99 | 31.62 | 66.38 | 58.44 | 62.00 | 36.77 | 40.78 | 71.99 | 48.01 | 45.67 |
| 4 | 42.30 | 56.80 | 49.20 | 47.50 | 79.80 | 64.40 | 46.10 | 57.30 | 74.80 | 77.80 | 72.20 | 64.70 | 36.60 | 70.60 | 46.60 | 36.50 |
| 5 | 30.70 | 37.10 | 70.70 | 31.60 | 48.00 | 42.30 | 35.40 | 44.20 | 38.30 | 35.50 | 52.80 | 36.20 | 32.70 | 40.20 | 44.30 | 43.50 |
| 6 | 61.20 | 48.80 | 53.60 | 34.20 | 43.40 | 38.40 | 32.00 | 38.10 | 35.20 | 55.00 | 36.50 | 35.10 | 31.00 | 48.00 | 38.00 | 50.80 |
| 7 | 31.40 | 36.30 | 46.90 | 45.20 | 33.00 | 29.70 | 35.90 | 47.10 | 29.00 | 44.80 | 44.40 | 25.00 | 37.00 | 31.60 | 34.50 | 35.00 |
| 8 | 50.90 | 34.90 | 63.50 | 40.10 | 47.40 | 38.70 | 36.80 | 36.70 | 34.10 | 33.00 | 48.60 | 34.50 | 27.40 | 29.30 | 38.80 | 37.40 |
| 9 | 29.80 | 46.50 | 33.40 | 36.50 | 38.90 | 33.80 | 30.60 | 32.00 | 31.10 | 23.10 | 91.20 | 41.00 | 31.90 | 45.30 | 31.90 | 34.20 |
| 11 | 42.60 | 63.20 | 27.40 | 39.30 | 44.80 | 39.40 | 39.40 | 30.70 | 27.60 | 38.80 | 31.80 | 22.30 | 19.10 | 22.00 | 26.60 | 27.30 |

*Values when normalized to WT 3 months (i.e. set WT 3 month vales to 1)*

| Time (months) | WT | | | | | | | | Polg | | | | | | | |
|---|---|---|---|---|---|---|---|---|---|---|---|---|---|---|---|---|
| 3 | 1.12 | 0.84 | 0.92 | 0.80 | 1.61 | 1.00 | 0.98 | 0.72 | 1.52 | 1.34 | 1.42 | 0.84 | 0.93 | 1.65 | 1.10 | 1.05 |
| 4 | 0.97 | 1.30 | 1.13 | 1.09 | 1.83 | 1.47 | 1.06 | 1.31 | 1.71 | 1.78 | 1.65 | 1.48 | 0.84 | 1.62 | 1.07 | 0.84 |
| 5 | 0.70 | 0.85 | 1.62 | 0.72 | 1.10 | 0.97 | 0.81 | 1.01 | 0.88 | 0.81 | 1.21 | 0.83 | 0.75 | 0.92 | 1.01 | 1.00 |
| 6 | 1.40 | 1.12 | 1.23 | 0.78 | 0.99 | 0.88 | 0.73 | 0.87 | 0.81 | 1.26 | 0.84 | 0.80 | 0.71 | 1.10 | 0.87 | 1.16 |
| 7 | 0.72 | 0.83 | 1.07 | 1.03 | 0.76 | 0.68 | 0.82 | 1.08 | 0.66 | 1.03 | 1.02 | 0.57 | 0.85 | 0.72 | 0.79 | 0.80 |
| 8 | 1.17 | 0.80 | 1.45 | 0.92 | 1.08 | 0.89 | 0.84 | 0.84 | 0.78 | 0.76 | 1.11 | 0.79 | 0.63 | 0.67 | 0.89 | 0.86 |
| 9 | 0.68 | 1.06 | 0.76 | 0.84 | 0.89 | 0.77 | 0.70 | 0.73 | 0.71 | 0.53 | 2.09 | 0.94 | 0.73 | 1.04 | 0.73 | 0.78 |
| 11 | 0.98 | 1.45 | 0.63 | 0.90 | 1.03 | 0.90 | 0.90 | 0.70 | 0.63 | 0.89 | 0.73 | 0.51 | 0.44 | 0.50 | 0.61 | 0.62 |

**LYSINE**

Concentration (µM)

| Time (months) | WT | | | | | | | | Polg | | | | | | | |
|---|---|---|---|---|---|---|---|---|---|---|---|---|---|---|---|---|
| 3 | 473.49 | 477.38 | 398.58 | 432.71 | 372.50 | 341.28 | 271.21 | 259.56 | 440.59 | 491.87 | 373.56 | 501.28 | 305.76 | 385.48 | 423.43 | 349.29 |
| 4 | 328.70 | 438.70 | 376.00 | 354.70 | 349.30 | 357.80 | 322.70 | 411.00 | 329.40 | 368.40 | 354.10 | 458.20 | 358.30 | 394.10 | 317.00 | 359.20 |
| 5 | 313.00 | 457.10 | 484.60 | 319.20 | 327.30 | 391.40 | 366.30 | 379.10 | 354.60 | 384.70 | 314.50 | 465.30 | 397.70 | 315.70 | 398.30 | 405.40 |
| 6 | 334.30 | 447.70 | 439.90 | 231.10 | 364.50 | 396.30 | 323.00 | 391.80 | 308.90 | 331.60 | 236.40 | 311.20 | 380.00 | 415.00 | 415.80 | 376.40 |
| 7 | 399.30 | 396.50 | 474.50 | 334.90 | 367.80 | 410.30 | 368.80 | 413.50 | 281.30 | 266.10 | 314.10 | 307.50 | 306.10 | 304.40 | 335.50 | 361.00 |
| 8 | 402.00 | 384.00 | 326.90 | 457.80 | 386.50 | 385.20 | 315.20 | 358.50 | 318.30 | 270.20 | 342.10 | 278.50 | 263.10 | 290.30 | 309.00 | 284.50 |
| 9 | 338.00 | 318.40 | 319.10 | 349.80 | 358.40 | 352.10 | 282.30 | 319.40 | 252.40 | 221.70 | 547.10 | 271.20 | 260.80 | 329.70 | 286.10 | 302.60 |
| 11 | 396.10 | 350.50 | 388.20 | 395.50 | 271.70 | 331.30 | 307.90 | 277.50 | 217.90 | 245.30 | 308.20 | 287.60 | 273.90 | 274.80 | 324.00 | 265.60 |

*Values when normalized to WT 3 months (i.e. set WT 3 month vales to 1)*

| Time (months) | WT | | | | | | | | Polg | | | | | | | |
|---|---|---|---|---|---|---|---|---|---|---|---|---|---|---|---|---|
| 3 | 1.25 | 1.26 | 1.05 | 1.14 | 0.98 | 0.90 | 0.72 | 0.69 | 1.16 | 1.30 | 0.99 | 1.32 | 0.81 | 1.02 | 1.12 | 0.92 |
| 4 | 0.87 | 1.16 | 0.99 | 0.94 | 0.92 | 0.95 | 0.85 | 1.09 | 0.87 | 0.97 | 0.94 | 1.21 | 0.95 | 1.04 | 0.84 | 0.95 |
| 5 | 0.83 | 1.21 | 1.28 | 0.84 | 0.87 | 1.03 | 0.97 | 1.00 | 0.94 | 1.02 | 0.83 | 1.23 | 1.05 | 0.83 | 1.05 | 1.07 |
| 6 | 0.88 | 1.18 | 1.16 | 0.61 | 0.96 | 1.05 | 0.85 | 1.04 | 0.82 | 0.88 | 0.62 | 0.82 | 1.00 | 1.10 | 1.10 | 0.99 |
| 7 | 1.06 | 1.05 | 1.25 | 0.89 | 0.97 | 1.08 | 0.97 | 1.09 | 0.74 | 0.70 | 0.83 | 0.81 | 0.81 | 0.80 | 0.89 | 0.95 |
| 8 | 1.06 | 1.01 | 0.86 | 1.21 | 1.02 | 1.02 | 0.83 | 0.95 | 0.84 | 0.71 | 0.90 | 0.74 | 0.70 | 0.77 | 0.82 | 0.75 |
| 9 | 0.89 | 0.84 | 0.84 | 0.92 | 0.95 | 0.93 | 0.75 | 0.84 | 0.67 | 0.59 | 1.45 | 0.72 | 0.69 | 0.87 | 0.76 | 0.80 |
| 11 | 1.05 | 0.93 | 1.03 | 1.05 | 0.72 | 0.88 | 0.81 | 0.73 | 0.58 | 0.65 | 0.81 | 0.76 | 0.72 | 0.73 | 0.86 | 0.70 |

**TYROSINE**

Concentration (µM)

| Time (months) | WT | | | | | | | | Polg | | | | | | | |
|---|---|---|---|---|---|---|---|---|---|---|---|---|---|---|---|---|
| 3 | 158.74 | 105.07 | 113.76 | 149.29 | 112.77 | 106.68 | 96.57 | 90.92 | 146.70 | 163.16 | 181.31 | 137.41 | 111.44 | 130.02 | 123.75 | 107.56 |
| 4 | 111.20 | 107.30 | 130.00 | 123.10 | 97.60 | 94.10 | 98.90 | 115.50 | 73.10 | 94.60 | 148.30 | 104.00 | 104.20 | 111.90 | 80.70 | 92.70 |
| 5 | 80.70 | 170.80 | 123.60 | 62.90 | 72.80 | 148.80 | 80.50 | 108.50 | 105.80 | 131.00 | 147.80 | 144.10 | 103.00 | 89.70 | 175.20 | 127.40 |
| 6 | 121.90 | 177.80 | 143.90 | 95.60 | 103.70 | 130.60 | 91.60 | 137.00 | 97.30 | 116.00 | 124.10 | 96.00 | 132.30 | 85.40 | 206.40 | 119.20 |
| 7 | 134.60 | 170.70 | 154.70 | 140.00 | 97.10 | 124.90 | 119.70 | 137.20 | 98.60 | 86.50 | 156.90 | 91.40 | 114.30 | 104.80 | 136.80 | 104.90 |
| 8 | 149.30 | 116.00 | 93.60 | 168.00 | 126.10 | 103.70 | 109.00 | 113.50 | 92.50 | 88.50 | 174.60 | 89.60 | 102.70 | 89.40 | 103.60 | 78.10 |
| 9 | 133.00 | 126.70 | 108.50 | 138.00 | 151.00 | 113.20 | 129.90 | 152.10 | 86.30 | 79.90 | 253.10 | 101.30 | 89.10 | 118.40 | 105.50 | 93.50 |
| 11 | 175.40 | 176.20 | 137.10 | 151.90 | 84.50 | 101.80 | 101.80 | 120.70 | 87.70 | 87.70 | 145.40 | 98.60 | 84.50 | 78.20 | 107.50 | 75.10 |

*Values when normalized to WT 3 months (i.e. set WT 3 month vales to 1)*

| Time (months) | WT | | | | | | | | Polg | | | | | | | |
|---|---|---|---|---|---|---|---|---|---|---|---|---|---|---|---|---|
| 3 | 1.36 | 0.90 | 0.97 | 1.28 | 0.97 | 0.91 | 0.83 | 0.78 | 1.26 | 1.40 | 1.55 | 1.18 | 0.95 | 1.11 | 1.06 | 0.92 |
| 4 | 0.95 | 0.92 | 1.11 | 1.05 | 0.84 | 0.81 | 0.85 | 0.99 | 0.63 | 0.81 | 1.27 | 0.89 | 0.89 | 0.96 | 0.69 | 0.79 |
| 5 | 0.69 | 1.46 | 1.06 | 0.54 | 0.62 | 1.27 | 0.69 | 0.93 | 0.91 | 1.12 | 1.27 | 1.23 | 0.88 | 0.77 | 1.50 | 1.09 |
| 6 | 1.04 | 1.52 | 1.23 | 0.82 | 0.89 | 1.12 | 0.78 | 1.17 | 0.83 | 0.99 | 1.06 | 0.82 | 1.13 | 0.73 | 1.77 | 1.02 |
| 7 | 1.15 | 1.46 | 1.33 | 1.20 | 0.83 | 1.07 | 1.03 | 1.18 | 0.84 | 0.74 | 1.34 | 0.78 | 0.98 | 0.90 | 1.17 | 0.90 |
| 8 | 1.28 | 0.99 | 0.80 | 1.44 | 1.08 | 0.89 | 0.93 | 0.97 | 0.79 | 0.76 | 1.50 | 0.77 | 0.88 | 0.77 | 0.89 | 0.67 |
| 9 | 1.14 | 1.09 | 0.93 | 1.18 | 1.29 | 0.97 | 1.11 | 1.30 | 0.74 | 0.68 | 2.17 | 0.87 | 0.76 | 1.01 | 0.90 | 0.80 |
| 11 | 1.50 | 1.51 | 1.17 | 1.30 | 0.72 | 0.87 | 0.87 | 1.03 | 0.75 | 0.75 | 1.25 | 0.84 | 0.72 | 0.67 | 0.92 | 0.64 |

**HISTIDINE**

Concentration (µM)

| Time (months) | WT | | | | | | | | Polg | | | | | | | |
|---|---|---|---|---|---|---|---|---|---|---|---|---|---|---|---|---|
| 3 | 83.23 | 77.00 | 69.37 | 72.86 | 93.96 | 72.26 | 90.63 | 71.17 | 92.55 | 83.54 | 128.29 | 96.72 | 69.07 | 88.25 | 77.16 | 75.91 |
| 4 | 69.50 | 95.40 | 73.70 | 73.10 | 75.00 | 67.10 | 73.80 | 78.90 | 89.70 | 108.30 | 137.40 | 93.50 | 85.80 | 91.80 | 78.10 | 96.50 |
| 5 | 68.50 | 78.10 | 91.00 | 57.70 | 84.90 | 80.20 | 82.30 | 80.50 | 92.00 | 89.60 | 139.30 | 102.10 | 87.70 | 92.00 | 92.60 | 106.70 |
| 6 | 86.40 | 91.60 | 102.50 | 64.80 | 88.20 | 88.70 | 80.80 | 86.90 | 104.70 | 90.60 | 86.20 | 91.30 | 93.90 | 117.40 | 102.70 | 108.60 |
| 8 | 93.00 | 93.60 | 96.10 | 86.20 | 79.40 | 95.40 | 77.40 | 77.90 | 110.70 | 85.70 | 132.90 | 111.50 | 88.90 | 91.90 | 103.50 | 95.40 |
| 9 | 79.90 | 80.60 | 69.20 | 64.50 | 68.40 | 79.10 | 59.60 | 68.70 | 82.80 | 74.20 | 211.00 | 99.10 | 70.80 | 94.00 | 87.80 | 97.40 |
| 11 | 79.90 | 73.50 | 82.50 | 79.20 | 74.30 | 79.90 | 76.10 | 64.30 | 98.40 | 88.80 | 116.00 | 112.70 | 107.20 | 97.40 | 105.70 | 103.50 |

*Values when normalized to WT 3 months (i.e. set WT 3 month vales to 1)*

| Time (months) | WT | | | | | | | | Polg | | | | | | | |
|---|---|---|---|---|---|---|---|---|---|---|---|---|---|---|---|---|
| 3 | 1.06 | 0.98 | 0.88 | 0.92 | 1.19 | 0.92 | 1.15 | 0.90 | 1.17 | 1.06 | 1.63 | 1.23 | 0.88 | 1.12 | 0.98 | 0.96 |
| 4 | 0.88 | 1.21 | 0.94 | 0.93 | 0.95 | 0.85 | 0.94 | 1.00 | 1.14 | 1.37 | 1.74 | 1.19 | 1.09 | 1.16 | 0.99 | 1.22 |

| | | | | | | | | | | | | | | | | |
|---|---|---|---|---|---|---|---|---|---|---|---|---|---|---|---|---|
| 5 | 0.87 | 0.99 | 1.15 | 0.73 | 1.08 | 1.02 | 1.04 | 1.02 | 1.17 | 1.14 | 1.77 | 1.30 | 1.11 | 1.17 | 1.17 | 1.35 |
| 6 | 1.10 | 1.16 | 1.30 | 0.82 | 1.12 | 1.13 | 1.03 | 1.10 | 1.33 | 1.15 | 1.09 | 1.16 | 1.19 | 1.49 | 1.30 | 1.38 |
| 8 | 1.18 | 1.19 | 1.22 | 1.09 | 1.01 | 1.21 | 0.98 | 0.99 | 1.40 | 1.09 | 1.69 | 1.41 | 1.13 | 1.17 | 1.31 | 1.21 |
| 9 | 1.01 | 1.02 | 0.88 | 0.82 | 0.87 | 1.00 | 0.76 | 0.87 | 1.05 | 0.94 | 2.68 | 1.26 | 0.90 | 1.19 | 1.11 | 1.24 |
| 11 | 1.01 | 0.93 | 1.05 | 1.00 | 0.94 | 1.01 | 0.97 | 0.82 | 1.25 | 1.13 | 1.47 | 1.43 | 1.36 | 1.24 | 1.34 | 1.31 |

**GLUTAMINE**

Abundance relative to internal standard

| Time (months) | WT | | | | | | | | Polg | | | | | | | |
|---|---|---|---|---|---|---|---|---|---|---|---|---|---|---|---|---|
| 3 | 7.60 | 6.85 | 5.35 | 7.79 | 7.81 | 10.87 | 6.84 | 11.01 | 5.80 | 6.03 | 10.58 | 7.80 | 8.34 | 8.94 | 10.13 | 11.91 |
| 4 | 7.79 | 10.10 | 9.34 | 8.82 | 9.67 | 11.75 | 9.48 | 11.81 | 11.39 | 14.49 | 16.76 | 8.84 | 10.34 | 14.52 | 12.35 | 8.86 |
| 5 | 8.59 | 13.26 | 8.25 | 11.02 | 8.38 | 7.09 | 10.82 | 9.82 | 15.09 | 11.17 | 23.46 | 11.86 | 9.58 | 7.68 | 13.17 | 8.84 |
| 6 | 7.23 | 8.55 | 8.07 | 6.69 | 6.98 | 9.16 | 6.26 | 4.17 | 7.87 | 6.57 | 6.44 | 7.19 | 6.31 | 7.20 | 4.24 | 6.21 |
| 7 | 6.49 | 4.75 | 4.58 | 4.82 | 7.12 | 6.63 | 5.89 | 5.58 | 9.88 | 5.37 | 7.06 | 7.15 | 4.88 | 4.69 | 7.76 | 6.13 |
| 8 | 4.08 | 9.13 | 6.92 | 7.15 | 5.74 | 6.72 | 8.19 | 7.96 | 5.65 | 7.29 | 7.57 | 7.62 | 6.35 | 6.89 | 6.36 | 5.64 |
| 9 | 4.56 | 7.27 | 5.89 | 8.03 | 5.67 | 6.28 | 4.41 | 4.61 | 5.03 | 5.35 | 8.85 | 5.32 | 4.44 | 6.10 | 7.86 | 6.18 |
| 11 | 16.74 | 12.07 | 8.66 | 12.61 | 13.45 | 13.00 | 8.03 | 7.72 | 9.83 | 13.29 | 10.40 | 10.71 | 17.25 | 12.86 | 15.95 | 16.42 |

*Values when normalized to WT 3 months (i.e. set WT 3 month vales to 1)*

| Time (months) | WT | | | | | | | | Polg | | | | | | | |
|---|---|---|---|---|---|---|---|---|---|---|---|---|---|---|---|---|
| 3 | 0.95 | 0.85 | 0.67 | 0.97 | 0.97 | 1.36 | 0.85 | 1.37 | 0.72 | 0.75 | 1.32 | 0.97 | 1.04 | 1.12 | 1.26 | 1.49 |
| 4 | 0.97 | 1.26 | 1.17 | 1.10 | 1.21 | 1.47 | 1.18 | 1.47 | 1.42 | 1.81 | 2.09 | 1.10 | 1.29 | 1.81 | 1.54 | 1.11 |
| 5 | 1.07 | 1.65 | 1.03 | 1.37 | 1.05 | 0.88 | 1.35 | 1.23 | 1.88 | 1.39 | 2.93 | 1.48 | 1.20 | 0.96 | 1.64 | 1.10 |
| 6 | 0.90 | 1.07 | 1.01 | 0.83 | 0.87 | 1.14 | 0.78 | 0.52 | 0.98 | 0.82 | 0.80 | 0.90 | 0.79 | 0.90 | 0.53 | 0.77 |
| 7 | 0.81 | 0.59 | 0.57 | 0.60 | 0.89 | 0.83 | 0.73 | 0.70 | 1.23 | 0.67 | 0.88 | 0.89 | 0.61 | 0.58 | 0.97 | 0.77 |
| 8 | 0.51 | 1.14 | 0.86 | 0.89 | 0.72 | 0.84 | 1.02 | 0.99 | 0.70 | 0.91 | 0.94 | 0.95 | 0.79 | 0.86 | 0.79 | 0.70 |
| 9 | 0.57 | 0.91 | 0.73 | 1.00 | 0.71 | 0.78 | 0.55 | 0.58 | 0.63 | 0.67 | 1.10 | 0.66 | 0.55 | 0.76 | 0.98 | 0.77 |
| 11 | 2.09 | 1.51 | 1.08 | 1.57 | 1.68 | 1.62 | 1.00 | 0.96 | 1.23 | 1.66 | 1.30 | 1.34 | 2.15 | 1.60 | 1.99 | 2.05 |

**ASPARAGINE**

Abundance relative to internal standard

| Time (months) | WT | | | | | | | | Polg | | | | | | | |
|---|---|---|---|---|---|---|---|---|---|---|---|---|---|---|---|---|
| 3 | 0.61 | 0.55 | 0.45 | 0.60 | 0.63 | 0.80 | 0.41 | 0.47 | 0.58 | 0.59 | 1.82 | 0.65 | 0.38 | 0.67 | 0.75 | 0.72 |
| 4 | 0.60 | 0.73 | 0.74 | 0.73 | 0.62 | 0.76 | 0.81 | 0.98 | 0.76 | 1.14 | 2.58 | 1.01 | 0.75 | 1.11 | 0.93 | 0.64 |
| 5 | 0.77 | 1.45 | 1.15 | 0.70 | 0.38 | 0.71 | 0.67 | 0.94 | 1.27 | 1.09 | 3.69 | 1.40 | 0.91 | 0.72 | 1.64 | 0.96 |
| 6 | 0.64 | 1.24 | 1.03 | 0.50 | 0.65 | 0.99 | 0.61 | 0.48 | 0.90 | 1.06 | 1.66 | 0.78 | 0.93 | 1.13 | 1.17 | 1.03 |
| 7 | | | | | | | | | | | | | | | | |
| 8 | 0.44 | 0.67 | 0.54 | 0.69 | 0.41 | 0.51 | 0.49 | 0.50 | 0.64 | 0.62 | 1.08 | 0.73 | 0.84 | 0.66 | 0.83 | 0.45 |
| 9 | 0.38 | 0.67 | 0.64 | 0.77 | 0.52 | 0.59 | 0.57 | 0.64 | 0.70 | 0.59 | 1.84 | 0.71 | 0.55 | 1.00 | 0.94 | 0.65 |
| 11 | 1.77 | 1.34 | 0.88 | 1.69 | 1.29 | 1.24 | 0.81 | 0.74 | 1.08 | 1.47 | 1.88 | 1.35 | 1.84 | 1.57 | 2.29 | 1.79 |

*Values when normalized to WT 3 months (i.e. set WT 3 month vales to 1)*

| Time (months) | WT | | | | | | | | Polg | | | | | | | |
|---|---|---|---|---|---|---|---|---|---|---|---|---|---|---|---|---|
| 3 | 1.07 | 0.98 | 0.79 | 1.06 | 1.12 | 1.41 | 0.73 | 0.84 | 1.02 | 1.04 | 3.22 | 1.15 | 0.66 | 1.18 | 1.33 | 1.28 |
| 4 | 1.06 | 1.29 | 1.31 | 1.29 | 1.10 | 1.34 | 1.43 | 1.73 | 1.34 | 2.01 | 4.56 | 1.78 | 1.33 | 1.96 | 1.64 | 1.13 |
| 5 | 1.36 | 2.56 | 2.03 | 1.24 | 0.67 | 1.25 | 1.18 | 1.66 | 2.24 | 1.93 | 6.52 | 2.47 | 1.61 | 1.27 | 2.90 | 1.70 |
| 6 | 1.13 | 2.19 | 1.82 | 0.88 | 1.15 | 1.75 | 1.08 | 0.85 | 1.59 | 1.87 | 2.93 | 1.38 | 1.64 | 2.00 | 2.07 | 1.82 |
| 7 | 0.00 | 0.00 | 0.00 | 0.00 | 0.00 | 0.00 | 0.00 | 0.00 | 0.00 | 0.00 | 0.00 | 0.00 | 0.00 | 0.00 | 0.00 | 0.00 |
| 8 | 0.78 | 1.18 | 0.95 | 1.22 | 0.72 | 0.90 | 0.87 | 0.88 | 1.13 | 1.10 | 1.91 | 1.29 | 1.48 | 1.17 | 1.47 | 0.80 |
| 9 | 0.67 | 1.18 | 1.13 | 1.36 | 0.92 | 1.04 | 1.01 | 1.13 | 1.24 | 1.04 | 3.25 | 1.25 | 0.97 | 1.77 | 1.66 | 1.15 |
| 11 | 3.13 | 2.37 | 1.55 | 2.99 | 2.28 | 2.19 | 1.43 | 1.31 | 1.91 | 2.60 | 3.32 | 2.39 | 3.25 | 2.77 | 4.05 | 3.16 |

**TRYPTOPHAN**

Abundance relative to internal standard

| Time (months) | WT | | | | | | | | Polg | | | | | | | |
|---|---|---|---|---|---|---|---|---|---|---|---|---|---|---|---|---|
| 3 | 0.23 | 0.15 | 0.32 | 0.19 | 0.23 | 0.17 | 0.23 | 0.20 | 0.32 | 0.22 | 0.36 | 0.20 | 0.16 | 0.19 | 0.23 | 0.23 |
| 4 | 0.33 | 0.36 | 0.29 | 0.39 | 0.35 | 1.33 | 0.60 | 0.43 | 0.27 | 0.40 | 0.34 | 0.37 | 0.34 | 0.23 | 0.25 | 0.26 |
| 5 | 0.33 | 0.29 | 0.49 | 0.24 | 0.83 | 0.37 | 0.26 | 0.34 | | 0.46 | 0.51 | 0.32 | 0.23 | 0.29 | 0.34 | 0.26 |
| 6 | 0.49 | 0.62 | 0.58 | 0.35 | 0.38 | 0.42 | 0.42 | 0.55 | 0.37 | 0.36 | 0.41 | 0.27 | 0.51 | 0.44 | 0.77 | 0.53 |
| 7 | 0.36 | 0.46 | 0.67 | 2.60 | 1.19 | 0.63 | 0.60 | 0.49 | | 0.61 | 0.72 | 0.52 | 0.62 | 0.53 | 0.57 | 1.69 |
| 8 | 0.46 | 0.42 | 0.26 | 0.55 | 0.29 | 0.38 | 0.34 | 0.42 | 0.25 | 0.15 | 0.31 | 0.22 | 0.16 | 0.25 | 0.26 | 0.23 |
| 9 | 1.00 | 0.85 | 0.63 | 0.64 | 0.59 | 0.96 | 0.75 | 0.70 | 0.32 | 0.43 | 1.58 | 0.33 | 0.43 | 0.53 | 0.54 | 0.51 |
| 11 | 0.30 | 0.43 | 0.58 | 0.45 | 1.47 | 0.30 | 0.36 | 0.34 | 0.23 | 0.25 | 0.31 | 0.48 | 0.22 | 0.26 | 0.33 | 0.26 |

*Values when normalized to WT 3 months (i.e. set WT 3 month vales to 1)*

| Time (months) | WT | | | | | | | | Polg | | | | | | | |
|---|---|---|---|---|---|---|---|---|---|---|---|---|---|---|---|---|
| 3 | 1.07 | 0.68 | 1.50 | 0.90 | 1.07 | 0.79 | 1.06 | 0.92 | 1.48 | 1.04 | 1.70 | 0.92 | 0.77 | 0.88 | 1.08 | 1.07 |
| 4 | 1.55 | 1.69 | 1.36 | 1.83 | 1.64 | 6.24 | 2.82 | 2.02 | 1.27 | 1.88 | 1.60 | 1.74 | 1.60 | 1.08 | 1.17 | 1.22 |
| 5 | 1.55 | 1.36 | 2.30 | 1.13 | 3.90 | 1.74 | 1.22 | 1.60 | 0.00 | 2.16 | 2.39 | 1.50 | 1.08 | 1.36 | 1.60 | 1.22 |
| 6 | 2.30 | 2.91 | 2.72 | 1.64 | 1.78 | 1.97 | 1.97 | 2.58 | 1.74 | 1.69 | 1.92 | 1.27 | 2.39 | 2.07 | 3.61 | 2.49 |
| 7 | 1.69 | 2.16 | 3.14 | 12.20 | 5.59 | 2.96 | 2.82 | 2.30 | 0.00 | 2.86 | 3.38 | 2.44 | 2.91 | 2.49 | 2.68 | 7.93 |
| 8 | 2.16 | 1.97 | 1.22 | 2.58 | 1.36 | 1.78 | 1.60 | 1.97 | 1.17 | 0.70 | 1.45 | 1.03 | 0.75 | 1.17 | 1.22 | 1.08 |
| 9 | 4.69 | 3.99 | 2.98 | 3.01 | 2.75 | 4.49 | 3.52 | 3.28 | 1.48 | 2.03 | 7.42 | 1.54 | 2.04 | 2.50 | 2.52 | 2.38 |
| 11 | 1.41 | 2.03 | 2.74 | 2.11 | 6.88 | 1.40 | 1.67 | 1.60 | 1.08 | 1.16 | 1.45 | 2.25 | 1.02 | 1.21 | 1.53 | 1.22 |

**Supplemental Table 2. Tissue metabolite abundance and fold-change (Polg/WT) as shown in Figure 3D**

*Abundance per mg protein, relative to internal standard*

| LIVER | WT | | | | | | | | Polg | | | | | | |
|---|---|---|---|---|---|---|---|---|---|---|---|---|---|---|---|
| Lac | 2145.0 | 1753.5 | 2006.0 | 1683.5 | 1876.8 | 1956.0 | 1593.4 | 1408.4 | 744.3 | 2133.1 | 2149.8 | 1752.6 | 2048.3 | 1573.7 | 1642.5 |
| Pyr | 19.1 | 17.4 | 25.1 | 18.5 | 23.4 | 27.2 | 17.1 | 19.8 | 11.1 | 29.8 | 27.7 | 21.6 | 19.2 | 20.0 | 20.1 |
| Cit | 16.5 | 6.2 | 23.4 | 5.0 | 9.7 | 22.0 | 9.3 | 12.2 | 6.3 | 28.2 | 22.5 | 102.8 | 15.2 | 13.9 | 71.3 |
| aKG | 3.4 | 2.6 | 3.2 | 2.4 | 3.4 | 4.5 | 2.5 | 3.3 | 1.7 | 3.3 | 3.0 | 3.1 | 2.0 | 2.4 | 2.9 |
| Suc | 275.3 | 267.7 | 224.4 | 190.1 | 218.9 | 230.1 | 222.0 | 200.9 | 206.1 | 582.9 | 572.4 | 431.8 | 494.4 | 363.3 | 522.3 |
| Fum | 119.3 | 134.1 | 117.0 | 79.5 | 125.6 | 157.1 | 103.4 | 117.7 | 176.7 | 426.8 | 429.0 | 415.1 | 354.8 | 259.2 | 355.9 |
| Mal | 80.8 | 64.1 | 74.7 | 36.4 | 49.8 | 65.3 | 43.7 | 46.3 | 91.9 | 208.8 | 232.5 | 214.2 | 161.6 | 104.8 | 183.4 |

*Abundance per mg protein, relative to internal standard*

| KIDNEY | WT | | | | | | | | Polg | | | | | | |
|---|---|---|---|---|---|---|---|---|---|---|---|---|---|---|---|
| Lac | 3787.7 | 3912.1 | 3890.3 | 3106.1 | 3588.1 | 3305.7 | 3306.6 | 3360.9 | 5027.8 | 5339.1 | 4996.6 | 5800.1 | 5112.8 | 4765.0 | 4118.4 |
| Pyr | 220.2 | 192.7 | 179.7 | 230.2 | 282.9 | 210.8 | 208.9 | 224.4 | 299.3 | 402.0 | 232.8 | 380.4 | 346.6 | 294.7 | 183.2 |
| Cit | 44.6 | 106.4 | 22.8 | 36.1 | 36.4 | 86.4 | 158.2 | 60.8 | 103.9 | 44.0 | 184.5 | 84.7 | 39.2 | 120.3 | 137.0 |
| aKG | 3.2 | 7.7 | 3.7 | 8.8 | 5.2 | 8.2 | 13.8 | 5.0 | 17.5 | 11.7 | 11.3 | 8.2 | 7.1 | 17.7 | 8.9 |
| Suc | 527.8 | 465.4 | 1020.5 | 641.0 | 486.8 | 509.9 | 632.0 | 521.0 | 1932.6 | 1190.2 | 1231.2 | 1629.1 | 1241.4 | 1339.6 | 1221.5 |
| Fum | 897.1 | 855.7 | 734.2 | 1170.5 | 977.1 | 939.1 | 938.4 | 819.0 | 2405.9 | 2173.2 | 1652.7 | 1373.3 | 1488.0 | 1519.1 | 2043.0 |
| Mal | 452.3 | 492.8 | 464.9 | 553.6 | 441.3 | 436.9 | 473.2 | 390.2 | 974.6 | 850.7 | 683.6 | 599.1 | 592.2 | 723.5 | 725.5 |

*Abundance per mg protein, relative to internal standard*

| SK MUSCLE | WT | | | | | | | | Polg | | | | | | |
|---|---|---|---|---|---|---|---|---|---|---|---|---|---|---|---|
| Lac | 2369.4 | 2980.4 | 2277.4 | 2741.1 | 2489.1 | 2897.8 | 2607.0 | 1963.9 | 2994.1 | 2753.3 | 2465.8 | 2713.9 | 3316.5 | 2917.5 | 2989.1 |
| Pyr | 87.7 | 225.7 | 203.3 | 70.5 | 121.0 | 115.7 | 131.4 | 88.4 | 81.6 | 134.6 | 83.1 | 152.0 | 163.1 | 138.0 | 109.6 |
| Cit | 110.5 | 110.6 | 95.2 | 87.7 | 92.8 | 73.9 | 91.6 | 114.6 | 93.8 | 126.5 | 83.3 | 111.6 | 186.5 | 85.9 | 73.5 |
| aKG | 12.3 | 13.1 | 10.9 | 12.7 | 11.1 | 12.7 | 11.2 | 9.8 | 11.5 | 13.8 | 10.4 | 11.5 | 16.4 | 11.1 | 12.2 |
| Suc | 579.1 | 812.5 | 629.2 | 765.9 | 654.7 | 722.3 | 714.3 | 608.7 | 540.4 | 361.6 | 242.0 | 260.9 | 402.0 | 242.8 | 303.6 |
| Fum | 388.9 | 538.0 | 455.6 | 372.4 | 347.5 | 398.4 | 416.9 | 471.3 | 404.0 | 182.0 | 197.5 | 260.8 | 487.7 | 175.3 | 194.0 |
| Mal | 108.2 | 148.1 | 149.3 | 108.7 | 96.9 | 107.0 | 113.6 | 160.5 | 143.1 | 123.6 | 81.1 | 97.6 | 166.0 | 89.2 | 95.3 |

*Fold-Change of Polg to WT (* ***Figure 3D*** *)*

| | Liver | | | | | | | Kidney | | | | | | | Heart | | | | | | |
|---|---|---|---|---|---|---|---|---|---|---|---|---|---|---|---|---|---|---|---|---|---|
| Lac | 0.41 | 1.18 | 1.19 | 0.97 | 1.14 | 0.87 | 0.91 | 1.42 | 1.51 | 1.41 | 1.64 | 1.45 | 1.35 | 1.17 | 1.18 | 1.08 | 0.97 | 1.07 | 1.31 | 1.15 | 1.18 |
| Pyr | 0.53 | 1.42 | 1.32 | 1.03 | 0.92 | 0.95 | 0.96 | 1.37 | 1.84 | 1.06 | 1.74 | 1.58 | 1.35 | 0.84 | 0.63 | 1.03 | 0.64 | 1.16 | 1.25 | 1.06 | 0.84 |
| Cit | 0.48 | 2.16 | 1.73 | 7.88 | 1.17 | 1.07 | 5.46 | 1.51 | 0.64 | 2.67 | 1.23 | 0.57 | 1.74 | 1.99 | 0.97 | 1.30 | 0.86 | 1.15 | 1.92 | 0.88 | 0.76 |
| aKG | 0.54 | 1.06 | 0.95 | 0.99 | 0.64 | 0.76 | 0.91 | 2.51 | 1.69 | 1.63 | 1.19 | 1.02 | 2.54 | 1.28 | 0.98 | 1.18 | 0.89 | 0.98 | 1.40 | 0.95 | 1.04 |
| Suc | 0.90 | 2.55 | 2.50 | 1.89 | 2.16 | 1.59 | 2.28 | 3.22 | 1.98 | 2.05 | 2.71 | 2.07 | 2.23 | 2.03 | 0.79 | 0.53 | 0.35 | 0.38 | 0.59 | 0.35 | 0.44 |
| Fum | 1.48 | 3.58 | 3.60 | 3.48 | 2.98 | 2.17 | 2.99 | 2.63 | 2.37 | 1.80 | 1.50 | 1.62 | 1.66 | 2.23 | 0.95 | 0.43 | 0.47 | 0.62 | 1.15 | 0.41 | 0.46 |
| Mal | 1.59 | 3.62 | 4.03 | 3.72 | 2.80 | 1.82 | 3.18 | 2.10 | 1.84 | 1.48 | 1.29 | 1.28 | 1.56 | 1.57 | 1.15 | 1.00 | 0.65 | 0.79 | 1.34 | 0.72 | 0.77 |

**Supplemental Table 3. Tissue amino acid concentration and fold-change (Polg/WT) as shown in Figure 3E**

*Concentration ( nmol/mg protein )*

| LIVER | WT | | | | | | | | Polg | | | | | | |
|---|---|---|---|---|---|---|---|---|---|---|---|---|---|---|---|
| Ala | 216.2 | 186.4 | 185.6 | 167.2 | 164.8 | 174.3 | 178.1 | 140.8 | 82.2 | 212.6 | 305.4 | 220.7 | 242.0 | 172.6 | 230.9 |
| Asn | 4.6 | 4.1 | 4.5 | 4.0 | 3.5 | 3.0 | 3.0 | 3.5 | 1.7 | 4.8 | 5.9 | 3.9 | 4.7 | 4.0 | 5.4 |
| Asp | 12.5 | 11.0 | 19.8 | 8.5 | 9.1 | 13.2 | 7.1 | 9.4 | 8.9 | 30.2 | 42.7 | 33.0 | 24.3 | 19.1 | 31.1 |
| Gln | 149.2 | 114.7 | 155.1 | 123.4 | 128.6 | 118.0 | 121.4 | 93.1 | 72.7 | 221.7 | 258.7 | 193.1 | 231.7 | 204.0 | 200.7 |
| Glu | 305.8 | 393.2 | 276.0 | 395.5 | 307.3 | 408.4 | 312.9 | 336.2 | 180.3 | 1050.8 | 833.3 | 1022.1 | 611.0 | 1035.5 | 1399.8 |
| Gly | 140.0 | 112.4 | 121.6 | 90.9 | 115.3 | 104.1 | 90.5 | 84.5 | 37.0 | 89.9 | 114.9 | 57.5 | 74.7 | 75.9 | 66.5 |
| His | 23.2 | 17.6 | 21.9 | 24.5 | 23.2 | 21.3 | 16.6 | 16.6 | 11.4 | 27.6 | 38.8 | 28.8 | 33.1 | 29.2 | 30.6 |
| Ile | 8.8 | 9.6 | 9.3 | 8.3 | 9.2 | 8.0 | 7.4 | 7.6 | 4.0 | 8.6 | 14.4 | 8.4 | 8.3 | 9.6 | 10.0 |
| Leu | 18.1 | 18.0 | 17.3 | 16.8 | 16.2 | 14.8 | 13.6 | 14.8 | 7.5 | 15.6 | 23.3 | 13.7 | 14.4 | 16.1 | 19.8 |
| Lys | 26.5 | 24.4 | 20.1 | 22.8 | 15.7 | 16.9 | 16.2 | 18.8 | 7.1 | 17.1 | 26.6 | 18.1 | 24.7 | 15.6 | 20.4 |
| Met | 3.9 | 3.5 | 3.4 | 3.4 | 3.1 | 3.2 | 2.4 | 3.3 | 1.5 | 4.2 | 4.2 | 2.8 | 2.9 | 2.4 | 3.6 |
| Phe | 6.9 | 5.8 | 5.9 | 5.9 | 5.8 | 4.9 | 4.6 | 4.8 | 2.1 | 5.4 | 6.6 | 4.0 | 4.7 | 4.6 | 6.2 |
| Pro | 17.8 | 17.1 | 15.5 | 15.8 | 11.2 | 10.6 | 13.1 | 14.9 | 7.7 | 15.7 | 22.0 | 13.9 | 19.9 | 15.8 | 21.7 |
| Ser | 17.7 | 12.8 | 15.3 | 11.4 | 14.6 | 12.5 | 13.9 | 12.9 | 6.2 | 17.8 | 24.9 | 18.4 | 17.8 | 15.6 | 19.6 |
| Thr | 18.6 | 13.5 | 16.4 | 15.6 | 13.7 | 11.9 | 13.4 | 12.3 | 5.7 | 13.0 | 18.2 | 13.7 | 15.8 | 14.6 | 15.1 |
| Trp | 2.3 | 2.9 | 3.0 | 2.2 | 2.6 | 2.5 | 1.8 | 2.2 | 0.9 | 2.4 | 2.8 | 1.4 | 1.6 | 1.9 | 2.4 |
| Tyr | 9.2 | 7.8 | 7.4 | 6.3 | 6.2 | 6.4 | 6.5 | 6.0 | 2.8 | 5.5 | 9.4 | 4.2 | 4.3 | 4.4 | 5.0 |
| Val | 24.8 | 26.4 | 25.7 | 24.6 | 22.6 | 21.7 | 18.9 | 21.6 | 9.5 | 19.9 | 30.8 | 18.7 | 18.5 | 21.9 | 27.9 |

*Concentration ( nmol/mg protein )*

| KIDNEY | WT | | | | | | | | Polg | | | | | | |
|---|---|---|---|---|---|---|---|---|---|---|---|---|---|---|---|
| Ala | 126.2 | 99.6 | 127.7 | 107.6 | 86.5 | 92.3 | 102.8 | 97.1 | 214.8 | 207.7 | 232.7 | 247.8 | 253.0 | 240.3 | 167.3 |
| Asn | 16.6 | 12.3 | 14.7 | 14.8 | 11.7 | 10.9 | 12.8 | 11.6 | 18.1 | 14.9 | 16.0 | 13.8 | 14.7 | 15.0 | 15.4 |
| Asp | 705.7 | 1134.1 | 441.0 | 552.0 | 983.0 | 1038.9 | 2540.9 | 739.9 | 1777.7 | 2366.5 | 1187.4 | 1186.9 | 1862.1 | 2576.2 | 404.7 |
| Gln | 115.0 | 81.9 | 114.3 | 99.3 | 87.0 | 79.2 | 84.5 | 72.5 | 126.5 | 76.7 | 112.2 | 75.3 | 78.8 | 132.0 | 70.8 |
| Glu | 895.0 | 1194.3 | 967.0 | 1050.2 | 1207.3 | 1304.3 | 1829.7 | 1143.6 | 2514.9 | 1898.7 | 1837.0 | 807.1 | 1010.2 | 2257.9 | 855.2 |
| Gly | 651.6 | 553.5 | 659.2 | 564.6 | 567.1 | 478.9 | 549.0 | 567.0 | 635.4 | 480.8 | 546.6 | 480.0 | 481.5 | 542.9 | 330.3 |
| His | 17.4 | 13.1 | 17.1 | 17.8 | 15.5 | 13.4 | 14.6 | 13.0 | 25.4 | 16.9 | 22.4 | 24.5 | 24.8 | 24.9 | 19.3 |
| Ile | 18.7 | 15.8 | 20.5 | 20.2 | 15.4 | 15.7 | 17.8 | 16.5 | 26.3 | 19.0 | 27.1 | 23.6 | 20.7 | 23.6 | 23.3 |
| Leu | 43.1 | 35.4 | 46.1 | 44.6 | 33.8 | 33.0 | 39.3 | 38.0 | 52.2 | 36.0 | 46.5 | 41.1 | 37.2 | 42.7 | 45.3 |
| Lys | 46.6 | 37.4 | 48.3 | 45.9 | 37.6 | 37.1 | 38.9 | 39.4 | 39.4 | 32.9 | 40.2 | 34.3 | 31.5 | 31.1 | 32.3 |
| Met | 17.1 | 13.5 | 16.5 | 17.4 | 12.1 | 12.2 | 14.6 | 15.6 | 19.1 | 12.8 | 16.3 | 12.6 | 12.0 | 13.7 | 12.9 |
| Phe | 17.7 | 15.2 | 18.4 | 17.4 | 14.4 | 14.2 | 14.9 | 14.4 | 18.5 | 15.2 | 13.9 | 14.6 | 13.8 | 13.5 | 17.8 |
| Pro | 22.5 | 16.7 | 19.1 | 19.2 | 16.5 | 16.3 | 18.6 | 18.6 | 37.8 | 28.3 | 37.3 | 33.3 | 36.3 | 42.8 | 30.5 |
| Ser | 46.8 | 35.2 | 38.9 | 38.7 | 34.0 | 31.1 | 35.5 | 33.7 | 62.3 | 46.2 | 57.3 | 53.8 | 57.4 | 60.7 | 41.9 |
| Thr | 38.2 | 29.7 | 35.2 | 36.1 | 26.8 | 28.3 | 32.4 | 30.7 | 55.0 | 43.1 | 51.4 | 50.7 | 55.7 | 62.9 | 38.8 |
| Trp | 6.6 | 7.9 | 7.6 | 7.5 | 6.6 | 7.0 | 7.3 | 6.9 | 7.9 | 6.3 | 9.2 | 4.8 | 4.6 | 11.3 | 6.1 |
| Tyr | 28.6 | 27.0 | 27.5 | 26.5 | 18.4 | 21.6 | 26.2 | 24.6 | 21.8 | 17.4 | 20.9 | 16.7 | 13.4 | 15.2 | 16.7 |
| Val | 44.1 | 38.7 | 47.0 | 47.7 | 31.3 | 33.5 | 39.7 | 37.4 | 58.6 | 42.0 | 50.7 | 48.3 | 43.1 | 52.0 | 57.4 |

*Concentration ( nmol/mg protein )*

| SK MUSCLE | WT | | | | | | | | Polg | | | | | | |
|---|---|---|---|---|---|---|---|---|---|---|---|---|---|---|---|
| Ala | 222.7 | 228.3 | 205.3 | 261.8 | 208.7 | 219.8 | 209.4 | 227.4 | 227.6 | 263.0 | 244.2 | 285.4 | 337.7 | 227.2 | 100.3 |
| Asn | 7.0 | 7.1 | 6.5 | 7.7 | 5.0 | 5.5 | 6.8 | 8.7 | 8.3 | 8.3 | 6.4 | 7.9 | 12.8 | 9.2 | 10.2 |
| Asp | 13.9 | 15.5 | 18.2 | 13.0 | 9.9 | 11.4 | 12.9 | 72.1 | 19.0 | 17.7 | 7.2 | 11.7 | 33.5 | 11.2 | 8.9 |
| Gln | 165.1 | 134.7 | 174.5 | 143.1 | 108.8 | 125.7 | 134.2 | 160.4 | 184.3 | 165.3 | 141.2 | 202.4 | 346.1 | 162.1 | 90.7 |
| Glu | 130.4 | 160.9 | 186.9 | 105.9 | 74.8 | 94.6 | 116.8 | 182.9 | 166.4 | 111.4 | 49.5 | 75.1 | 187.7 | 66.4 | 29.0 |
| Gly | 159.8 | 151.4 | 104.1 | 158.8 | 143.1 | 149.0 | 141.3 | 136.5 | 148.3 | 168.3 | 170.5 | 235.8 | 197.3 | 168.3 | 129.5 |
| His | 13.1 | 12.9 | 13.7 | 13.9 | 12.0 | 12.4 | 13.1 | 14.5 | 16.6 | 19.2 | 13.9 | 18.1 | 23.2 | 15.0 | 13.6 |
| Ile | 10.2 | 10.2 | 10.3 | 14.0 | 9.4 | 10.3 | 11.8 | 9.5 | 15.0 | 12.1 | 13.5 | 16.1 | 17.1 | 16.1 | 18.1 |
| Leu | 18.0 | 17.6 | 17.6 | 23.9 | 14.8 | 17.2 | 20.2 | 17.0 | 24.9 | 18.2 | 19.3 | 23.8 | 25.2 | 25.5 | 30.8 |
| Lys | 30.9 | 31.7 | 41.5 | 37.4 | 24.0 | 33.9 | 30.6 | 35.1 | 46.2 | 38.6 | 35.6 | 47.3 | 52.3 | 29.3 | 23.0 |
| Met | 9.2 | 7.5 | 7.6 | 9.9 | 5.8 | 6.8 | 7.8 | 8.2 | 7.9 | 7.3 | 7.2 | 7.5 | 8.5 | 8.2 | 8.5 |
| Phe | 10.7 | 11.4 | 10.0 | 12.9 | 9.3 | 10.7 | 10.6 | 9.6 | 8.9 | 8.5 | 7.3 | 8.6 | 9.4 | 8.9 | 12.5 |
| Pro | 15.6 | 18.6 | 13.1 | 20.5 | 13.7 | 15.7 | 16.5 | 19.6 | 32.0 | 33.6 | 30.6 | 35.2 | 39.7 | 34.9 | 35.5 |
| Ser | 22.1 | 20.2 | 17.6 | 21.8 | 17.7 | 19.0 | 20.2 | 23.5 | 31.0 | 37.9 | 32.2 | 39.7 | 66.5 | 41.1 | 22.2 |
| Thr | 29.5 | 26.8 | 24.2 | 32.0 | 20.1 | 23.0 | 23.7 | 26.5 | 37.2 | 43.9 | 36.0 | 50.2 | 60.8 | 50.1 | 26.8 |
| Trp | 3.0 | 4.4 | 3.5 | 3.9 | 3.1 | 3.8 | 3.8 | 3.4 | 3.0 | 2.4 | 1.8 | 2.0 | 2.3 | 2.3 | 3.3 |
| Tyr | 20.5 | 21.5 | 16.2 | 19.2 | 11.2 | 16.0 | 19.4 | 17.5 | 11.4 | 9.2 | 9.3 | 8.9 | 8.2 | 9.1 | 9.4 |
| Val | 33.9 | 34.8 | 31.4 | 45.3 | 25.7 | 28.9 | 32.8 | 31.2 | 42.8 | 34.8 | 34.8 | 38.5 | 41.1 | 43.5 | 47.9 |

*Fold-Change of Polg to WT ( **Figure 3E** )*

| | Liver | | | | | | | Kidney | | | | | | | Heart | | | | | | |
|---|---|---|---|---|---|---|---|---|---|---|---|---|---|---|---|---|---|---|---|---|---|
| Ala | 0.47 | 1.20 | 1.73 | 1.25 | 1.37 | 0.98 | 1.31 | 2.05 | 1.98 | 2.22 | 2.36 | 2.41 | 2.29 | 1.59 | 1.02 | 1.18 | 1.10 | 1.28 | 1.51 | 1.02 | 0.45 |
| Asn | 0.45 | 1.27 | 1.56 | 1.03 | 1.25 | 1.06 | 1.43 | 1.37 | 1.13 | 1.21 | 1.05 | 1.12 | 1.14 | 1.17 | 1.22 | 1.22 | 0.94 | 1.16 | 1.89 | 1.36 | 1.50 |
| Asp | 0.79 | 2.67 | 3.77 | 2.91 | 2.15 | 1.69 | 2.75 | 1.75 | 2.33 | 1.17 | 1.17 | 1.83 | 2.53 | 0.40 | 0.91 | 0.85 | 0.35 | 0.56 | 1.61 | 0.54 | 0.43 |
| Gln | 0.58 | 1.77 | 2.06 | 1.54 | 1.85 | 1.63 | 1.60 | 1.38 | 0.84 | 1.22 | 0.82 | 0.86 | 1.44 | 0.77 | 1.29 | 1.15 | 0.99 | 1.41 | 2.42 | 1.13 | 0.63 |
| Glu | 0.53 | 3.07 | 2.44 | 2.99 | 1.79 | 3.03 | 4.09 | 2.10 | 1.58 | 1.53 | 0.67 | 0.84 | 1.88 | 0.71 | 1.26 | 0.85 | 0.38 | 0.57 | 1.43 | 0.50 | 0.22 |
| Gly | 0.34 | 0.84 | 1.07 | 0.54 | 0.70 | 0.71 | 0.62 | 1.11 | 0.84 | 0.95 | 0.84 | 0.84 | 0.95 | 0.58 | 1.04 | 1.18 | 1.19 | 1.65 | 1.38 | 1.18 | 0.91 |
| His | 0.55 | 1.34 | 1.88 | 1.40 | 1.61 | 1.42 | 1.48 | 1.67 | 1.11 | 1.47 | 1.61 | 1.63 | 1.63 | 1.27 | 1.26 | 1.45 | 1.05 | 1.37 | 1.76 | 1.14 | 1.03 |
| Ile | 0.47 | 1.01 | 1.69 | 0.99 | 0.97 | 1.13 | 1.17 | 1.50 | 1.08 | 1.54 | 1.34 | 1.18 | 1.34 | 1.33 | 1.40 | 1.13 | 1.26 | 1.50 | 1.60 | 1.50 | 1.69 |
| Leu | 0.46 | 0.96 | 1.44 | 0.85 | 0.89 | 0.99 | 1.22 | 1.33 | 0.92 | 1.19 | 1.05 | 0.95 | 1.09 | 1.16 | 1.36 | 1.00 | 1.06 | 1.30 | 1.38 | 1.39 | 1.68 |
| Lys | 0.35 | 0.85 | 1.32 | 0.90 | 1.22 | 0.77 | 1.01 | 0.95 | 0.79 | 0.97 | 0.83 | 0.76 | 0.75 | 0.78 | 1.39 | 1.16 | 1.07 | 1.43 | 1.58 | 0.88 | 0.69 |
| Met | 0.46 | 1.28 | 1.28 | 0.85 | 0.89 | 0.73 | 1.10 | 1.28 | 0.86 | 1.10 | 0.85 | 0.81 | 0.92 | 0.87 | 1.01 | 0.93 | 0.92 | 0.96 | 1.08 | 1.04 | 1.08 |
| Phe | 0.38 | 0.97 | 1.18 | 0.72 | 0.84 | 0.83 | 1.11 | 1.17 | 0.96 | 0.88 | 0.92 | 0.87 | 0.85 | 1.12 | 0.84 | 0.80 | 0.69 | 0.81 | 0.88 | 0.84 | 1.17 |
| Pro | 0.53 | 1.08 | 1.52 | 0.96 | 1.37 | 1.09 | 1.50 | 2.05 | 1.53 | 2.02 | 1.81 | 1.97 | 2.32 | 1.65 | 1.92 | 2.02 | 1.84 | 2.11 | 2.38 | 2.09 | 2.13 |
| Ser | 0.45 | 1.28 | 1.79 | 1.32 | 1.28 | 1.12 | 1.41 | 1.70 | 1.26 | 1.56 | 1.46 | 1.56 | 1.65 | 1.14 | 1.53 | 1.87 | 1.59 | 1.96 | 3.28 | 2.03 | 1.10 |
| Thr | 0.40 | 0.90 | 1.26 | 0.95 | 1.10 | 1.01 | 1.05 | 1.71 | 1.34 | 1.60 | 1.58 | 1.73 | 1.95 | 1.21 | 1.45 | 1.71 | 1.40 | 1.95 | 2.36 | 1.95 | 1.04 |
| Trp | 0.37 | 0.98 | 1.15 | 0.57 | 0.66 | 0.78 | 0.98 | 1.10 | 0.88 | 1.28 | 0.67 | 0.64 | 1.57 | 0.85 | 0.83 | 0.66 | 0.50 | 0.55 | 0.64 | 0.64 | 0.91 |
| Tyr | 0.40 | 0.79 | 1.35 | 0.60 | 0.62 | 0.63 | 0.72 | 0.87 | 0.69 | 0.83 | 0.67 | 0.53 | 0.61 | 0.67 | 0.64 | 0.52 | 0.53 | 0.50 | 0.46 | 0.51 | 0.53 |
| Val | 0.41 | 0.85 | 1.32 | 0.80 | 0.79 | 0.94 | 1.20 | 1.47 | 1.05 | 1.27 | 1.21 | 1.08 | 1.30 | 1.44 | 1.30 | 1.05 | 1.05 | 1.17 | 1.25 | 1.32 | 1.45 |

#### Supplemental Table 4. Tissue nucleotide phosphate abundance and fold-change (Polg/WT) as shown in Figure 4

*Abundance per mg protein, relative to internal standard*

| | | WT | | | | | | | | POLG | | | | | | |
|---|---|---|---|---|---|---|---|---|---|---|---|---|---|---|---|---|
| LIVER | ATP | 0.019 | 0.013 | 0.013 | 0.023 | 0.038 | 0.028 | 0.007 | 0.022 | 0.0014 | 0.0018 | 0.0004 | 0.0007 | 0.0005 | 0.0001 | 0.0008 |
| | ADP | 0.123 | 0.116 | 0.075 | 0.101 | 0.089 | 0.074 | 0.048 | 0.127 | 0.035 | 0.048 | 0.028 | 0.025 | 0.013 | 0.021 | 0.069 |
| | AMP | 0.472 | 0.412 | 0.279 | 0.266 | 0.190 | 0.159 | 0.200 | 0.336 | 0.996 | 0.630 | 0.619 | 0.497 | 0.650 | 0.467 | 0.658 |

*Values normalized to WT (i.e. WT values set to 1)*

| | | WT | | | | | | | | POLG | | | | | | |
|---|---|---|---|---|---|---|---|---|---|---|---|---|---|---|---|---|
| LIVER | ATP | 0.93 | 0.64 | 0.63 | 1.13 | 1.87 | 1.38 | 0.35 | 1.07 | 0.07 | 0.09 | 0.02 | 0.04 | 0.03 | 0.01 | 0.04 |
| | ADP | 1.31 | 1.23 | 0.79 | 1.07 | 0.94 | 0.79 | 0.51 | 1.35 | 0.38 | 0.51 | 0.29 | 0.27 | 0.14 | 0.23 | 0.73 |
| | AMP | 1.63 | 1.43 | 0.96 | 0.92 | 0.66 | 0.55 | 0.69 | 1.16 | 3.44 | 2.18 | 2.14 | 1.72 | 2.25 | 1.61 | 2.27 |

**Supplemental Table 5. Tissue purine catabolism intermediate abundance and fold-change (Polg/WT) as shown in Figure 5H and Supplemental Figure 4I**

*Abundance per mg protein, relative to internal standard*

| LIVER | WT | | | | | | | | POLG | | | | | | |
|---|---|---|---|---|---|---|---|---|---|---|---|---|---|---|---|
| Adenosine | 0.004 | 0.004 | 0.003 | 0.003 | 0.002 | 0.001 | 0.002 | 0.004 | 0.008 | 0.006 | 0.006 | 0.005 | 0.005 | 0.005 | 0.006 |
| Adenine | 0.061 | 0.061 | 0.047 | 0.046 | 0.027 | 0.021 | 0.032 | 0.055 | 0.117 | 0.076 | 0.084 | 0.062 | 0.080 | 0.067 | 0.081 |
| GMP | 0.047 | 0.043 | 0.030 | 0.031 | 0.025 | 0.016 | 0.018 | 0.034 | 0.043 | 0.036 | 0.028 | 0.027 | 0.024 | 0.024 | 0.033 |
| Guanosine | 0.007 | 0.005 | 0.005 | 0.006 | 0.004 | 0.003 | 0.003 | 0.006 | 0.004 | 0.004 | 0.004 | 0.003 | 0.004 | 0.003 | 0.004 |
| IMP | 0.039 | 0.033 | 0.018 | 0.017 | 0.018 | 0.013 | 0.016 | 0.022 | 0.065 | 0.126 | 0.087 | 0.051 | 0.087 | 0.062 | 0.069 |
| Inosine | 0.032 | 0.051 | 0.021 | 0.021 | 0.012 | 0.009 | 0.024 | 0.031 | 0.060 | 0.087 | 0.066 | 0.027 | 0.091 | 0.051 | 0.055 |
| Hypoxanthine | 0.008 | 0.011 | 0.005 | 0.005 | 0.003 | 0.002 | 0.005 | 0.007 | 0.021 | 0.019 | 0.017 | 0.009 | 0.025 | 0.014 | 0.017 |
| Xanthine | 0.043 | 0.059 | 0.031 | 0.029 | 0.017 | 0.013 | 0.031 | 0.045 | 0.116 | 0.109 | 0.101 | 0.046 | 0.138 | 0.072 | 0.091 |

*Values normalized to WT (i.e. WT values set to 1) (* ***Figure 5H*** *)*

| LIVER | WT | | | | | | | | POLG | | | | | | |
|---|---|---|---|---|---|---|---|---|---|---|---|---|---|---|---|
| Adenosine | 1.37 | 1.37 | 1.07 | 1.10 | 0.61 | 0.45 | 0.75 | 1.27 | 2.80 | 1.98 | 1.98 | 1.67 | 1.87 | 1.73 | 2.26 |
| Adenine | 1.39 | 1.40 | 1.07 | 1.04 | 0.62 | 0.49 | 0.72 | 1.26 | 2.69 | 1.75 | 1.92 | 1.42 | 1.84 | 1.54 | 1.86 |
| GMP | 1.53 | 1.39 | 0.97 | 1.02 | 0.83 | 0.54 | 0.59 | 1.12 | 1.40 | 1.17 | 0.92 | 0.89 | 0.77 | 0.78 | 1.09 |
| Guanosine | 1.44 | 1.15 | 1.02 | 1.22 | 0.77 | 0.56 | 0.61 | 1.23 | 0.91 | 0.83 | 0.81 | 0.59 | 0.79 | 0.61 | 0.93 |
| IMP | 1.76 | 1.49 | 0.81 | 0.79 | 0.83 | 0.59 | 0.73 | 1.01 | 2.96 | 5.68 | 3.94 | 2.30 | 3.91 | 2.79 | 3.14 |
| Inosine | 1.28 | 2.02 | 0.85 | 0.82 | 0.49 | 0.34 | 0.97 | 1.22 | 2.41 | 3.45 | 2.64 | 1.08 | 3.64 | 2.03 | 2.21 |
| Hypoxanthine | 1.33 | 1.86 | 0.96 | 0.84 | 0.50 | 0.37 | 0.91 | 1.23 | 3.67 | 3.32 | 3.00 | 1.55 | 4.40 | 2.52 | 3.01 |
| Xanthine | 1.29 | 1.76 | 0.92 | 0.86 | 0.51 | 0.39 | 0.94 | 1.33 | 3.46 | 3.26 | 3.01 | 1.36 | 4.09 | 2.16 | 2.71 |

*Abundance per mg protein, relative to internal standard*

| KIDNEY | WT | | | | | | | | POLG | | | | | | |
|---|---|---|---|---|---|---|---|---|---|---|---|---|---|---|---|
| Adenosine | 0.002 | 0.004 | 0.004 | 0.005 | 0.007 | 0.004 | 0.006 | | 0.006 | 0.005 | 0.006 | 0.006 | 0.005 | 0.006 | 0.003 |
| Adenine | 0.039 | 0.050 | 0.061 | 0.073 | 0.093 | 0.061 | 0.086 | | 0.081 | 0.072 | 0.086 | 0.093 | 0.073 | 0.082 | 0.051 |
| GMP | 0.013 | 0.011 | 0.013 | 0.016 | 0.017 | 0.012 | 0.013 | | 0.017 | 0.016 | 0.016 | 0.014 | 0.018 | 0.020 | 0.018 |
| Guanosine | 0.002 | 0.003 | 0.003 | 0.004 | 0.004 | 0.003 | 0.005 | | 0.004 | 0.004 | 0.004 | 0.005 | 0.003 | 0.004 | 0.003 |
| IMP | 0.043 | 0.032 | 0.030 | 0.048 | 0.033 | 0.030 | 0.040 | | 0.041 | 0.045 | 0.042 | 0.028 | 0.038 | 0.047 | 0.034 |
| Inosine | 0.096 | 0.118 | 0.154 | 0.180 | 0.223 | 0.193 | 0.261 | | 0.158 | 0.118 | 0.155 | 0.089 | 0.118 | 0.116 | 0.068 |
| Hypoxanthine | 0.063 | 0.086 | 0.125 | 0.142 | 0.173 | 0.156 | 0.210 | | 0.146 | 0.095 | 0.147 | 0.064 | 0.088 | 0.095 | 0.058 |
| Xanthine | 0.048 | 0.072 | 0.099 | 0.108 | 0.129 | 0.120 | 0.141 | | 0.144 | 0.099 | 0.164 | 0.102 | 0.127 | 0.118 | 0.083 |

*Values normalized to WT (i.e. WT values set to 1) (* ***Supplemental Figure 4I*** *)*

| KIDNEY | WT | | | | | | | | POLG | | | | | | |
|---|---|---|---|---|---|---|---|---|---|---|---|---|---|---|---|
| Adenosine | 0.52 | 0.78 | 0.93 | 1.12 | 1.46 | 0.88 | 1.32 | | 1.24 | 1.11 | 1.35 | 1.34 | 1.19 | 1.25 | 0.75 |
| Adenine | 0.59 | 0.75 | 0.93 | 1.11 | 1.40 | 0.92 | 1.30 | | 1.22 | 1.09 | 1.30 | 1.41 | 1.11 | 1.24 | 0.77 |
| GMP | 0.96 | 0.80 | 0.94 | 1.17 | 1.26 | 0.87 | 1.00 | | 1.27 | 1.20 | 1.19 | 1.05 | 1.32 | 1.53 | 1.37 |
| Guanosine | 0.47 | 0.91 | 0.77 | 1.13 | 1.34 | 1.00 | 1.38 | | 1.17 | 1.05 | 1.07 | 1.41 | 0.99 | 1.19 | 0.79 |
| IMP | 1.18 | 0.86 | 0.83 | 1.31 | 0.90 | 0.82 | 1.10 | | 1.12 | 1.25 | 1.14 | 0.77 | 1.04 | 1.30 | 0.92 |
| Inosine | 0.55 | 0.67 | 0.88 | 1.03 | 1.27 | 1.10 | 1.49 | | 0.90 | 0.68 | 0.89 | 0.51 | 0.68 | 0.67 | 0.39 |
| Hypoxanthine | 0.46 | 0.63 | 0.91 | 1.04 | 1.27 | 1.14 | 1.54 | | 1.07 | 0.69 | 1.08 | 0.47 | 0.65 | 0.70 | 0.42 |
| Xanthine | 0.47 | 0.70 | 0.96 | 1.06 | 1.26 | 1.17 | 1.38 | | 1.40 | 0.96 | 1.60 | 1.00 | 1.24 | 1.15 | 0.81 |

#### Supplemental Table 6. m/z and retention time for LC/MS (iHILIC) analysis

| Metabolite | Formula | m/z (negative mode) | Retention time (min) |
|---|---|---|---|
| 3-Hydroxybutyric acid | $C_4H_8O_3$ | 103.0401 | 2.9 |
| Acetoacetate | $C_4H_6O_3$ | 101.0244 | 2.6 |
| Adenine | $C_5H_5N_5$ | 134.0472 | 3.3 |
| Adenosine | $C_{10}H_{13}N_5O_4$ | 266.0895 | 3.3 |
| ADP | $C_{10}H_{15}N_5O_{10}P_2$ | 426.0221 | 9.9 |
| AMP | $C_{10}H_{14}N_5O_7P$ | 346.0558 | 8.3 |
| ATP | $C_{10}H_{16}N_5O_{13}P_3$ | 505.9885 | 10.9 |
| GMP | $C_{10}H_{14}N_5O_8P$ | 362.0507 | 10.4 |
| Guanosine | $C_{10}H_{13}N_5O_5$ | 282.0844 | 6.1 |
| Hypoxanthine | $C_5H_4N_4O$ | 135.0312 | 3.8 |
| IMP | $C_{10}H_{13}N_4O_8P$ | 347.0398 | 9.6 |
| Inosine | $C_{10}H_{12}N_4O_5$ | 267.035 | 4.4 |
| Uric Acid | $C_5H_4N_4O_3$ | 167.0211 | 6.1 |
| Xanthine | $C_5H_4N_4O_2$ | 151.0261 | 4.6 |

#### Supplemental Table 7. Ion transitions for LC/MS (Accucore C30) analysis

| Lipid Species | Precursor Ion | Product Ion |
|---|---|---|
| ***Triacylglycerols (TAG)*** | | |
| TAG(48:0) | 824.8 | 551.5 |
| TAG(48:3) | 818.7 | 547.5 |
| TAG(50:2) | 848.8 | 549.5 |
| TAG(50:3) | 846.8 | 549.5 |
| TAG(52:1) | 878.8 | 605.6 |
| TAG(52:2) | 876.8 | 577.5 |
| ***Diacylglycerols (DAG)*** | | |
| DAG(16:0/18:1) | 612.6 | 313.3 |
| DAG(16:0/18:2) | 610.5 | 313.3 |
| DAG(16:0/18:3) | 608.5 | 313.3 |
| DAG(16:0/22:6) | 658.5 | 313.3 |
| DAG(16:1/18:1) | 610.5 | 311.0 |
| ***Phosphatidylethanolamine (PE)*** | | |
| PE(34:2) | 716.5 | 575.5 |
| PE(36:3) | 742.5 | 601.5 |
| PE(36:2) | 744.6 | 603.5 |
| PE(38:7) | 762.5 | 621.5 |
| PE(38:6) | 764.5 | 623.5 |
| PE(38:3) | 770.6 | 629.6 |
| PE(40:7) | 790.5 | 649.5 |
| ***Phosphatidylcholine (PC)*** | | |
| PC(33:1) | 746.4 | 184.1 |
| PC(34:0) | 762.6 | 184.1 |
| PC(34:1) | 760.6 | 184.1 |
| PC(36:0) | 790.6 | 184.1 |
| PC(36:1) | 788.6 | 184.1 |
| PC(36:2) | 786.6 | 184.1 |
| PC(38:1) | 816.6 | 184.1 |
| PC(38:6) | 806.6 | 184.1 |
